# Species-specific gene duplication in *Arabidopsis thaliana* evolved novel phenotypic effects on morphological traits under strong positive selection

**DOI:** 10.1101/2021.04.05.438504

**Authors:** Yuan Huang, Jiahui Chen, Chuan Dong, Dylan Sosa, Shengqian Xia, Yidan Ouyang, Chuanzhu Fan, Dezhu Li, Emily Mortola, Manyuan Long, Joy Bergelson

**Author notes:** co-first authors for equal contributions (Yuan Huang, Jiahui Chen).

## Abstract

Gene duplication is increasingly recognized as an important mechanism for the origination of new genes, as revealed by comparative genomic analysis. However, the ways in which new duplicate genes contribute to phenotypic evolution remain largely unknown, especially in plants, owing to a lack of experimental and phenotypic data. In this study, we identified the new gene *Exov,* derived from a partial gene region duplication of its parental gene *Exov-L*, which is a member of an exonuclease family, into a different chromosome in *Arabidopsis thaliana*. We experimentally investigated the phenotypic effects of *Exov* and *Exov-L* in an attempt to understand how the new gene diverged from the parental copy and contributes to phenotypic evolution. Evolutionary analysis demonstrated that *Exov* is a species-specific gene that originated within the last 3.5 million years and shows strong signals of positive selection. Unexpectedly, RNAseq analyses reveal that the new gene, despite its young age, has acquired a large number of novel direct and indirect interactions in which the parental gene does not engage. This is consistent with a high, selection-driven substitution rate in the protein sequence encoded by *Exov* in contrast to the slowly evolving *Exov-L*, suggesting an important role for *Exov* in phenotypic evolution. We analyzed phenotypic effects of *exov* and *exov-l* single T-DNA-insertion mutants;double *exov, exov-l* T-DNA insertion mutants; and CRISPR/Cas9-mediated *exov^crp^* and *exov-l^crp^* knockouts on seven morphological traits in both the new and parental genes. We detected significant segregation of morphological changes for all seven traits when assessed in terms of single mutants, as well as morphological changes for seven traits associated with segregation of double *exov, exov-l* mutants. Substantial divergence of phenotypic effects between new and parental genes was revealed by principal component analyses, suggesting neofunctionalization in the new gene. These results reveal a young gene that plays critical roles in biological processes that underlie morphological and developmental evolution in *Arabidopsis thaliana*.

## Introduction

The origination of novel genes is an important process contributing to the evolution of organisms, as new genes have the potential to become genetic sources of evolutionary innovation (Long et al., 2013; Chen et al., 2013). Recent studies have identified lineage-specific and species-specific genes with important effects on diverse phenotypes, including development, sexual reproduction, brain functions, and behavior (Park et al. 2008; Ding et al., 2010; Chen et al., 2010; Zhang et al., 2011; VanKuren et al., 2018; Lee et al., 2019). However, all of these studies have focused on metazoans, such as invertebrates, including fruit flies, and mammals. Consequently, little is known about the extent to which new gene evolution has coordinated phenotypic changes in plants, leading to a gap in our understanding of molecular and phenotypic evolution.

New genes typically arise through the duplication of existing genes at the DNA level, although a number of other mechanisms have been reported (Long et al., 2003 and 2013). These new genes may maintain functions similar to the parental gene or may undergo a process of diversification until a completely novel function has evolved. Recently born genes, especially those appearing within the past few million years, provide excellent opportunities to study gene formation and associated phenotypic evolution, since all or most incipient changes are clearly recorded and preserved in extant organisms (Chen et al., 2013; Long et al., 2013; Zhang et al, 2019). As such, one can relate evolutionary changes in the genes to corresponding phenotypic expression.

In this study, we examine *Exov* (AT3G57110), a species-specific *Arabidopsis* gene that originated in the *A. thaliana* lineage 3.5 million years ago (MYA) through the duplication of the *Exov-L* (AT5G60370) gene in chromosome 5, which was partially copied into a new locus in chromosome 3. We perform a comprehensive investigation of its phenotypic effects within an evolutionary context and analyze the selective forces acting upon it. Our results reveal the unexpectedly large effects of this new gene on the evolution of morphological traits, demonstrating that new genes can drive rapid phenotypic evolution *in planta*.

## Materials and methods

### Plant materials and growth conditions

*Arabidopsis* seeds were surface sterilized with 50% commercial bleach for 5 min and then rinsed five times with sterile water. Following 2-3 days of stratification at 4 °C, *Arabidopsis* plants, including several related species (*A. thaliana*, *A. lyrata* subsp. *lyrata*, *A. lyrata* subsp*. petraea*, and *A. halleri*), were grown under a long-day condition (16 hours light / 8 hours dark at 22 °C) in the University of Chicago greenhouse for 5-6 weeks.

The *Arabidopsis* T-DNA insertion lines for *Exov*, including *exov-1* (Salk-103969), *exov-2* (Salk-036494) and *exov-3* (Salk-064431), and for *Exov-L*, *exov-l* (Salk-101821) were ordered from the *Arabidopsis* Biological Resource center at Ohio State University (http://www.arabidopsis.org/). These T-DNA mutants were identified as single mutants by adaptor-nested PCR (Huang et al. 2007). The locations of the T-DNA insertions in the sequence-indexed *Arabidopsis* mutant seeds were confirmed by PCR amplification using the T-DNA border primers (LBb1.3) and gene-specific primer (LPs (Left Primers), RPs (Right Primers) for both new gene and parental gene). Plants with a homozygous T-DNA insertion were identified by screening self-fertilized progeny from the mutants using PCR amplification. Homozygous lines were identified by negative LP-RP amplification and positive LBb1.3-RP amplification. The exact DNA insertion positions were verified by sequencing the LBb1.3-RP PCR products. The LBb1.3 for all SALK lines is 5’-ATTTTGCCGATTTCGGAAC-3’. The LPs and RPs are 5’-GAAAAATTAGTCAGCAGTCGGG-3’ and 5’-CAATCATGGTGAGATTCCAAAG-3’ for SALK_103969, 5’-TGGAAGACGAAGTGGTAGGTG-3’ and 5’-CGTCGTCGCTACTATTCGATC-3’ for SALK_064431, 5’-CTCTCACAATTAGCCGCTGTC-3’ and 5’-TTGGAGAAATCATGGAGATCG-3’ for SALK_036494, and 5’-TAGCAAATTGGCAATACCGAC-3’ and 5’-AGCTGTTGAATTCCATTGCTG-3’ for SALK_101821. Double mutant lines were created by crossing Salk_101821 with Salk_103969, Salk_036494, and Salk_064431, respectively. Homozygous double *exov, exov-l* mutant plants were identified by using 4xPCR reactions, showing negative LP-RP amplification and positive LBb1-RP amplification of both genotypes. T2 homozygous plants for T-DNA insertion were used to evaluate phenotypic changes through a comparison to wild type individuals (Col-0). The consistent phenotypic effects among the T-DNA lines for single and double mutants and the knockout lines created by CRISPR/Cas9 (see the section below) further suggest that both T-DNA and CRISPR/Cas9 lines are lacking substantial background mutations, including additional insertions of the T-DNA.

### Generation of the *exov^crp^* and *exov-l^crp^* mutants of the new gene and parental gene using CRISPR/Cas9

CRISPR/Cas9 vector pCAMBIA1300 was used to create knock-out (KO) mutations in *Exov* and *Exov-L* (Yan et al., 2015). Complete sequence information for the vector, the map, and the annotated vector sequences are shown (Supplementary Figure S1). The CRISPR/Cas9 constructs were transformed into *A. thaliana* wild-type Columbia-0 (Col-0) through floral dipping. T1 plants were selected either by red fluorescence or on 16 mg L21 hygromycin. Genomic DNA samples extracted from leaf tissues of 2-week-old T1 plants were used as templates for PCR. To screen mutations at the *Exov* and *Exov-L* targets, we used the primer pairs 57110R (5’-TTCCTATGATATGACTGTGATATA-3’) and 57110F (5’-GCATAGACATGAAAAAAGAAGAA-3’), and 60370R(5’-CACATGTTGGTTCCGAATAAAACA-3’) and 60370F(5’-GCTTTATTGACTTTTCTCCTGCCA-3’), respectively, to amplify the target-containing fragments. We focused our PCR screening for mutants on plants that we identified as Cas9-free. All of these homozygous T2 transgenic lines (*exov^crp^, exov-l^crp^*) were identified by directly sequencing PCR products and the whole genome sequencing as below.

### Identification of mutation sites of T-DNA lines and CRISPR lines

Whole genome sequencing data were generated to identify mutation sites using Illumina Sequencing with the genome coverage greater that 99% and read depth higher than 50 (Supplementary Table S1). For T-DNA insertion mutants, raw reads were *de novo* assembled by SOAPdenovo2 (Luo, et al., 2015) and chimeric sequences bridging T-DNA plasmid and *Arabidopsis* genome were identified by BLAT (Kent, 2002). For CRISPR mutants, raw reads were first mapped to TAIR10 (Berardini, et al., 2015) by BWA (Li and Durbin, 2010) and VCF files were generated by GATK (Van der Auwera, et al., 2013) and corrected with 1001 genomes (Genomes Consortium. Electronic address and Genomes, 2016). After that, on-target and off-target sites were predicted by CRISPR-P 2.0 (Liu, et al., 2017) online and mutation sites were retrieved in 100 bp region centering on the expected target loci. Furthermore, mapping T-DNA insertion sites were conducted by fusion primers and nested integrated PCR (Wang, et al., 2011). The potential on-target and off-target sites were mapped on the genome sequence. Target products of through FPNI-PCR including T-DNA insertion flanking sequence and target genome sequence were sequenced and blasted in whole genome of *A. thaliana* to confirm the insertion positions.

**Table 1.**
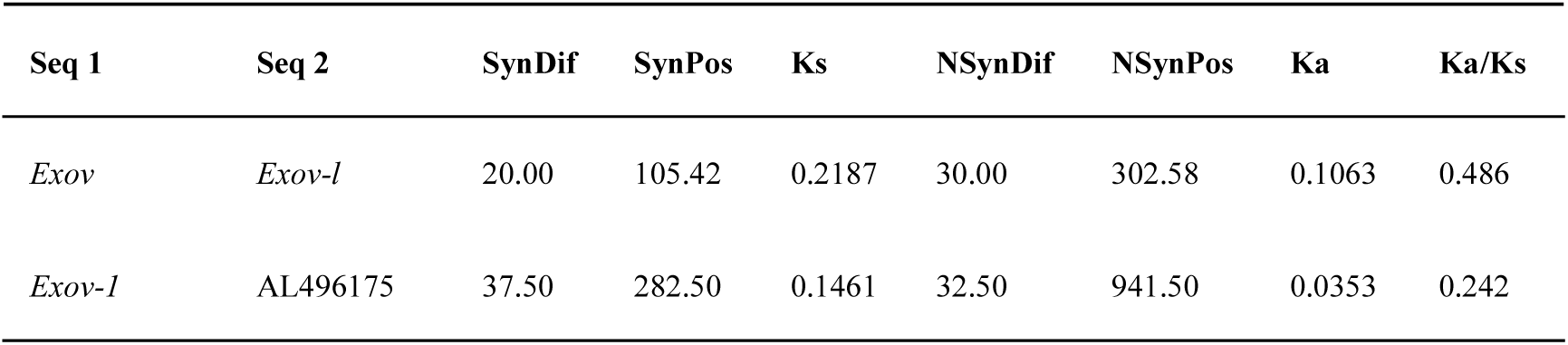
Ka/Ks ratio of new gene and parental genes.

We identified single T-DNA insertion target new gene and parental sequences based on whole genome sequencing data (Supplementary Table S1). The insertion sites were verified by mapping. Excepted target positions, no insertion were mapped to other position of chromosomes. The mapped flanking sequences indicate the chromosomal insertion positions of the corresponding T-DNA lines of new gene and parental gene, 21134854 to 21135628 bp on chromosome 3 and 24283931 to 24291840 bp on chromosome 5 respectively (Supplementary file 1 and Supplementary Table S1). The consistence of genome data and mapping T-DNA sites proved single TDNA insertion mutant lines of *Exov* and *Exov-L* genes.

For CRISPR lines, we used the whole genomes of 1001 accessions as background to filter the off-target sites. No off-targets were detected in both *exov* and *exov-l* CRISPR lines. The on-target was confirmed in the *exov* CRISPR KO line by insertion T while deletion G was detected in the *exov-l* CRISPR KO line (Supplementary file 2 and Supplementary Table S1).

### DNA sequencing, qRT-PCR, and transcriptome analysis

#### DNA sequencing

The new gene *Exov* and old gene *Exov-L* were PCR-amplified from genomic DNA in four separate reactions using the primer pairs in Supplementary Table S2 and Supplementary Figure S1. Following PCR, the amplified products were sequenced from both strands using the primer pairs, BidDye chemistry, and a 3730 automated sequencer (Applied Biosystems).

**Table 2:** The McDonald-Kreitman test of natural selection.

| | Substitutions | | Polymorphisms | | $\alpha$ | Fisher exact Probability |
| --- | --- | --- | --- | --- | --- | --- |
|  | Dn | Ds | Pn | Ps |  |  |
| A. thaliana Exov-L | 3 | 4 | 7 | 4 | -1.33 | 0.6534 |
| A. thalian Exov | 22 | 12 | 3 | 9 | 0.82 | 0.0229 |

Note: The subscripts n and s indicate nonsynonymous and synonymous changes, respectively. $\alpha$ for *Exov* is the proportion of substitution driven by positive selection; $\alpha$ for *Exov-L* may be the sampling error or segregation of deleterious mutations (Smith and Eyre-Walker, 2002).

#### Quantitative RT-PCR

To compare the expression levels of the new and parental genes in different tissues of our set of mutants and the wild type plants, leaves, flowers, young siliques, and stems were collected for RNA extraction. Total RNA was extracted using the Eastep® Super Total RNA Extraction Kit (Promega) and reverse transcribed using the Reverse Transcription System (Promega) according to the manufacturer’s protocol. Quantitative real-time PCR was performed with the ABI7500 real-time PCR system using TransStart® Top Green qPCR SuperMix (TransGen, Beijing, China). The relative gene expression level was calculated by normalizing against the internal control ACTIN8. Three biological replicates were carried out for each sample. All primers used for RT-qPCR are listed in Supplementary Table S2.

#### RNA-seq transcriptome analysis

To compare the expression patterns and biological processes of the new and parental genes, the whole plants of wild-type and mutant genotypes growing under a long-day condition (16 hours light / 8 hours dark at 22 °C) in KIB greenhouse for 6-8 weeks, including leaf, flower, stem and all other tissues, were sampled in liquid nitrogen upon collection for RNA sequencing. Total RNA from three biological replicates of wild-type *A. thaliana*, T-DNA mutants (*exov* and *exov-l*), and CRISPR/Cas9 mutants (*exov^crp^, exov-l^crp^*) were extracted with Trizol reagents. mRNAs were purified using an Oligotex mRNA Mini Kit (QIAGEN). Next, cDNA libraries were prepared using the mRNA-Seq Sample Preparation Kit™ (Illumina) following a non-strand-specific protocol. Briefly, mRNAs were fragmented by exposure to divalent cations at 94°C, and fragmented mRNAs were converted into double-stranded cDNA. Then, cDNA ends were polished with the 39-hydroxls extended with A bases and ligated to Illumina-specific adapter-primers. The resulting DNA was amplified by 15 cycles of PCR followed by purification using the Qiagen™ PCR Purification Kit to obtain the final library for sequencing on the Illumina HiSeq2000 platform. The DNA yield and fragment insert size distribution of sequencing libraries were determined on the Agilent Bioanalyzer.Tophat version 2.0.12 was used to map reads to the *A. thaliana* genome version TAIR10. Next, cuffdiff version 2.2.1 was used to find differentially expressed genes between samples (Trapnell et al., 2012), which were then applied to GOrilla for gene ontology enrichment analysis (Eden et al., 2009). To check the knockdown efficiency of mutants, we counted uniquely mapped reads as the expression levels of the parental gene and new gene using HTSeq with “union” mode (Anders, Pyl, and Huber, 2014).

### Measurement and analysis of phenotypes

#### Measurement

A set of 7 morphological traits--the length of the rosette major axis, length of the rosette minor axis, leaf number, number of stem branches on main bolts, number of side bolts, time until the first open flower, and height of the main bolt at landmark growth stages--were collected (Figure 1). About 400 individuals of each genotype, including wild-type (WT); single T-DNA insertion lines and double *exov, exov-l* mutant lines; and 100 individuals of each of CRISPR/Cas9 lines and WT --were grown in soil-flats for observation of phenotypes in the greenhouses at the University of Chicago (for T-DNA lines and their control) and Kunming Institute of Botany (for CRISPR-Cas9 lines and their control). For the calculation of rosette area and the number of rosette leaves, soil-grown plants at stage 1.04 (15 days) were measured with a vernier caliper, and leaves were counted. The time at which the first flower opened was collected between stage 3.00 (23 days) and stage 6.90 (50 days). In addition, the height of soil-grown plants at stage 6.10 (36 days) was measured with a vernier caliper and ruler, and the number of bolting shoots was counted (Supplementary Table S3). The analysis of *Arabidopsis* growth and development presented here provides a framework for identifying and interpreting phenotypic differences in plants resulting from genetic variation caused by mutations (Boyes et al., 2001).

**Figure 1.**
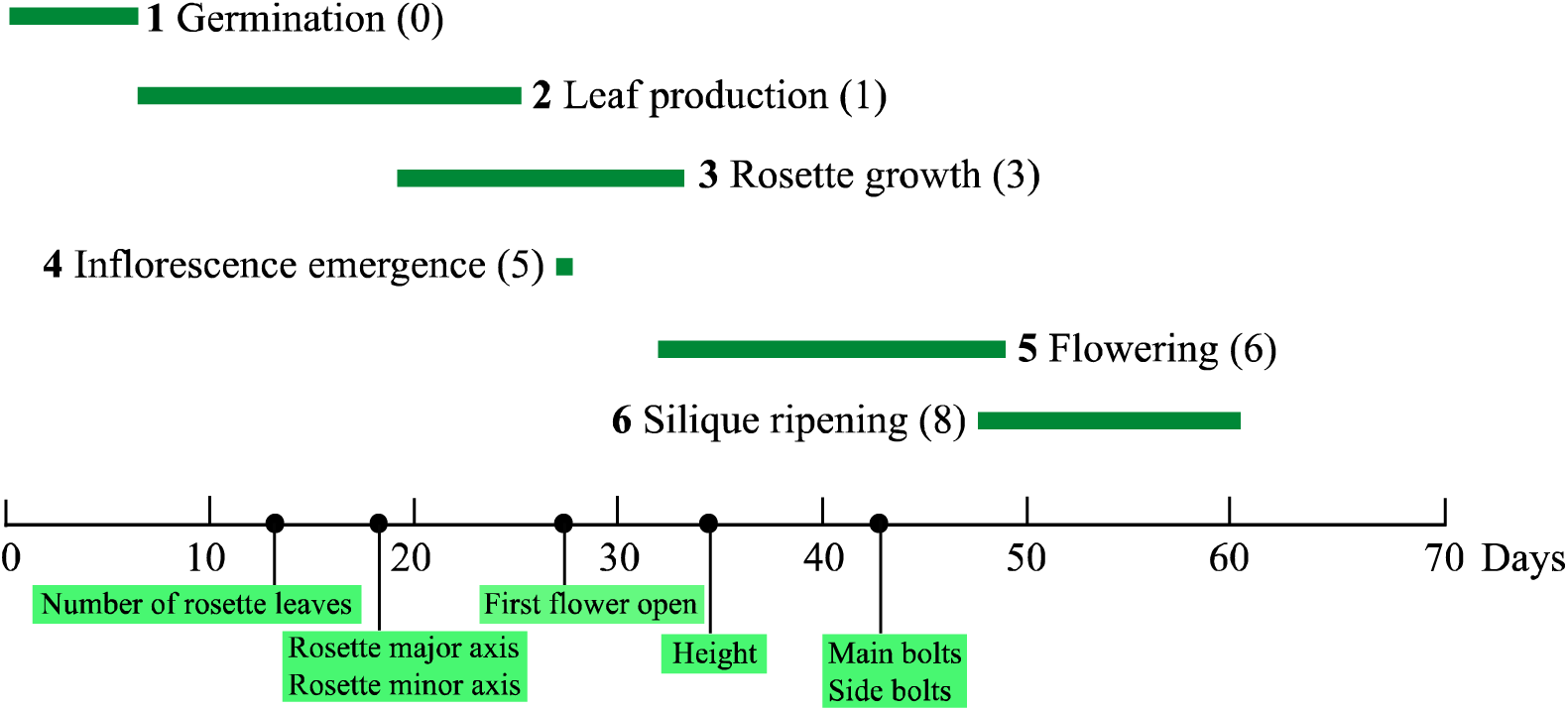
**The distribution of observed traits in the growth of *A. thaliana*** as adapted from Boyes et al. (2001). The black dots on the X axis represent the timing of phenotypic measurements.

#### Estimating the phenotypic effects distribution of mutants

To estimate the distribution of the phenotypic effects of mutations on the trait, we analyzed the phenotypes associated with the new and parental genes. For analytical tractability, we adopted the models of Turelli (1984), Sawyer et al. (2003), and Jones et al. (2007), assuming that the phenotypic effects of mutant and wild type alleles on a trait follow a Gaussian distribution with mean μ and standard deviation σ (Jones, Arnold, and Bürger, 2007; Sawyer et al., 2003; Turelli, 1984).

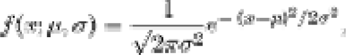

The distribution of mutational effects on each trait was inferred from the changes in the trait value among the mutants and the wild-type. Phenotypic differences in each of our seven traits between wild-type and mutant lines were assessed for both the T-DNA insertions and CRISPR/Cas9 mutations. Although the formal distribution of the mutational effects for any given trait is unknown, the change in the distribution of mutational effects on a trait can be inferred by the deviation from the distribution of trait value in the wild-type, such as a shift in the frequency peak. The theoretical curve for each of the observed trait distributions was determined as the best fitted curve of a Gaussian distribution using R (v4.0.4).

#### Principal component analysis

To characterize the growth of *Arabidopsis*, we performed principal component analysis on the seven morphological traits, using phenotypes measured on the T-DNA insertion lines and double mutant lines, CRISPR-Cas9 lines, and the wild-type plants. Because the T-DNA insertion lines and the CRISPR KO lines were grown in two separate experiments, they were considered separately. PCA was performed using the R program (predict and princomp in v4.0.4).

A technical issue is that data involved are large numbers of data points (e.g. *exov* has more than one thousand individuals of three mutants), which would make it hard to visualize the phenotypic differences of various mutants. We developed a simple geometric method to calculate the phenotypic distance between the new gene *Exov* and the parental gene *Exov-L*, which are defined by the pairs of average principle components of each genotype. The first two principle components, PC1 and PC2, which are highly representative of the variation of morphological traits we investigated (∼80%), were used to form a two dimensional space. If we use Gi to denote a gene i in a pair of average PC values, PC1(Gi) and PC2(Gi), that are given by PCA for a population, then the difference in phenotypic evolution (PED) between the two genes can be mathematically described by using a geometric distance between gene mutants i and j measured by the following formula:

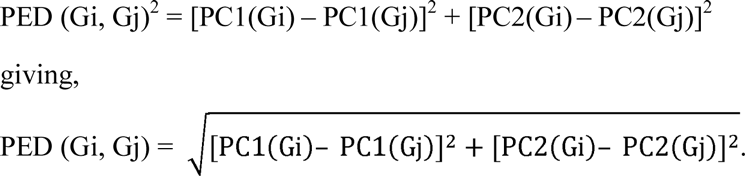

Thus, the PED describes a distance of phenotypic evolution that occurs in the two genes in terms of eigenvectors of the measured morphological traits. We will show that this geometrical description is helpful when we compare the contribution of new gene and parental gene in a large dataset of measured morphological traits.

### Evolutionary Analysis

#### Sequence comparison of Exov and Exov-L

Protein sequences of EXOV and EXOV-L were downloaded from TAIR (http://www.arabidopsis.org/) and aligned by Geneious (Drummond et al., 2011). Orthologous coding sequences of *Exov-L* were downloaded from phytozome v9.1 (http://www.phytozome.net/). Alignments of coding sequences mentioned below were performed by MEGA 3.2, considering the coding structures. For synteny analysis, genetic location information on *Exov* and *Exov-L* were obtained from the TAIR website (http://www.arabidopsis.org/). The syntenic relationship among *Exov, Exov-L,* and the orthologous genes Aly496175 (*Arabidopsis lyrata*), Cru10026530 (*Capsella rubella*), Tha10013696m (name species), Bra020254 (*Brassica rapa*), and Osa05g03200 (name species) are displayed by Phytozome (http://www.phytozome.net/). For phylogenetic analysis, gene sequences of *Exov* and *Exov-L* were aligned with *Capsella, Eutrema, Brassica,* and *Oryza* using Geneious and manually adjusted. A phylogenetic tree was created according to the maximum likelihood method using the MEGA 5.2.2 program (Tamura et al., 2011).

#### Population genetics of *Exov* and *Exov-L*

Genotypes of worldwide accessions were obtained from the *Arabidopsis* 1001 Genomes Project (Supplementary Table S4). This dataset was used for population genetic analysis, including the 851 accessions that remained after filtering accessions that were misidentified and discarding sequences of poor quality or with sequencing errors (Anastasio et al., 2011). Basic population genetic analyses were implemented in the DnaSP5 program. Sequence diversity was calculated using nucleotide diversity (π) and the population mutation parameter of Watterson’s estimator. Synonymous substitution rates (Ks) and non-synonymous substitution rates (Ka) were calculated using DnaSP5.10.1 (Rozas et al., 2003).

#### Substitution analysis and testing selection

Following strict parsimony, we identified all the substitutions that contribute to the divergence of *Exov* and *Exov-L* and assigned them to one of the two gene lineages following the duplication event. We conducted these analyses from a multiple gene sequence alignment, based on the states of the orthologues in outgroup species, defined by a phylogeny {[(*A. thaliana,* (*A. lyrate, A helleri*)), (*C. rubella, C. sativa*)], (*B. rapa and E. salsugineum*)} (genus names: *C., Cannabis*; *B., Brassica; E., Eutrema*). Meanwhile, all sites revealing substitutions on *Exov-L* before the duplication event were also counted. These sites were compared to the polymorphism tables from the 851 *A. thaliana* accessions, which produced 709 *Exov* alleles and 455 *Exov-L* alleles. While most substitutions are present in 100% of the accessions, a few are present in ∼99% of alleles, with no ancestral alleles detected in the population. Tests of deviation from neutrality were conducted by comparing the observed substitutions with the polymorphisms at synonymous and nonsynonymous sites to test the distinctive prediction of neutral theory that the rates of mutation and evolution are equal, following a pipeline we designed for the algorithm (Supplementary Figure S2). In particular, the McDonald-Kreitman test (McDonald and Kreitman, 1991; Smith and Eyre-Walker, 2002) was performed to detect positive selection acting on *Exov* since its origination from the parental gene *Exov-L*.

## Results

### Evolutionary analysis of the new gene *Exov* and the parental gene *Exov-L*

We first describe the history of gene evolution in which the new gene *Exov* was duplicated from the parental copy *Exov-L*, involving the movement from chromosome 5 to chromosome 3 (their sequences and related molecular features are summarized in Supplementary Figure S1). Given the observed gene evolution, we explored the role of positive selection on the new gene locus.

#### The species-specific duplication between chromosome 5 and chromosome 3 gave rise to a new duplicate gene Exov

Analysis of synteny indicates that the parental gene *Exov-L* has orthologs in all 5 related species that we investigated: *A. thaliana*, *A. lyrata*, *C. rubella*, *B. rapa*, and *T. halophile*. Previous phylogenetic analyses estimated that *A. thaliana* split from *A. lyrata* ∼ 5 MYA (Beilstein et al., 2010), from *B. rapa* ∼ 13-17 MYA (Town et al., 2006; Yang et al., 1999), and from *C. rubella* ∼ 10-14 MYA (Koch and Kiefer, 2005). The new gene *Exov* in chromosome 3, which is a duplicate of a portion of the parental gene (Figure 2) in chromosome 5, is present only in the genome of *A. thaliana.* This species-specific copy, *Exov*, was detected in all *A thaliana* accessions used in the population structural analyses of the 1001 Genomes Project, including the genomes of Columbia (Col-0) and Landsburg (Ler-0). These observations suggest that the new gene *Exov* is species-specific and has been fixed in *A. thaliana* since emerging after the recent split between *A. thaliana* and *A. lyrata*.

**Figure 2.**
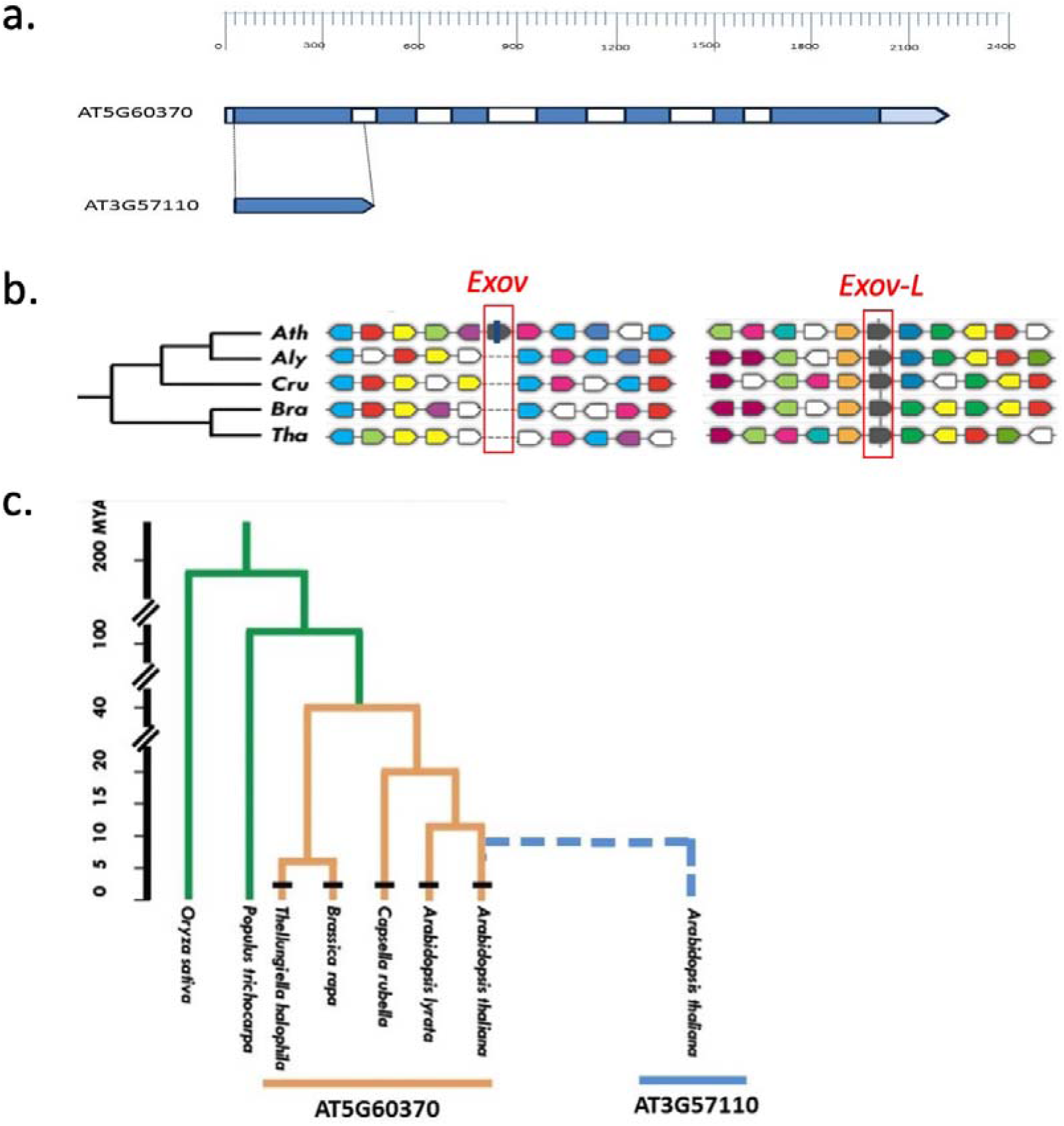
Evolution of *Exov* (AT3G57110) duplicated from *Exov-L* (AT5G60370) inferred from gene structure and syntenic analysis. **a.** Duplication mode and gene structure of new gene and parental gene. Blue boxes, exons; white, introns; gray, untranslated regions (UTRs). **b**. Syntenic analysis of the new gene *Exov* (AT3G57110) and parental gene *Exov-L* (AT5G60370) based on the phylogenic tree. Ath: *A. thaliana*; Aly: *A. lyrata*; Cru: *C. rubella*; Bra: *B. rapa*; Tha: *Thellungiella halophila*. The red blocks highlight the orthologous regions of *Exov* and *Exov-L* in the other 4 related species, showing no orthologous copies for *Exov* and 4 orthologous copies for Aly (*496275*), Cru (*Carubv10026530m*), Bra (*Bra020254*) and Tha (*Thhalv10013696*). Inspection of 10 genes that flank *Exov* and *Exov-l* (the grey arrow blocks with bars) indicates orthologous syntenous arrangement of these genes in support of the orthologous comparison in the highlighted genomic regions of *Exov* and *Exov-L* in the relatives of Ath. The arrows show the orientation of the genes. The colors represent homologous relationships and a color represents a distinct homologous gene. **c.** The phylogeny and divergence time between *A. thaliana* and its relatives and the species distribution of new gene *Exov* (AT3G57110) and *Exov-L* (AT5G60370).

#### Detecting an asymmetrically high rate of substitution in *Exov* in contrast to slow substitution in *Exov-L*

We performed a sliding window analysis of the Ka/Ks ratio between *Exov* and the duplicated portion of *Exov-L* within *A. thaliana*. The Ka/Ks ratio was higher than 1 in the first 100 bp, suggesting that this region is under positive selection. However, in the region between 120-400 bp, the Ka/Ks ratios between *Exov* and *Exov-L* were <0.5, suggesting evolutionary constraint on the protein sequence in this region (Table 1, Figure 3). Notably, the Ka value measuring divergence between *Exov* and *Exov-L* is remarkably high for a duplicated region dating less than 5 million years (0.1063). Indeed, this rate is 3.01 times the Ka value (0.0353) between the *Exov-L* orthologues in *A. thaliana* and *A. lyrata* that diverged earlier than the duplication time of *Exov*. Taking *A. lyrata* and other more distant species, e.g. *C. rubella* and *B. rapa,* as outgroup species in a parsimony analysis, we detected an asymmetrical distribution of substitutions accumulating on *Exov* and *Exov-L* since the duplication event: 22 nonsynonymous substitutions on *Exov* and only 3 nonsynonymous substitutions on *Exov-L* (Table 2, Materials and Methods); values that differ significantly from a null hypothesis of neutrality that predicts equal substitution between the two duplicates (L2 = 14.44, df=1, p = 0.0001).

**Figure 3.**
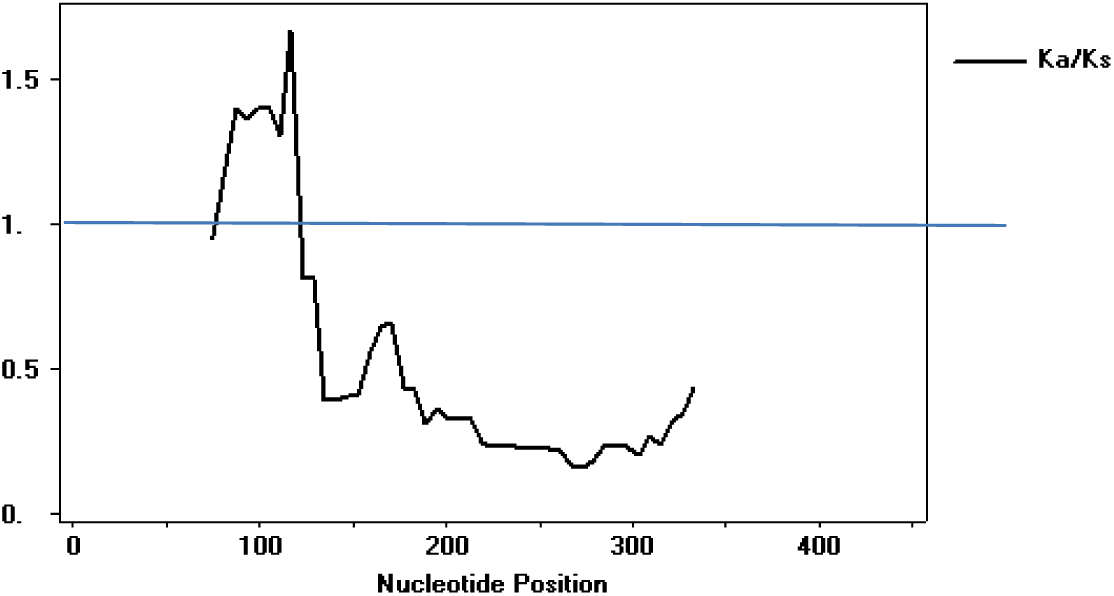
Ka/Ks sliding window analysis. (Window length: 150 bp. Step size: 6 bp.)

The unexpectedly high rate of protein evolution in *Exov* implicates positive selection acting on *Exov*. We took two approaches to test for putative positive selection: a population genetic test of selective sweeps and additional substitution analysis to compare with the population genetic prediction of neutrality. However, before pursuing these approaches, it is necessary to understand the population structures of *A. thaliana* because demographic processes have the potential to impact the population genetic inferences and substitution analyses. Previous analyses (Nordborg et al, 2005; Horton et al, 2003) detected significant population structures using then-large datasets in *A. thaliana*, revealing the need to consider demographic factors when testing selective forces. We used the significantly expanded sequence information in the 1001 genomes project (the 1001 Genomes Consortium, 2016) to update previous population structure analyses for their incorporation in our population genetic analyses.

First, to infer population structure and assign accessions to populations, we used ADMIXTURE1.23 (Alexander et al., 2009), which adopts the likelihood model embedded in STRUCTURE (Raj et al., 2014). To cluster all accessions on the basis of geographic distribution (Supplementary Table S4), we analyzed the data by successively increasing K from 2 to 8 (Supplementary Figure S3a) using the ADMIXTURE likelihood algorithm. The cross-validation error was smallest when K was set equal to 8 (Supplementary Figure 3b), revealing clear global population structure among these 8 subgroups (Supplementary Figure 3c). The population structure was consistent with earlier analyses (Nordborg, 2005; Horton, 2012) that detected population clustering, but with most polymorphisms shared species wide.

This, and previous observations of global population structure across the *A. thaliana* genome (Nordborg et al, 2005; Wright and Gaut, 2005), reveal potential demographic processes that render tests of positive selection too liberal if a comparison is made to a theoretical distribution, which could cause a deviation from expected values for the Tajima D test, the Fay-Wu test, the Fu-Li tests (Fu and Li, 1993; Tajima, 1989; Fay and Wu, 2000b), even in the absence of positive selectioin. We therefore computed the empirical distributions of these statistic tests across the whole genome (Supplementary Table S5; Supplementary Figure S4) using the worldwide accessions (the 1001 Genomes, Supplementary Table 4). Compared to these empirical distributions, we failed to find significance for any of the above population genetic statistics calculated for the *Exov* and *Exov-L* genes (Supplementary Figure S4), suggesting that neither *Evov* nor *Evov-L* has undergone a selective sweep.

We next used the McDonald-Kreitman test (McDonald and Kreitman, 1991) to test for positive selection on the substitutions of *Exov*. Again, such a test would be too liberal due to increased deleterious replacement polymorphisms in local and small populations. In this test, polymorphism within *Exov* in *A. thaliana* was compared to sequence divergence between *Exov* in *A. thaliana* and two outgroup species, *A. lyrata* and *C. rubella.* We also performed the same test for *Exov-L*, comparing polymorphism with species to divergence between species.

We furthermore assigned divergence between *Exov* and *Exov-L* to each lineage since the duplication event and measured the time since the duplication by counting the number of shared synonymous substitutions in *Exov* and *Exov-L* that occurred between the speciation of *A. thaliana* and the duplication of *Exov*. Two of 6 *Exov-L-specific* synonymous substitutions were shared with *Evox* (those at sites 204 and 216), suggesting that *Exov* was duplicated soon after the speciation of *A. thaliana.* We estimated that the duplication occurred 3.5 million years ago (mya), roughly one third of the time since emergence of the *Arabidopsis* lineage 5 mya (Yogeeswaran et al, 2005).

For the McDonald-Kreitman test, we counted polymorphisms in synonymous and nonsynonymous sites in the *Exov* and the duplicated portion of *Evov-L* in a dataset of 709 *Exov* sequences and 455 *Exov-L* sequences computationally extracted from the *A. thaliana* accessions in the 1001 Genomes (The 1001 Genomes Consortium, 2016) (Table 2, Supplementary Table S5). In only 3.5 million years, *Exov* changed its protein sequence dramatically: 22 nonsynonymous substitutions led to a modification of 21 (15%) of the 136 amino acid residues that this gene encodes (Table 2). In contrast, the ancestral region of *Exov-L* evolved slowly, with only 3 amino acid residues changes. The McDonald-Kreitman test detected strong positive selection acting on *Exov* (Fisher exact test: two-tailed p = 0.0229). A high L value (=1-Neutral Index) of 0.82 revealed that a vast majority of the detected amino acid substitutions on *Exov* were driven by positive selection. *Exov-L*, on the other hand, evolved slowly, showing no signal of positive selection except, perhaps, a segregation of deleterious genetic variation, as its negative L value (−1.33) suggests.

### Molecular and expression analyses of *Exov* and *Exov-L*

Given that our evolutionary analysis revealed a signature consistent with a functional gene evolving under natural selection, we sought signals of functional evolution. First, we investigated changes in the molecular structure and sequence that have the potential to underlie functional change. Second, we assessed differences in the expression patterns of new and parental genes.

#### The new gene *Exov* was duplicated from the highly conserved region of the parental gene *Exov-L*

To understand the functional significance of the new gene *Exov*, we investigated the relationship between evolutionary changes in *Exov* and known molecular functions of the parental gene *Exov-L*.

We first examined the evolution of the parental gene *Exov-L*. Sequence alignment of *Exov-L* and its orthologs revealed high conservation from mammalian to plant species, especially within the N-terminal region in plants (Supplementary Figure S5a). Sequence alignment of *Exov-L* and its orthologs also showed high similarity in the DEM domain, which is known to encodes exonuclease (*EXO5* named in human and yeast) (Burgers et al., 2010; Sparks et al., 2012, Yeeles et al, 2009). One unique feature of this catalytic domain is its iron-sulfur cluster structure motif, which is a motif identified as an essential component of many DNA and RNA processing enzymes (White and Dillingham, 2012). The cysteine residues that form the critical Fe-S cluster motif in *EXOV-L* and its homologs in mammals and zebrafish are identical (Supplementary Figure S5a).

As shown in Figure 2a, the new gene *Exov* is a partial duplicate from the N-terminal region encoded by exon 1 (the *EXO5* homologous catalytic domain) of the parental gene *Exov-L.* Although *Exov-L* in plants is highly conserved in the N-terminal region, especially at positions R63, K85, and D103 (Supplementary Figure S5b), the conserved polar charged residues in the parental gene have been replaced in *EXOV* with more neutral histidine, isoleucine, and tyrosine residues, respectively (Supplementary Figure S5b). The corresponding region of AddB regulates the catalytic activity by forming contacts with AddA subunits (Supplementary Figure S5c). In contrast to the conservation defined by the parental gene *Exov-L*, which may be involved in the fine-tuned catalytic activities during DNA metabolism (Burgers et al., 2010; Sparks et al., 2012), the N-terminal region of the new gene *Exov* has accumulated many sequence changes. This variation indicates that *Exov* has evolved a smaller and distinct protein sequence with a diverged function.

#### Expression profiles of the new gene *Exov* and the parental gene *Exov-L* are overlapping

To quantify expression of the new and parental genes, we first performed RT-qPCR, using T-DNA mutant plants. We found that both *Exov* and *Exov-L* are transcribed in all tested organs: leaves, stems, flowers, and siliques. The results of our RT-qPCR experiments revealed that when compared to WT, *exov* and *exov-l* display significantly reduced expression in all the tissues except *exov* in siliques, where expression was often reduced by as much as 50% or more (Figure 4a).

**Figure 4.**
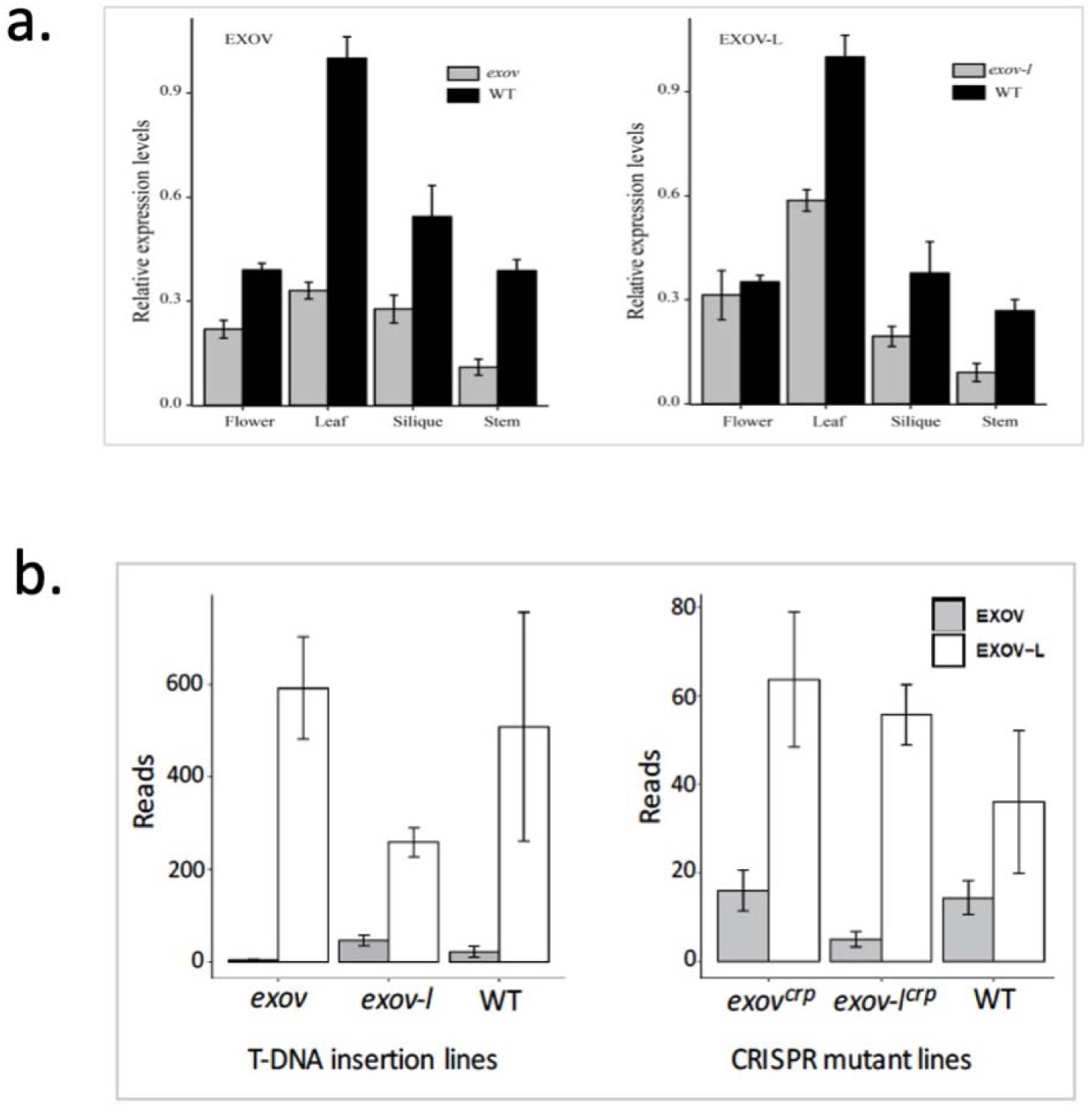
Expression analyses of mutants for Exov and Evov-L using RT-PCR and RNAseq. **a.** The expression levels in leaf of *Exov* and *Exov-L* in the wild-type are each set to 1. Relative expression of each gene in a specific tissue was calculated by normalizing to the value in WT plants. Error bars represent SE of triplicate experiments. The T-tests for the expression reduction in these organs in comparison to WT show all these except *exov* in silique are significant: *exov*: flower, p = 0.0015; leaf, p = 0.0015; silique, p = 0.3861; stem, 8.82e-05. *exov-l*: flower, p = 0.0101; leaf, p = 0.0408; silique, p = 0.0031; stem, p = 0.0069. **b.** The expression level of *Exov* and *Exov-L* in the transcriptomes generated by RNAseq of whole plants, presenting as FPKM (reads). The lines of *exov* and *exov-l* were created by T-DNA insertions; *exov^crp^* and *exov-l^crp^* were created using CRISPR/Cas9. WT is the wild-type line, Col-0. The standard error bars were derived from three biological replicates. T-tests for *exov-l* vs WT, p = 0.0168; for *exov* vs WT, p = 0.1230. T-tests for CRISPR mutant lines: *exov^crp^* vs WT, p = 0.7957; *exov-l^crp^* vs WT, p = 0.3524.

Reduced expression in T-DNA mutants of *Exov* and *Exov-L* is consistent with RNAseq transcriptome analyses of the whole plants, revealing significant or marginally significant reductions in expression, by as much as 50% (Figure 4b, T-DNA insertion lines). Our comparison of the transcriptomes of *exov* and *exov-l* with the wild-type revealed changes in the expression of 819 genes. Of these, 255 identical genes were shared between the expression networks of *Exov* and *Exov-L*. 361 genes uniquely changed expression in *exov* lines and 203 genes uniquely changed expression in *exov-l* lines. These data provide evidence for a functional divergence after the duplication of *Exov* from *Exov-L*, suggesting that *Exov* and *Exov-L* each interact to carry out unique functions (Supplementary Table S6).

#### The new gene *Exov* evolved to regulate additional biological processes beyond those regulated by the parental gene *Exov-L*

To better understand how the species-specific *Exov* gene diverged in its function as a consequence of distinct mutations, we generated specific mutations of *exov^crp^* and *exov-l^crp^* using the clustered regularly interspaced short palindromic repeats (CRISPR)/CRISPR-associated protein-9 nuclease (Cas9) system (Supplementary Figure S1). CRISPR/Cas9-induced mutants *exov^crp^* and *exov-l^crp^* with insertions (+) or deletions (-) at the desired target sites were identified (Figure 5). In order to assess changes in expression levels, we performed RT-qPCR for the wild-type, *exov*, *exov-l, exov^crp^,* and *exov-l^crp^*. *Exov* was duplicated from *Exov-L,* and as expected, their sequences are mostly identical (Supplementary Figure S5b). Because we could not distinguish the source of reads that could be mapped to both genes, we report only uniquely mapped reads for *exov* and *exov-l* in each sample (Figure 4b). In contrast to the T-DNA mutants, the expression levels of *Exov* and *Exov-L* in *exov^crp^* and *exov-l^crp^* lines do not appear to change significantly from the wild-type (for all the T-tests, 0.7957 > p > 0.1208) (Figure 4b). This may be a consequence of the specific single-nucleotide changes in *exov^crp^* and *exov-l^crp^* not changing the regulatory regions. The potential changes to functionality would be made by the reading frame shift by the single nucleotide deletions (Figure 5b and 5c). The asymmetric correlation of the parental gene and new gene in different mutants support a functional divergence after the duplication of *Exov* from *Exov-L*.

**Figure 5.**
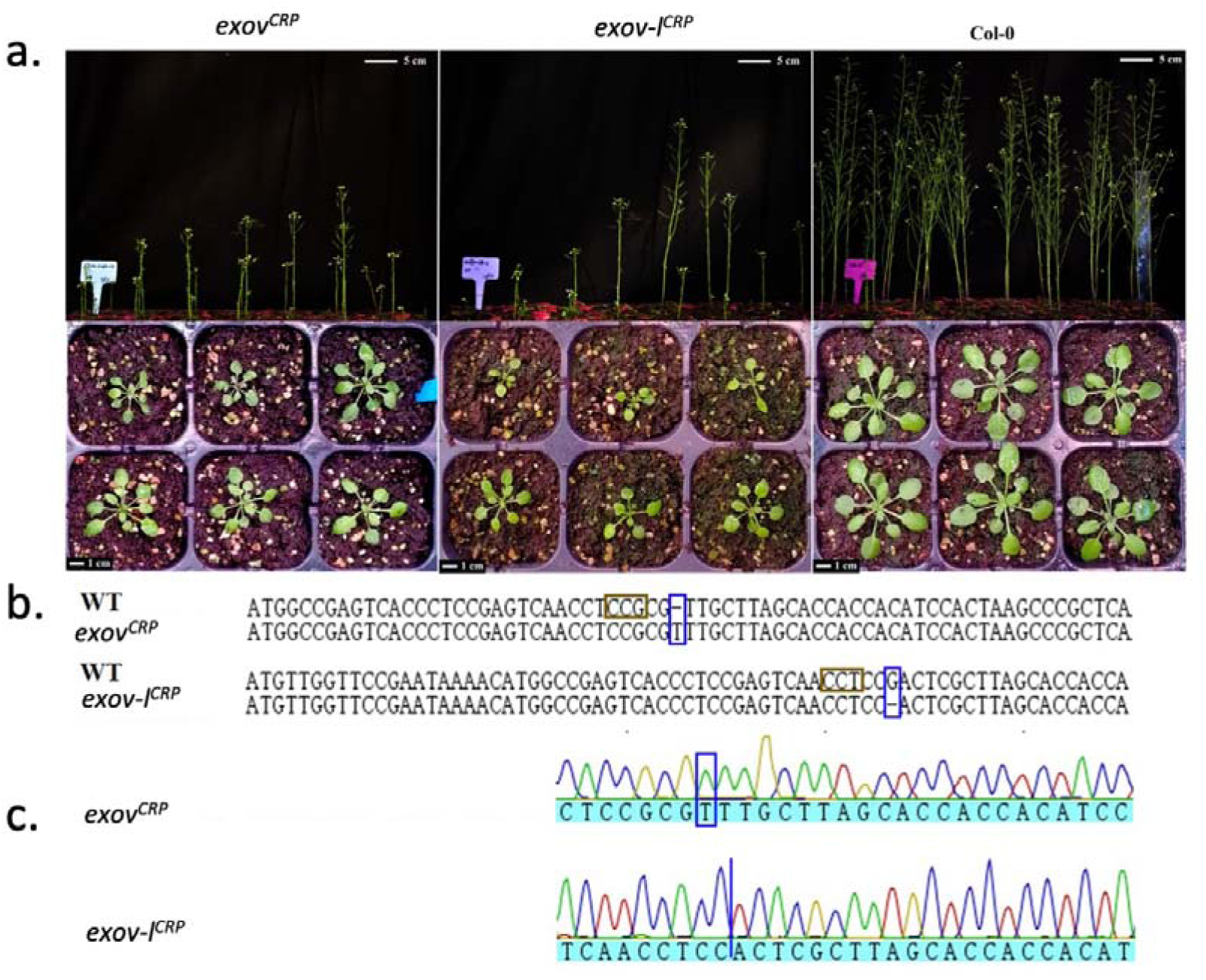
Generation of CRISPR-Cas9 mutants and measurement of their phenotypic effects. **a.** Phenotypes of T2 transgenic plants of the sgRNA target. *Exov-L* (AT5G60370): pCAMBIA1300-sgRNA T2 and *Exov* (AT3G57110): pCAMBIA1300-sgRNA T2 transgenic plants lines exhibited a small-seedling phenotype compared with the wild-type Col-0. Similar to T-DNA mutants of AT3G57110 and AT5G60370, they showed dwarfed and retarded growth. **b.** Representative sequences of several mutant alleles of sgRNA target identified from the *AT5G60370: pCAMBIA1300-sgRNA T2* and *AT3G57110: pCAMBIA1300-sgRNA T2* transgenic plants lines. The wild-type (WT) sequence is shown at the top with the PAM sequence highlighted in the red frame. Nucleotide deletion and insertion of transgenic lines were highlighted in the blue frames. **c**. DNA sequencing peaks showed evidence of successful gene editing in the target regions.

Based on these transcriptome data, we identified genes that were significantly differentially expressed in mutants versus WT despite a lack of difference in the level of *Exov* and *Exov-L* expression. In particular, 967 genes were down-regulated and 153 genes were up-regulated in *exov^crp^* relative to the wild-type. Meanwhile, 750 genes were down-regulated and 198 genes were up-regulated in *exov-l^crp^* (Supplementary Table S6c). Surprisingly, the new gene appears to interact with more genes (1,120 genes being down- or up-regulated if mutated, including both direct and indirect interactions) than does the parental gene (948 being down- or upregulated if mutated) (*X^2^=18.511, P= 1.689e-05,* under the null hypothesis of equal number of interacting genes). This pattern was also confirmed in T-DNA-insertion mutants, with 616 genes being down-/up-regulated (535/81) in *exov* compared to 458 genes being down-/up-regulated (340/118) in *exov-l* (*X^2^=26.863, P=* 2.185e-07) (Supplementary Table S6b). This provides a striking example of a recently formed gene evolving more interactions with other genes in the genome than the parental gene. This observation contrasts with the conventional view that new genes are integrated into the ancestral gene-gene interaction network and remain less integrated into cellular networks than old genes. It also provides a counter example to the observation of reduced levels of co-expression for new genes in mammalian evolution (Zhang et al, 2015).

Differentially expressed genes were ranked based on the p-values for simple t-tests comparing the wild-type and CRISPR/Cas9 mutants. The ranked list was used as input to GOrilla with default running parameters (Supplementary Figure S6). The results highlight a unique set of enriched GO terms that were identified at different cutoffs, including pollen tube development, pollination, multicellular organism processes, cell tip growth, cell morphogenesis involved in differentiation, developmental cell growth, pollen tube growth, aging, movement of the cell or subcellular components, and actin filament-based movement. While both the parental and new genes may be involved in aging, the new gene appears to additionally regulate novel biological processes such as the movement of the cell or subcellular components, including actin filament-based movement (Supplementary Figure S6), potentially explaining its increased genetic interactions. The information from the GO analyses suggests a valuable, albeit broad, picture of genetic mechanisms that, with further analysis, would enhance our understanding of the evolutionary forces on the parental and new genes that we investigated.

### Detection of the phenotypic effects of Exov and Exov-L on morphological traits

Our evolutionary analyses detected signatures of positive selection in the gene sequences, as well as the evolution of hundreds of new expression interactions involving the new gene. These evolutionary changes at the sequence and transcriptome levels are expected to have functional repercussions. To understand the functional divergence of *Exov* and *Exov-L*, we next scored seven important developmental traits in both wild-type plants and mutants harboring their CRISPR and T-DNA derived knockouts.

#### Seven morphological traits exhibit significant phenotypic effects in *Exov* and Exov-L

We measured and compared seven growth traits and flowering time among wild-type, T-DNA insertions and CRISPR/Cas9 knockout lines (Supplementary Figure S7; Supplementary Table S7).

In general, the mutants of *Exov* and *Exov-L* showed significant phenotypic effects compared to the wild-type in all seven traits examined (Figure 8. Supplementary Table S7). In 21 comparisons of T-DNA insertions (*exov, exov-l* and *exov/exov-l*) with wild-type, all are significant with p 0.00001 except *exov-l* for Branch number ≦ on the main bolt that is not significant (Wilcoxon rank sum test. The Gaussian-based test gave similar results). Among all 14 comparisons of CRISPR knockouts (*exov^crp^ and exov-l^crp^*) with the wild-type (Supplementary Table S7b, 11 with p 0.00001, ≦ only 2 (*Exov* in Rosette minor axis and *Exov-L* in Branch number) is not significant.

Further, we detected significant differences between *exov* and *exov-l in 5* of the 7 traits in the T-DNA insertions (p ≦ 0.0001, Wilcoxon rank sum test. The Gaussian-based test gave similar results) and similarly significant effects in 2 traits (Rosette major axis and Rosette minor axis). We detected significant differences between *exov* and *exov-l* in 4 of the 7 traits in the CRISPR knockouts and equally significant effects in other 3 traits (flowering time, Height and Branch on side bolts). In the cases of different effects between the two genes, *exov-l* more often has a stronger effect than *exov* (p < 2e-16). We observed that the plants in *exov* and *exov^crp^* were petite and displayed reduced growth rates (for example, Figure 5a). Remarkably, these mutants of the new gene *Exov* frequently show phenotypic effects as strong as the parental gene *Exov-l* whereas three traits even showed a stronger effect of *Exov* than *Exov-l* (*exov* in leaves number; *exov^crp^* in Height; *exov^crp^* in Main bolts number) (Figure 6, supplementary Table S7a). In general, we observed that all morphological traits examined differed significantly between the wild-type and mutants of the new gene and parental gene.

**Figure 6.**
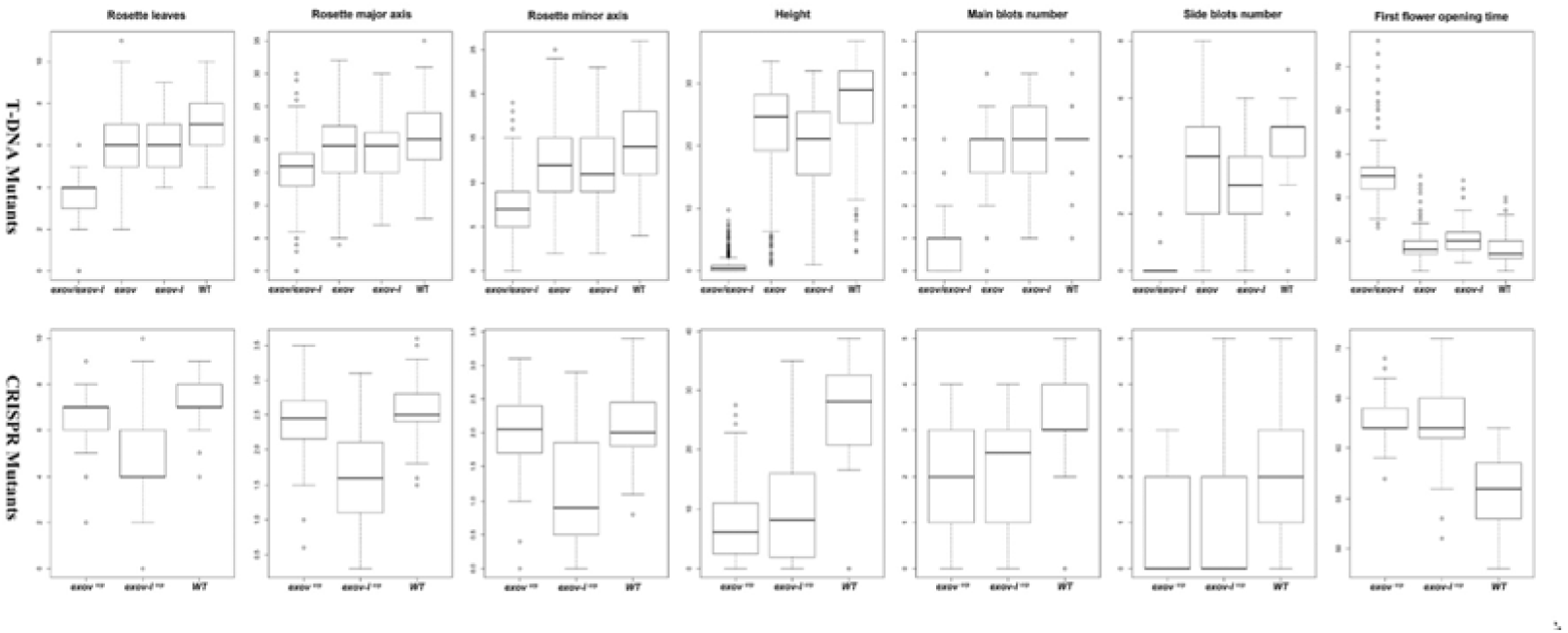
Distribution of phenotypic effects on seven traits of single *exov, exov-l* and double *exov, exov-l* mutants. Top: T-DNA insertions; Bottom: CRISP knockouts. WT: wildtype (Col-0).

Furthermore, the double mutant plants showed a strong and significant change in all 7 traits tested relative to single mutants and the wild-type (p < 2e-16, Wilcoxon rank sum test. The Gaussian-based test gave similar results) (Figure 8, top; Supplementary Table S7a). This observation suggests that the genetic bases of phenotypic changes in the two genes were not completely overlapping. For example, while the height of the main bolt reached 20-30 cm in 40-day-old plants of four single mutants and wild-type accessions, the double mutant did not produce a bolt within this time frame. In addition, the first flower did not open in the double mutant until 15 days later than in the single mutant and wild-type, suggesting stronger effects of the double mutant on these seven morphological traits.

We note that we determined the insertion sites for transgenic lines harboring T-DNA, including three wild-type allelic mutants for the new gene, using the whole genome sequencing. No additional insertion sites were detected in the mutant genomes. Using the similar genome sequencing, we confirmed that CRISPR/knockout lines are specific knockouts of both the new and parental gene, with no off-targets being detected in other parts of genomes.

#### Principal component analyses detected segregation of the phenotypic effects of mutants for *Exo*v and *Exov-L* from the wide-type genes

Principal component analysis was employed to obtain a global view of the differences between the phenotypes and across the mutants as represented in the data we created and described in Figure 6 and Supplementary Figure S7. PCA components 1 and 2 (Figure 7) contributed 58.8% and 14.5% for T-DNA insertion and 59.9% and 21.8% for the CRISPR mutants, respectively, to the total eigenvalues.

**Figure 7.**
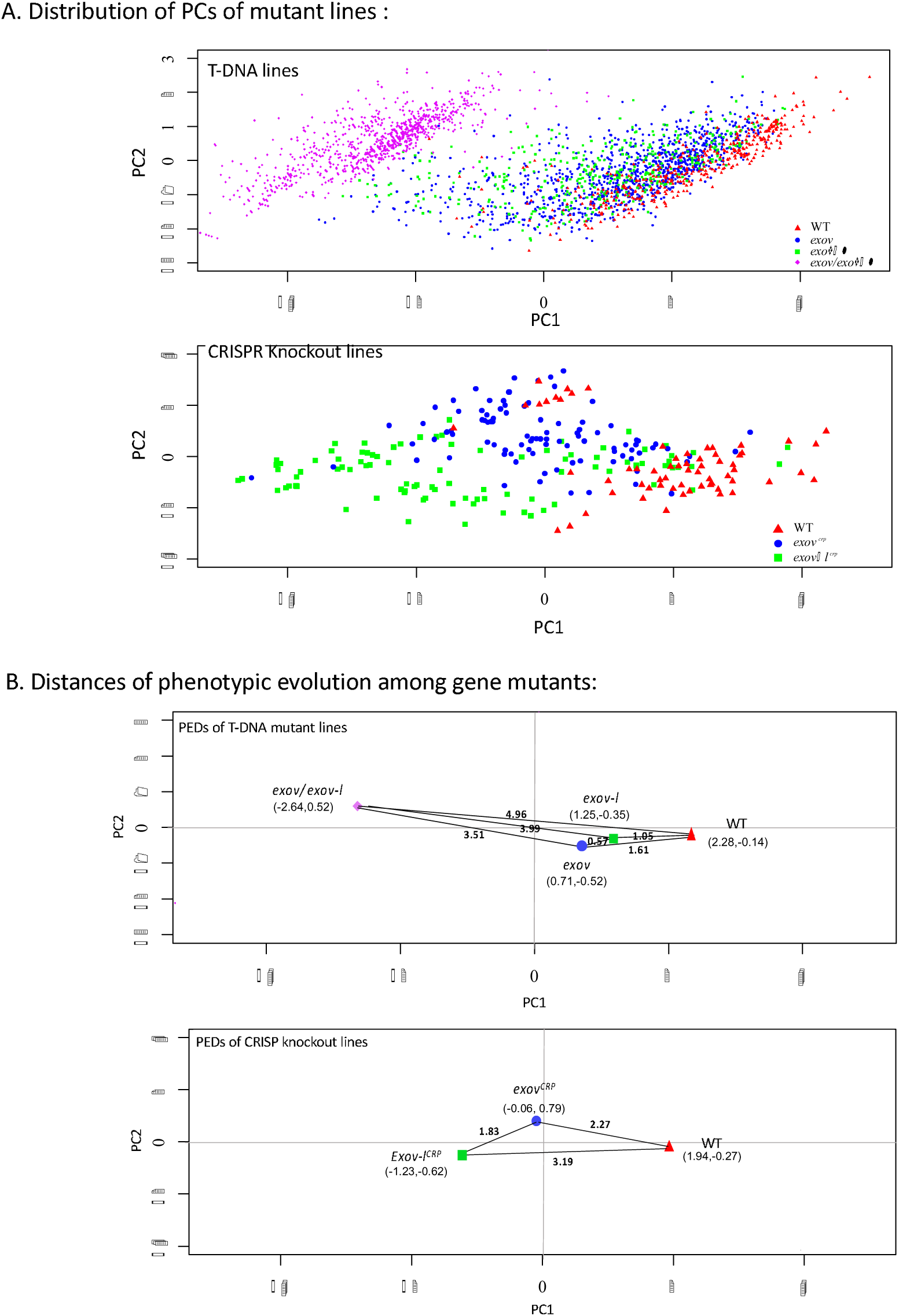
PCA analysis of the phenotypic effect of the new and parental genes and their distances of phenotypic evolution (PEDs). **a.** T-DNA insertions, Individual numbers: *exov*, 1098; *exov-l*, 389; double mutants, *exov/exov-l*, 1028; WT (Col-0), 413.B: CRISPR/Cas9 knockouts, individual numbers: *exov^crp^*, 96; *exov-l^crp^*, 96; WT, 64. **b.** the distance of phenotypic evolution (PED) among mutants defined as a geometric distance using the average values of PC1 and PC2 for each population (the pairs of coordinates in PC1 and PC2 respectively are given under each mutants and WT).

Interestingly, the two components in the two types of mutants showed remarkable segregation among wild-type, new gene mutant, and parental gene mutant plants. First, it is evident that mutants of both the new and old gene cause shifts away from the wild-type, revealing strong effects of these mutants on the overall phenotypes. Second, the mutants of *Exov* and *Exov-L* reveal distinct and separate distributions, revealing that phenotypic effects of *Exov* differ from those of *Exov-L*. Third, the long distances, 3.99 and 2.20, of phenotypic evolution (PED) of the double mutants *exov/exov-l* from single mutants *exov* and *exov-l* revealed additional phenotypic effects larger than the effects of the single mutants, 1.05 and 1.62. This reflects strong epistasic effects evolved by both *Exov* and *Exov-L*. Finally, the T-DNA insertions and CRISPR knockouts show a difference in the PED values between the single mutants and the wildtype: for the T-DNA insertion, *exov* > *exov-l* whereas for CRISPR KO *exov* < *exov-l*. This difference may reflect the difference in the mutations at transcriptional and translational levels. On the whole, the clear segregation of *exov* mutants (*exov* and *exov^crp^*, blue) from the wild-type and the mutants of the parental gene *Exov-L* reveals that the species-specific gene *Exov* evolved novel and strong phenotypic effects in a period of time as short as 3.5 MYA.

## Discussion

As our ability to study the roles of new genes in phenotypic evolution has become feasible, the importance of these genes is becoming apparent.. The present study reveals for the first time that a species-specific gene in *Arabidopsis* plays an important role in the phenotypic evolution of *A. thaliana*. We found that all seven major quantitative traits in development and reproduction are significantly impacted by the mutations of the species-specific *Exov* created by the T-DNA insertions and CRISPT-Cas9 knockout.

It is also remarkable that *Exov* developed more expression-interactions than the old parental gene *Exov-L*. It is important to note that these unexpected evolutional changes at the molecular and phenotypic levels were driven by the detected strong positive selection.

Our nucleotide substitution analyses revealed a Ka/Ks ratio much less than 1 in the new gene, *Exov,* suggesting strong selective constraints in the new gene *Exov*. Despite the young age of *Exov,* which was generated through gene duplication ∼3.5 million years ago, its divergence in nonsynonymous sites from the *Exov-L* reached a surprisingly high level of 14%. Further, the McDonald-Kreitman test detected a significant excess of nonsynonymous substitution compared to the within-species variation at nonsynonymous and synonymous sites. These analyses further detected that the protein sequence encoded by *Exov* evolved ∼7 times more rapidly than *Exov-L*, suggesting the significant impact of positive selection driving the neofunctionalization of *Exov*.

The old gene, *Exov-L,* possesses a highly conserved DEM (defects in morphology) domain, and members of this family of proteins were found to have exonuclease functions (Burgers et al., 2010; Sparks et al., 2012). However, no conserved domains have been identified in the new gene *Exov*, suggesting a recent appearance in *A. thaliana* of this novel gene may lead to a new function. Consistent with the analysis of the chloroplast transit signal prediction, the final destination of both new and old proteins is predicted to be the chloroplast (Bosco, 2003). The homologous gene to *Exov-L* is highly conserved across humans and yeast, where it has been shown to be involved in DNA metabolism and genome stability in mitochondria (Burgers et al., 2010; Sparks et al., 2012).

Our prediction that the new gene *Exov* is functional is further supported by the significant phenotypic effects on the morphological traits in T-DNA and CRISPR/Cas9 mutated lines. Interestingly, the new gene *Exov* shows a robust signal indicating positive selection in the N-termini. The residues of this regulatory domain evolved to give rise to new functional roles of *Exov*, but the catalytic domain was lost. This type of protein evolution implicates a fundamental role for proteins to gain new functions.

Furthermore, we found significant segregation of the phenotypic effects of the new gene versus the old gene among seven traits that are at least partially independent. Strong evidence for functional divergence introduced by the new gene was detected by PCA. The distribution of PCA scores showed functional shifts among mutants of the new gene and old gene. Unexpectedly, given the young age of *Exov*, these analyses detected a tremendous divergence from the parental gene to this new, species-specific gene, suggesting its critical roles in the evolution of morphological traits. Surprisingly, the T-DNA insertions and CRISPR Knockouts revealed that the new gene *Exov* can have as equal as or stronger effects on a few morphological straits than the old parental duplicate copy *Exov-L*. The whole genome sequencing of the mutant lines confirmed that these phenotypic effects were not caused by background mutations such as additional T-DNA insertions or CRISPR off-targets elsewhere in genomes. Furthermore, the multiple mutant lines revealed similar phenotypic effects support that the observed phenotypic effects are consequence of the mutations created in these lines.

Moreover, though both new and parental genes may be involved in the biosynthesis of secondary metabolites, the RNAseq comparison of the gene mutants and wild-type revealed that the new gene had evolved many more genetic interactions than the old genes (Supplementary Table S8). To our knowledge, this is the first example in plants in which a young gene quickly evolved many more co-expression interactions with other genes in the genome. The large number of interactions suggests a hub in genome interaction networks, potentially explaining its significant impact on morphological trait divergence and detected strong epistasis effects detected in T-DNA double mutants (Figure 9. A2). These newly evolved interactions give insight into the evidence for positive selection on phenotypic evolution, as well as suggesting that the new gene may have contributed to the phenotypic evolution underlying the examined morphological traits in *A. thaliana* through a neofunctionalization process.

Gene and mutants accession number:

*Exov*: new gene AT3G57110

*Exov-L*: parental gene AT5G60370

*exov*: AT3G57110 T-DNA insertion mutant

*exov-1*(Salk-103969), *exov-2* (Salk-036494), *exov-3* (Salk-064431)

*exov-l*: AT5G60370 T-DNA insertion mutant *exov-l* (Salk-101821)

*exov^crp^*: AT3G57110 CRISPR Cas9 mutant

*exov-l ^crp^*: AT5G60370 CRISPR Cas9 mutant

## Author Contributions and Acknowledgments

Y.H., M.L., and J.B. designed this research. Y. H. and J. C. performed the experiments and analysis, with significant contributions from J. B. and M. L. C. F. provided plant materials. C. F., Y. O., D. L., and E. M. revised the manuscript. Y. H., J. C., M. L., and J. B. wrote the manuscript with contribution from all authors.

This study was supported by the National Key Basic Research Program of China grant (Grant 2014CB954100) to D.L., the National Science Foundation grant (NSF1026200) to M.L., the NIH grant (R01GM83068) to J.B., the Natural Science Foundation of China (31560062) and Yunnan Education Department grant (2015Z057) to Y.H., and the scholarship from the Chinese Academy of Sciences and China Scholarship Council to Y.H. We are thankful for the valuable discussion with the members in the laboratories of M.L. and C. F. We are indebted to the technical help of John Zdenek, Sandra Suwanski and Qian Yang.

## Legends of Supplementary Files/Tables and Supplementary Figures

**Supplementary file 1**. Mapping the chromosomal insertion positions of the corresponding T-DNA lines of *Exov* and *Exov-l*.

**Supplementary file 2.** Mapping the on-targets of *Exov* and *Exov-l* in CRISPR KO lines.

## Supplementary Tables

**Table S1. Summary of the whole genome sequencing in the T-DNA insertion lines and CRISPR-target lines.**

**Table S2. Used for Allele-Specific PCR,RT-PCR and RT-qPCR Reactions.**

**Table S3. Measurements of phenotypic analysis.**

**Table S4. *A. thaliana* accessions for population structure analysis.**

**Table S5. The data of substitutions and polymorphisms for the McDonald-Kreitman test of positive selection.**

**Table S6. a. GO enrichment analysis of the set of genes that significantly differentially expressed between *exov* and *exov-l.* b. The significantly differentially expressed genes between *exov* and *exov-l. c.* The significantly differentially expressed genes between *exov^crp^* and *exov-l^crp^***

**Table S7. Pairwise comparisons using Wilcoxon rank sum tests for phenotypic traits of T-DNA mutants and CRISPR-Cas9 mutants.**

**Table S8. a. GO enrichment of analysis of the set of genes that significantly differentially expressed between wild type and *exov-l*. b. GO enrichment analysis of the set of genes that was significantly differentially expressed between wild type and *exov*.**

## Supplementary Figures

**Figure S1. Sequence, annotation and restriction map of pCAMBIA1300**

**Figure S2. Summary of neutrality test pipeline.**

**Figure S3. Analyses of population structure for the world-wide accessions used in this study (the 1001 Genomes). a.** Population structure under different assumptions about the number of clusters (K=2, 3, 4, 5, 6, 7). **b.** The cross-validation errors at various K values. **c.** Population structure analysis of 851 worldwide *A. thaliana* accessions (K = 8).

**Figure S4. The empirical distributions of several population genetic test parameters across the genome in *A. thaliana* and the probabilities of *Exov* and *Exov-l* in these distributions.**

**Figure S5. Protein sequence divergences of EXOV-L and EXOV. a.** Alignment of EXOV-L (AT5G60370) with its homologs from _erent_ species and AddB (*B.subtilis*). These homologs are from Human (NP_073611.1), Chimpanzee (XP_003308065.1), Monkey (XP_001084006.1), Mouse (NP_001153515.1), Rat (NP_001101443.1), Dog (XP_532542.1), Cattle (NP_001075077.1), Zebrafish (NP_001032490.1), *M. oryzae* (XP_003718794.1), and *N. crassa* (XP_955908.1). The conserved Cysteine residues that coordinate the Fe-S cluster are highlighted in red. **b**. Alignment of EXOV (AT3G57110) and EXOV-L with its orthologs in the plant. The conserved polar residues at positions 63, 85, and 103 of AT5G60370 and their orthologs are highlighted in red. At position 63 of AT5G60370, the conserved residue is the basic polar residue arginine (R). In AT3G57110, this residue evolved to histidine (H). At position 85, the residue is either basic polar residue lysine (K) or arginine (R) in all instances except for that of AT3G57110, where it is substituted with the hydrophobic residue isoleucine (I). The conserved residue at position 103 is the acidic charged residue aspartate (D), which is changed to tyrosine (Y) in AT3G57110. Other residues such as R77, I78, T79, S102, and A119 were substituted with Q77, M78, I79, L102, and S119, highlighted in green. The NCBI accession number for the orthologs from *A. lyrata, C. rubella, E. salsugineum, J. curcas,* apple, and tomato are XP_002864682, XP_006280574, XP_006400854, KDP44101, XP_008358302, and XP_004251259. **c.** The proposed structural model of EXOV-L showing its conservation.

**Figure S6. GO analyses.** a. GO enrichment analysis of the set of genes that is significantly differentially expressed between wild type and *exov^crp^*. b. GO enrichment of analysis of the set of genes that is significantly differentially expressed between wild type and *exov-l^crp^*.

**Figure S7. Distribution of the phenotypic effects on seven traits of T-DNA mutants lines (single *exov, exov-l* and double *exov/exov-l*) and CRISPR/Cas9 mutant lines (*exov^crp^, exov-l^crp^*) of the new gene and parental gene and wild type lines (Col-0).** The curves are theoretical distributions modelled as Gaussian distribution. The numbers of individual plants were measured and used to generate these distributions: 1. *exov*, 1098 (3 insertion mutants); 2. *exov-l*, 389; 3. Double mutants, *exov/exov-l*, 1028 (3 *exov* insertion mutants x *exov-l*); 4. WT for the insertion mutants, 413; 5. *exov^crp^*, 96; 6. *exov-l^crp^*, 96; 7. WT for the two CRISPR knockouts, 64.

