## Supplementary Table, Supplementary Figure for "Species-specific gene duplication in *Arabidopsis thaliana* evolved novel phenotypic effects on morphological traits under strong positive selection"

**Supplementary materials**

Table S1. Identification of T-DNA insertion and CRIRSPR target site in the mutant lines by using whole Genome Sequencing through Illumina Sequencing.

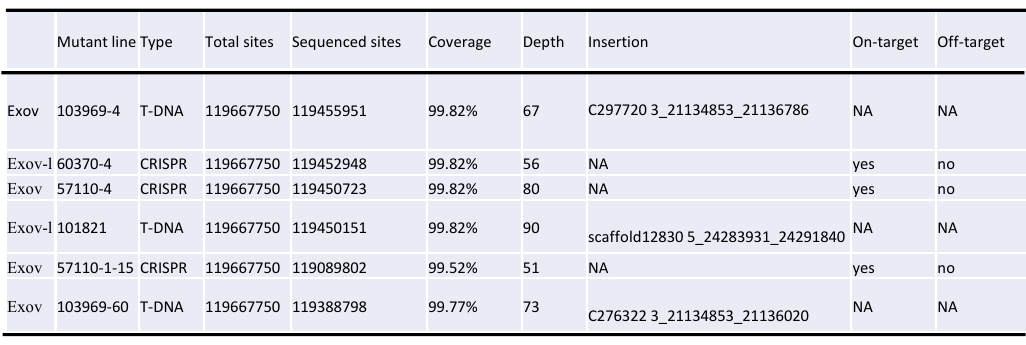

Table S2. Primers used for allele-specific PCR,RT-PCR and RT-qPCR reactions

| **Name** | **Primer sequence** | **Size (bp)** |
| --- | --- | --- |
| AT3G57110-LP: | 5’ TCTCACTTCTTTGTCCTTTT 3’ | 810 |
| AT3G57110-RP: | 5’ ATAAGAAACCCAAACAACAA 3’ |  |
| AT5G60370-1LP | 5’ TTAAGAACATGCAAAACTG 3’ | 1201 |
| AT5G60370-1RP | 5’ TTGACCTCCAACAAATCCT 3’ |  |
| AT5G60370-2LP | 5’ TGATTGTAGTTGAGTGGGAA 3’ | 919 |
| AT5G60370-2RP | 5’ CATAAATTTGAGGTTATTGG 3’ |  |
| AT5G60370-3LP | 5’ TAGCAAATTGGCAATACCGAC 3’ | 1082 |
| AT5G60370-3RP | 5’ AGCTGTTGAATTCCATTGCTG 3’ |  |
| AT3G57110-RT 1S | 5’ ATTTCAAGCCCTCAAAGTCA 3’ | 303 |
| AT3G57110-RT 1R | 5’ CAAACGTACAGTACCGGTGA 3’ |  |
| AT3G57110-RT 2S | 5’ GTCACCCTCCGAGTCAACCT 3’ | 297 |
| AT3G57110-RT 2R | 5’ AAACAAGCTGATAATTTCCT 3’ |  |
| AT5G60370-RT 1S | 5’ TTAGCACCACCACATCCACT 3’ | 429 |
| AT5G60370-RT 1R | 5’ AATGACGAGCCTGACCAACT3’ |  |
| Oligo(dT)20 | 5' TTTTTTTTTTTTTTTTTTTT 3' |  |
| TUB2-RTF | 5'GTTCTCGATGTTGTTCGTAAG3' |  |
| TUB2-RTS | 5'TGTAAGGCTCAACCACAGTAT3' |  |
| At3g57110-qRTF | 5' CATCCACTAAGCCCGCTCAA 3' |  |
| At3g57110-qRTR | 5' TGCAAGAGCTGCATCGAGAA 3' |  |
| AT5G60370-qRTF | 5' CTTCCCCGAGATCCCAATCG 3' |  |
| AT5G60370-qRTR | 5' ATCACGGAAGGGAGGATGGA 3' |  |
| Act8-qRTF | 5' TCAGCACTTTCCAGCAGATG 3' |  |
| Act8-qRTR | 5' CTGTGGACAATGCCTGGAC 3' |  |

Table S3. Measurements of phenotypic analysis

| **Measurement** | **unit** | **Growth stage** | **Description** |
| --- | --- | --- | --- |
| number of rosette leaves | count | 1.04 | 4 rosette leaves > 1mm in length |
| rosette major axis | mm | 1.08 | 8 rosette leaves > 1mm in length |
| rosette minor axis | mm | 1.08 | 8 rosette leaves > 1mm in length |
| Height | mm | 6.10 | 10% flower to be produced have opened |
| Bolt | count | 6.10 | 10% flower to be produced have opened  Main bold number  Side bolt number |
| First flower open | day | 6.00 | First flower open |

Table S4. *A. thaliana* accessions for population structure analysis

| **No.** | **Accession No.** | **Country of Origin** | **No.** | **Accession No.** | **Country of Origin** | **No.** | **Accession No.** | **Country of Origin** |
| --- | --- | --- | --- | --- | --- | --- | --- | --- |
| 1 | Pog_0 | Canada | 236 | Chat_1 | France | 471 | Kni-1 | Sweden |
| 2 | Van_0 | Canada | 237 | Di_G | France | 472 | Kor-3 | Sweden |
| 3 | Ri-0.JGI | Canada | 238 | Et_0 | France | 473 | Kru-3 | Sweden |
| 4 | Bg_2 | USA | 239 | Gy_0 | France | 474 | Kulturen-1 | Sweden |
| 5 | Buckhorn_Pass | USA | 240 | Ra_0 | France | 475 | Lan-1 | Sweden |
| 6 | CHA-41.932 | USA | 241 | Rennes_1 | France | 476 | Liarum | Sweden |
| 7 | Col-0.MPI | USA | 242 | Rou_0 | France | 477 | Lilloe-1 | Sweden |
| 8 | CSHL-5.6744 | USA | 243 | Com_1 | France | 478 | Lis-2 | Sweden |
| 9 | Dem_4 | USA | 244 | Lm_2 | France | 479 | Lis-3 | Sweden |
| 10 | Gre_0 | USA | 245 | An_1 | Belgium | 480 | Lund | Sweden |
| 11 | Knox_18 | USA | 246 | Ang_0 | Belgium | 481 | Oemoe1-7 | Sweden |
| 12 | KYC-33.801 | USA | 247 | Dolen-1.9697 | Belgium | 482 | Oemoe2-1 | Sweden |
| 13 | LI-OF-065.630 | USA | 248 | Dolna-1.9712 | Belgium | 483 | Oer-1 | Sweden |
| 14 | LP3413.41.8472 | USA | 249 | Dospa-1.9706 | Belgium | 484 | Rev-1 | Sweden |
| 15 | Mdn-1.1829 | USA | 250 | Goced-1.9698 | Belgium | 485 | Rev-2 | Sweden |
| 16 | MIC-31.870 | USA | 251 | Grivo-1.9714 | Belgium | 486 | Roed-17-319 | Sweden |
| 17 | MNF-Che-2.1925 | USA | 252 | Kolar-1.9699 | Belgium | 487 | Sim-1 | Sweden |
| 18 | MNF-Jac-12.1954 | USA | 253 | Kolar-2.9702 | Belgium | 488 | Sparta-1 | Sweden |
| 19 | MNF-Pin-39.2016 | USA | 254 | Koren-1.9719 | Belgium | 489 | Spr1-2 | Sweden |
| 20 | MNF-Pot-21.1853 | USA | 255 | Leska-1.9716 | Belgium | 490 | Spro-1 | Sweden |
| 21 | MNF-Pot-75.1872 | USA | 256 | Schip-1.9721 | Belgium | 491 | Spro-2 | Sweden |
| 22 | MNF-Riv-21.1890 | USA | 257 | Slavi-2.9723 | Belgium | 492 | Spro-3 | Sweden |
| 23 | Mv_0 | USA | 258 | Smolj-1.9718 | Belgium | 493 | Sr:3 | Sweden |
| 24 | NC_6 | USA | 259 | Stara-1.9713 | Belgium | 494 | Ste-3 | Sweden |
| 25 | Paw-26.2171 | USA | 260 | Hi-0.WTC | Netherlands | 495 | Ste-4 | Sweden |
| 26 | PNA3.40.7947 | USA | 261 | Lisse | Netherlands | 496 | Spr1-6 | Sweden |
| 27 | Pna_10 | USA | 262 | Amel_1 | Netherlands | 497 | T1000 | Sweden |
| 28 | Pna_17 | USA | 263 | Appt_1 | Netherlands | 498 | T1020 | Sweden |
| 29 | PT2.21.8077 | USA | 264 | Gel_1 | Netherlands | 499 | T1070 | Sweden |
| 30 | PT2.21.8077 | USA | 265 | Hey_1 | Netherlands | 500 | T1080 | Sweden |
| 31 | RMX3.22.8132 | USA | 266 | Le_0 | Netherlands | 501 | T1090 | Sweden |
| 32 | Rmx_A02 | USA | 267 | Nok_3 | Netherlands | 502 | T1110 | Sweden |
| 33 | Rmx_A180 | USA | 268 | Rhen_1 | Netherlands | 503 | T1130 | Sweden |
| 34 | RRs_10 | USA | 269 | Utrecht | Netherlands | 504 | T470 | Sweden |
| 35 | RRS_7 | USA | 270 | Baa_1 | Netherlands | 505 | T480 | Sweden |
| 36 | Seattle_0 | USA | 271 | Benk_1 | Netherlands | 506 | T530 | Sweden |
| 37 | SLSP-31.2276 | USA | 272 | Tha_1 | Netherlands | 507 | T540 | Sweden |
| 38 | SLSP-35.2278 | USA | 273 | Hau_0 | Denmark | 508 | T550 | Sweden |
| 39 | Tol_0 | USA | 274 | Ko_2 | Denmark | 509 | T570 | Sweden |
| 40 | Tul_0 | USA | 275 | Alt-1.9774 | Germany | 510 | T710 | Sweden |
| 41 | WAR.7477 | USA | 276 | Bach2-1.9796 | Germany | 511 | T720 | Sweden |
| 42 | Yo_0 | USA | 277 | Bach-7.9778 | Germany | 512 | T740 | Sweden |
| 43 | Agu-1.MPI | Spain | 278 | Bai-10.9779 | Germany | 513 | T780 | Sweden |
| 44 | Bla_1 | Spain | 279 | Berg-1.9775 | Germany | 514 | T790 | Sweden |
| 45 | Cdm-0.MPI | Spain | 280 | BI-4.9813 | Germany | 515 | T800 | Sweden |
| 46 | Don-0.MPI | Spain | 281 | Can-0.WTC | Germany | 516 | T840 | Sweden |
| 47 | Leo-1.MPI | Spain | 282 | Erg2-6.9784 | Germany | 517 | T850 | Sweden |
| 48 | Mer-6.MPI | Spain | 283 | Fell1-10.9814 | Germany | 518 | T860 | Sweden |
| 49 | Ped-0.MPI | Spain | 284 | Fell2-4.9780 | Germany | 519 | T880 | Sweden |
| 50 | Pla_0 | Spain | 285 | Fell3-7.9776 | Germany | 520 | T900 | Sweden |
| 51 | Pra-6.MPI | Spain | 286 | Gn-1.9777 | Germany | 521 | T930 | Sweden |
| 52 | Se_0 | Spain | 287 | Gn2-3.9790 | Germany | 522 | T960 | Sweden |
| 53 | Vie-0.MPI | Spain | 288 | Got_22 | Germany | 523 | T980 | Sweden |
| 54 | Pro_0 | Spain | 289 | Ha-HBT1-2.9785 | Germany | 524 | T990 | Sweden |
| 55 | Ts_1 | Spain | 290 | Ha-HBT2-10.9797 | Germany | 525 | TAAL-03 | Sweden |
| 56 | Bla-1.7015 | Spain | 291 | Ha-HBT3-11.9815 | Germany | 526 | TAAL-07 | Sweden |
| 57 | IP-Adm-0.9514 | Spain | 292 | Ha-P-13.9786 | Germany | 527 | TBO-01 | Sweden |
| 58 | IP-Ala-0.9515 | Spain | 293 | Ha-P2-1.9798 | Germany | 528 | TDr-13 | Sweden |
| 59 | IP-All-0.9517 | Spain | 294 | Ha-S-B.9800 | Germany | 529 | TDr-16 | Sweden |
| 60 | IP-Alm-0.9518 | Spain | 295 | Ha-SP-2.9801 | Germany | 530 | TDr-17 | Sweden |
| 61 | IP-Ang-0.9519 | Spain | 296 | Haes-1.9791 | Germany | 531 | TDr-2 | Sweden |
| 62 | IP-Ara-4.9520 | Spain | 297 | Hart-2.9799 | Germany | 532 | TDr-7 | Sweden |
| 63 | IP-Bar-1.9521 | Spain | 298 | HE-1.9769 | Germany | 533 | TDr-8 | Sweden |
| 64 | IP-Bea-0.9522 | Spain | 299 | Hof-1.9772 | Germany | 534 | TEDEN-02 | Sweden |
| 65 | IP-Ber-0.9524 | Spain | 300 | KBG1-14.9788 | Germany | 535 | TOM-07 | Sweden |
| 66 | IP-Cab-3.9526 | Spain | 301 | KBG2-13.9770 | Germany | 536 | Tottarp-2 | Sweden |
| 67 | IP-Cad-0.9527 | Spain | 302 | Koln | Germany | 537 | Tur-4 | Sweden |
| 68 | IP-Cal-0.9528 | Spain | 303 | Kus2-2.9781 | Germany | 538 | TV-10 | Sweden |
| 69 | IP-Cap-1.9529 | Spain | 304 | Ler-0.WTC | Germany | 539 | TV-22 | Sweden |
| 70 | IP-Car-1.9530 | Spain | 305 | Li-7.7231 | Germany | 540 | TV-30 | Sweden |
| 71 | IP-Cdc-3.9531 | Spain | 306 | Lu3-30.9782 | Germany | 541 | TV-38 | Sweden |
| 72 | IP-Cdo-0.9532 | Spain | 307 | Lu4-2.9792 | Germany | 542 | TV-7 | Sweden |
| 73 | IP-Coc-1.9535 | Spain | 308 | Muh-2.9803 | Germany | 543 | Ull2-3 | Sweden |
| 74 | IP-Cor-0.9536 | Spain | 309 | Obe1-15.9804 | Germany | 544 | Ull2-5 | Sweden |
| 75 | IP-Cum-1.9537 | Spain | 310 | Obh-13.9789 | Germany | 545 | Vaar-1 | Sweden |
| 76 | IP-Cur-4.9538 | Spain | 311 | Pfn-10.9805 | Germany | 546 | Vaar2-1 | Sweden |
| 77 | IP-Deh-1.9539 | Spain | 312 | Pfn-N2.2-6.9771 | Germany | 547 | Vaar2-6 | Sweden |
| 78 | IP-Elb-0.9540 | Spain | 313 | Po-0.WTC | Germany | 548 | Vimmerby | Sweden |
| 79 | IP-Fue-2.9541 | Spain | 314 | Rd-0.8411 | Germany | 549 | Vinsloev | Sweden |
| 80 | IP-Fun-0.9542 | Spain | 315 | Ru-2.9806 | Germany | 550 | Yst-1 | Sweden |
| 81 | IP-Gra-0.9543 | Spain | 316 | Ru-N2.9793 | Germany | 551 | Ting_1 | Sweden |
| 82 | IP-Gua-1.9544 | Spain | 317 | Ru4-16.9768 | Germany | 552 | Tamm-2 | Finland |
| 83 | IP-Her-12.9545 | Spain | 318 | Schl-7.9807 | Germany | 553 | Tamm_27 | Finland |
| 84 | IP-Cmo-3.9534 | Spain | 319 | Ste-0.7346 | Germany | 554 | Tamm_2 | Finland |
| 85 | IP-Hor-0.9547 | Spain | 320 | Tu-B1-2.9794 | Germany | 555 | In_0 | Austria |
| 86 | IP-Jim-1.9551 | Spain | 321 | Tu-B2-3.9808 | Germany | 556 | Pi_0 | Austria |
| 87 | IP-Lab-7.9552 | Spain | 322 | Tu-KB-6.9809 | Germany | 557 | Tscha_1 | Austria |
| 88 | IP-Ldd-0.9553 | Spain | 323 | Tu-KS-7.9810 | Germany | 558 | Uod_1 | Austria |
| 89 | IP-Lso-0.9554 | Spain | 324 | Tu-NK-12.9811 | Germany | 559 | Gr-5.7158 | Austria |
| 90 | IP-Mar-1.9555 | Spain | 325 | Tu-PK-7.9783 | Germany | 560 | Uod-2.8428 | Austria |
| 91 | IP-Moc-11.9558 | Spain | 326 | Tu-WH.9816 | Germany | 561 | Uod_7 | Austria |
| 92 | IP-Mon-5.9559 | Spain | 327 | Aa_0 | Germany | 562 | Blh-1.JGI | Czech |
| 93 | IP-Mot-0.9560 | Spain | 328 | Ak_1 | Germany | 563 | Borky1.428 | Czech |
| 94 | IP-Mun-0.9561 | Spain | 329 | Bch_1 | Germany | 564 | Doubravnik7.410 | Czech |
| 95 | IP-Mur-0.9562 | Spain | 330 | Bd_0 | Germany | 565 | Draha2.424 | Czech |
| 96 | IP-Nav-0.9563 | Spain | 331 | Bsch_0 | Germany | 566 | DraII_1 | Czech |
| 97 | IP-Hum-2.9549 | Spain | 332 | Bu_0 | Germany | 567 | DraII-6.5874 | Czech |
| 98 | IP-Iso-4.9550 | Spain | 333 | Db_1 | Germany | 568 | DraIII_1 | Czech |
| 99 | IP-Orb-10.9565 | Spain | 334 | Ei_2 | Germany | 569 | DraIV.5893 | Czech |
| 100 | IP-Oso-0.9566 | Spain | 335 | El_0 | Germany | 570 | DraIV.5907 | Czech |
| 101 | IP-Pal-0.9567 | Spain | 336 | En_2 | Germany | 571 | DraIV.5950 | Czech |
| 102 | IP-Pue-0.9572 | Spain | 337 | Est | Germany | 572 | DraIV.5993 | Czech |
| 103 | IP-Rds-0.9573 | Spain | 338 | Ey15-2.MPI | Germany | 573 | Duk | Czech |
| 104 | IP-Rel-0.9574 | Spain | 339 | Fi_0 | Germany | 574 | Hod | Czech |
| 105 | IP-Ren-6.9575 | Spain | 340 | Fr_2 | Germany | 575 | HSm | Czech |
| 106 | IP-Rev-0.9576 | Spain | 341 | Gie_0 | Germany | 576 | Lp2_6 | Czech |
| 107 | IP-Sac-0.9578 | Spain | 342 | Got_7 | Germany | 577 | Pu2_8 | Czech |
| 108 | IP-San-10.9579 | Spain | 343 | Gu_0 | Germany | 578 | Sap_0 | Czech |
| 109 | IP-Scm-0.9580 | Spain | 344 | HKT2.4.MPI | Germany | 579 | UduI.6296 | Czech |
| 110 | IP-Sdv-3.9581 | Spain | 345 | Hn_0 | Germany | 580 | UduI.6390 | Czech |
| 111 | IP-Ses-0.9582 | Spain | 346 | Hs_0 | Germany | 581 | UduI.6396 | Czech |
| 112 | IP-Sne-0.9583 | Spain | 347 | Is_0 | Germany | 582 | ZdrI.6424 | Czech |
| 113 | IP-Stp-0.9584 | Spain | 348 | Kro_0 | Germany | 583 | ZdrI.6434 | Czech |
| 114 | IP-Svi-0.9585 | Spain | 349 | Kro-0.MPI | Germany | 584 | ZdrI.6445 | Czech |
| 115 | IP-Tam-0.9586 | Spain | 350 | Krot_0 | Germany | 585 | Sav-0.UNIL | Czech |
| 116 | IP-Tdc-0.9587 | Spain | 351 | La_0 | Germany | 586 | Bor_4 | Czech |
| 117 | IP-Tor-1.9589 | Spain | 352 | Ler-1.MPI | Germany | 587 | Br_0 | Czech |
| 118 | IP-Trs-0.9590 | Spain | 353 | Ler-1 | Germany | 588 | Da1_12 | Czech |
| 119 | IP-Vad-0.9591 | Spain | 354 | Li_2:1 | Germany | 589 | Dra_0 | Czech |
| 120 | IP-Vae-2.9592 | Spain | 355 | Mnz_0 | Germany | 590 | Jl_3 | Czech |
| 121 | IP-Vaz-0.9593 | Spain | 356 | Mz_0 | Germany | 591 | Lp2_2 | Czech |
| 122 | IP-Vdm-0.9594 | Spain | 357 | Nie1-2.MPI | Germany | 592 | Pu2_23 | Czech |
| 123 | IP-Vdt-0.9595 | Spain | 358 | Np_0 | Germany | 593 | Pu2_7 | Czech |
| 124 | IP-Ver-5.9596 | Spain | 359 | Nw_0 | Germany | 594 | Jm_0 | Czech |
| 125 | IP-Vig-1.9597 | Spain | 360 | Ob_0 | Germany | 595 | ICE1.MPI | Romania |
| 126 | IP-Vim-0.9598 | Spain | 361 | Old_1 | Germany | 596 | ICE7.MPI | Romania |
| 127 | IP-Vin-0.9599 | Spain | 362 | Ove_0 | Germany | 597 | Iasi-1.9744 | Romania |
| 128 | IP-Vis-0.9600 | Spain | 363 | Pt_0 | Germany | 598 | Teiu-2.9736 | Romania |
| 129 | IP-Voz-0.9601 | Spain | 364 | Pt_0 | Germany | 599 | Toc-1.9739 | Romania |
| 130 | IP-Vpa-1.9602 | Spain | 365 | Rd_0 | Germany | 600 | Ulies-1.9737 | Romania |
| 131 | LL_0 | Spain | 366 | Sg_1 | Germany | 601 | Litva | Lithuania |
| 132 | Sf-2.WTC | Spain | 367 | Sp_0 | Germany | 602 | Kn-0.WTC | Lithuania |
| 133 | Sf_1 | Spain | 368 | Star-8.MPI | Germany | 603 | Wil_1 | Lithuania |
| 134 | Ts_5 | Spain | 369 | TueSB30-3.MPI | Germany | 604 | Lag1-2.9100 | Georgia |
| 135 | Co | Portugal | 370 | Tuescha9.MPI | Germany | 605 | Lag1-4.9102 | Georgia |
| 136 | IP-Alo-0.9506 | Portugal | 371 | TueWa1-2.MPI | Germany | 606 | Lag1-6.9104 | Georgia |
| 137 | IP-Coa-0.9507 | Portugal | 372 | Uk_1 | Germany | 607 | Bak-2.MPI | Georgia |
| 138 | IP-Mos-1.9508 | Portugal | 373 | WalhaesB4.MPI | Germany | 608 | Bak-7.MPI | Georgia |
| 139 | IP-Rei-0.9510 | Portugal | 374 | Wc_1 | Germany | 609 | Lag2.2.MPI | Georgia |
| 140 | IP-Vav-0.9511 | Portugal | 375 | Wl_0 | Germany | 610 | Vash-1.MPI | Georgia |
| 141 | IP-Vid-1.9512 | Portugal | 376 | Wt_5 | Germany | 611 | En_D | Ukraine |
| 142 | Co_1 | Portugal | 377 | Anholt_1 | Germany | 612 | Kastel-1.MPI | Ukraine |
| 143 | Fei-0.MPI | Portugal | 378 | Do_0 | Germany | 613 | Rubeznhoe_1 | Ukraine |
| 144 | C24.MPI | Portugal | 379 | Ga_0 | Germany | 614 | Koch-1.MPI | Ukraine |
| 145 | 11C1.9503 | UK | 380 | Hh_0 | Germany | 615 | Istisu-1.MPI | Azerbaijan |
| 146 | CIBC_17 | UK | 381 | Rue3-1-31.MPI | Germany | 616 | Lerik1-3.MPI | Azerbaijan |
| 147 | CIBC_5 | UK | 382 | Mh_0 | Poland | 617 | Xan-1.MPI | Azerbaijan |
| 148 | Cnt-1.5726 | UK | 383 | Bs_1 | Switzerland | 618 | Adam-1.9609 | Russia |
| 149 | Edi-0.WTC | UK | 384 | Wei_0 | Switzerland | 619 | Balan-1.9613 | Russia |
| 150 | Gol-2.9314 | UK | 385 | Ge_0 | Switzerland | 620 | Basta-1.9619 | Russia |
| 151 | HR_10 | UK | 386 | Nd-1.CeBiT | Switzerland | 621 | Basta-2.9620 | Russia |
| 152 | Kent | UK | 387 | Zu-0.WTC | Switzerland | 622 | Bijisk-4.9622 | Russia |
| 153 | NFA_10 | UK | 388 | Zu-1.7418 | Switzerland | 623 | Chaba-2.9624 | Russia |
| 154 | NFA_8 | UK | 389 | Aiell-1.9646 | Italy | 624 | K-oze-1.9629 | Russia |
| 155 | Set-1.5772 | UK | 390 | Bivio-1.9649 | Italy | 625 | K-oze-3.9630 | Russia |
| 156 | Sq_1 | UK | 391 | Castelfed-1-197.9681 | Italy | 626 | Karag-1.9617 | Russia |
| 157 | Ty-1.5784 | UK | 392 | Castelfed-4-211.9695 | Italy | 627 | Karag-2.9608 | Russia |
| 158 | UKID107.5811 | UK | 393 | Castelfed-4-214.9696 | Italy | 628 | Kolyv-2.9625 | Russia |
| 159 | UKID114.5818 | UK | 394 | Cimin-1.9661 | Italy | 629 | Kolyv-3.9626 | Russia |
| 160 | UKID63.5768 | UK | 395 | Corig-1.9650 | Italy | 630 | Kolyv-5.9627 | Russia |
| 161 | UKID74.5779 | UK | 396 | Ct-1.WTC | Italy | 631 | Kolyv-6.9628 | Russia |
| 162 | UKID96.5800 | UK | 397 | Filet-1.9651 | Italy | 632 | Lebja-2.9632 | Russia |
| 163 | UKNW06-003.5353 | UK | 398 | Fondi-1.9652 | Italy | 633 | Lebja-4.9633 | Russia |
| 164 | UKNW06-403.5577 | UK | 399 | Giffo-1.9653 | Italy | 634 | Lesno-1.9611 | Russia |
| 165 | UKNW06-481.5644 | UK | 400 | Liri-1.9654 | Italy | 635 | Lesno-2.9612 | Russia |
| 166 | UKSE06-118.5023 | UK | 401 | Marce-1.9655 | Italy | 636 | Lesno-4.9610 | Russia |
| 167 | UKSE06-252.5104 | UK | 402 | Melic-1.9657 | Italy | 637 | Masl-1.9634 | Russia |
| 168 | UKSE06-325.5151 | UK | 403 | Mir_0 | Italy | 638 | N13 | Russia |
| 169 | UKSE06-362.5165 | UK | 404 | Mitterberg-1-180.9665 | Italy | 639 | Nosov-1.9635 | Russia |
| 170 | UKSE06-432.5210 | UK | 405 | Mitterberg-1-182.9666 | Italy | 640 | Noveg-1.9636 | Russia |
| 171 | UKSE06-470.5236 | UK | 406 | Mitterberg-1-183.9667 | Italy | 641 | Noveg-2.9637 | Russia |
| 172 | UKSE06-500.5253 | UK | 407 | Mitterberg-2-184.9668 | Italy | 642 | Noveg-3.9638 | Russia |
| 173 | UKSE06-533.5276 | UK | 408 | Mitterberg-2-185.9669 | Italy | 643 | Panik-1.9607 | Russia |
| 174 | UKSW06-179.4779 | UK | 409 | Mitterberg-3-187.9671 | Italy | 644 | Panke-1.9639 | Russia |
| 175 | UKSW06-207.4807 | UK | 410 | Nicas-1.9658 | Italy | 645 | Parti-1.9615 | Russia |
| 176 | UKSW06-226.4826 | UK | 411 | PHW_2 | Italy | 646 | Petergof.7296 | Russia |
| 177 | UKSW06-285.4884 | UK | 412 | Pigna-1.9659 | Italy | 647 | Rakit-1.9640 | Russia |
| 178 | UKSW06-302.4900 | UK | 413 | Sarno-1.9660 | Italy | 648 | Rakit-3.9642 | Russia |
| 179 | UKSW06-333.4931 | UK | 414 | Stilo-1.9662 | Italy | 649 | Sever-1.9643 | Russia |
| 180 | UKSW06-360.4958 | UK | 415 | Teano-1.9663 | Italy | 650 | Ws-0.WTC | Russia |
| 181 | Ullapool-8.9312 | UK | 416 | Etna_2 | Italy | 651 | Est-1.MPI | Russia |
| 182 | Abd_0 | UK | 417 | ICE102.MPI | Italy | 652 | ICE130.MPI | Russia |
| 183 | Alst_1 | UK | 418 | ICE104.MPI | Italy | 653 | ICE134.MPI | Russia |
| 184 | Ba_1 | UK | 419 | ICE106.MPI | Italy | 654 | ICE138.MPI | Russia |
| 185 | Boot_1 | UK | 420 | ICE107.MPI | Italy | 655 | ICE70.MPI | Russia |
| 186 | Cal_0 | UK | 421 | ICE111.MPI | Italy | 656 | ICE71.MPI | Russia |
| 187 | Durh_1 | UK | 422 | ICE112.MPI | Italy | 657 | Ms_0 | Russia |
| 188 | Ema_1 | UK | 423 | ICE119.MPI | Italy | 658 | Per_1 | Russia |
| 189 | HR_5 | UK | 424 | ICE120.MPI | Italy | 659 | Rld_1 | Russia |
| 190 | Mc_0 | UK | 425 | ICE163.MPI | Italy | 660 | Stw_0 | Russia |
| 191 | Sq_8 | UK | 426 | ICE169.MPI | Italy | 661 | Kondara | Tadjikistan |
| 192 | Su_0 | UK | 427 | ICE173.MPI | Italy | 662 | Sorbo | Tadjikistan |
| 193 | Ty_0 | UK | 428 | ICE181.MPI | Italy | 663 | Neo_6 | TJK |
| 194 | Vind_1 | UK | 429 | ICE212.MPI | Italy | 664 | Dja_1 | Kyrgyzstan |
| 195 | Bur-0.MPI | Ireland | 430 | ICE216.MPI | Italy | 665 | Kar_1 | Kyrgyzstan |
| 196 | ARGE-1-15.9911 | France | 431 | ICE228.MPI | Italy | 666 | Sus_1 | Kyrgyzstan |
| 197 | ARR-17.9927 | France | 432 | ICE79.MPI | Italy | 667 | Westkar_4 | Kyrgyzstan |
| 198 | BEZ-9.9928 | France | 433 | ICE91.MPI | Italy | 668 | Epidauros-1.9725 | Greece |
| 199 | BRE-14.9919 | France | 434 | ICE92.MPI | Italy | 669 | Faneronemi-3.9726 | Greece |
| 200 | BRI-2.9910 | France | 435 | ICE93.MPI | Italy | 670 | Olympia-2.9727 | Greece |
| 201 | CATS-6.9937 | France | 436 | ICE97.MPI | Italy | 671 | ICE36.MPI | Serbia |
| 202 | CON-7.9913 | France | 437 | ICE98.MPI | Italy | 672 | Knjas-1.9749 | Serbia |
| 203 | CYR.88 | France | 438 | Rome_1 | Italy | 673 | Malii-1.9746 | Serbia |
| 204 | DIR-9.9920 | France | 439 | Sei_0 | Italy | 674 | Staro-1.9757 | Serbia |
| 205 | ESP-1-11.9908 | France | 440 | Tu_0 | Italy | 675 | Zagub-1.9748 | Serbia |
| 206 | GEN-8.9909 | France | 441 | Oy-0.JGI | Norway | 676 | Bela-1.9730 | Slovakia |
| 207 | ISS-20.9929 | France | 442 | Oy-0.WTC | Norway | 677 | Bela-2.9733 | Slovakia |
| 208 | IST-29.9914 | France | 443 | Ale-Stenar-44-4 | Sweden | 678 | Stiav-1.9728 | Slovakia |
| 209 | Jea.JGI | France | 444 | Ale-Stenar-64-24 | Sweden | 679 | Dog-4.MPI | Turkey |
| 210 | LDV-18.108 | France | 445 | Algutsrum | Sweden | 680 | Nemrut-1.MPI | Turkey |
| 211 | LDV-46.139 | France | 446 | App1-12 | Sweden | 681 | ICE33.MPI | Bulgaria |
| 212 | LEC-25.9930 | France | 447 | App1-14 | Sweden | 682 | Gradi-1.9645 | Croatia |
| 213 | MAR-4-16.9915 | France | 448 | App1-16 | Sweden | 683 | Sakata.JGI | JPN |
| 214 | MAR2-3.159 | France | 449 | Baa1-2 | Sweden | 684 | Bik_1 | Lebanon |
| 215 | MIL-2.9922 | France | 450 | Bil-5 | Sweden | 685 | Altai_5 | Mongolia |
| 216 | MOL-1.9916 | France | 451 | Bil-7 | Sweden | 686 | Kas_2 | India |
| 217 | MOU2-25.9931 | France | 452 | Boo2-1.8266 | Sweden | 687 | Kas_1 | India |
| 218 | Na_1 | France | 453 | Broesarp-34-145 | Sweden | 688 | ICE150.MPI | Uzbekistan |
| 219 | NOZ-6.9932 | France | 454 | Dra2-1 | Sweden | 689 | ICE152.MPI | Uzbekistan |
| 220 | PHW_34 | France | 455 | Fjae1-1 | Sweden | 690 | ICE153.MPI | Uzbekistan |
| 221 | PLO-1.9923 | France | 456 | Fjae1-2 | Sweden | 691 | Geg-14.9125 | Armenia |
| 222 | PLY-20.9924 | France | 457 | Fjae1-5 | Sweden | 692 | Yeg-2.9128 | Armenia |
| 223 | PYL-6.265 | France | 458 | Fjae2-4 | Sweden | 693 | Yeg-4.9130 | Armenia |
| 224 | QUI-8.9934 | France | 459 | Fly2-1 | Sweden | 694 | Yeg-5.9131 | Armenia |
| 225 | RAD-21.9917 | France | 460 | Fly2-2 | Sweden | 695 | Yeg-7.9133 | Armenia |
| 226 | Ren_1 | France | 461 | Groen-12 | Sweden | 696 | Yeg-8.9134 | Armenia |
| 227 | Ren_11 | France | 462 | Had-1 | Sweden | 697 | Cnt-1.5726 | unknown |
| 228 | RUM-20.9925 | France | 463 | Had-2 | Sweden | 698 | IP-Orb-10.9565 | unknown |
| 229 | SAUL-24.9918 | France | 464 | Ham-1 | Sweden | 699 | ICE49.MPI | Morocco |
| 230 | TOU-A1-88.350 | France | 465 | Hel-3 | Sweden | 700 | ICE50.MPI | Morocco |
| 231 | TOU-A1-89.351 | France | 466 | Hov1-7 | Sweden | 701 | Aitba-1.9606 | Morocco |
| 232 | TRE-1.9926 | France | 467 | Hovdala-2 | Sweden | 702 | Anz_0 | Tanzania |
| 233 | VED-10.9933 | France | 468 | Kaevlinge-1 | Sweden | 703 | Cvi-0 | Cape Verdi |
| 234 | WAV-8.9938 | France | 469 | Kal-2 | Sweden | 704 | Cvi_0 | Cape Verde |
| 235 | Ann_1 | France | 470 | Kia-1 | Sweden |  |  |  |

**Table S5. The substitution and polymorphism data for the McDonald-Kreitman test**

**A. Substitution sites:**

*A. lyrata*

*A. thaliana*

*Exov-l*

*Exov*

C

C

T

C

C

C🡪T

The substitutions are inferred from the parsimonious method taking *A. lyrate* as outgroup. The example below shows the divergence between *Exov-l* and *Exov* at site 120 as C and T, respectively. Thus, substitution at this site is parsimoniously inferred as C🡪T in the *Exov* lineage (red) because the *Exov-l* in *A. thaliana* and its orthologue in the outgroup *A. lyrata* are C at the site, which indicate that the two ancestral states are C.

**Synonymous substitutions:**

Exov: 33G (x0.5), 120T, 207C, 210T, 225G, 252A, 276T, 339A, 348G, 375T,402C(x0.5), 408A, 198A. Sum: 12

Exov-L: 249T, 255A, 264C, 375A. Sum: 4.

**Nonsynonymous substitutions:**

Exov: 32C(x1.5), 35T, 139A, 143A, 158T, 188A, 230A,254T, 305T, 307T, 349T, 380T, 385T, 395T, 398C, 400G, 401T (x1.5), 404T, 156A, 234G, 255A . Sum: 22.

Exov-L: 184T, 206T, 236C. Sum: 3.

**B. Polymorphisms:**

Synonymous polymorphisms (gene/accession numbers, proportion for nucleotide)

| **Sites** | **117** | **183** | **225** | **282** | **312** | **339** | **348** | **351** | **360** | **363** | **375** | **411** |
| --- | --- | --- | --- | --- | --- | --- | --- | --- | --- | --- | --- | --- |
| Exov  (709) | 0.990C  0.010T | 0.997C  0.003T |  | 0.993C  0.007T | 0.032A  0.968C |  |  | 0.999T  0.001C | 0.993C  0.007T | 0.994A  0.006G | 0.024C  0.976T | 0.001A  0.999G |
| Exov-l  (455)  Nonsynonymous polymorphisms (gene/accession numbers, proportion for nucleotide) |  |  | 0.154A  0.846G |  |  | 0.402A  0.598G | 0.613A  0.387G |  |  |  | 0.730A  0.270C |  |

| **Sites** | **64** | **88** | **139** | **143** | **163** | **202** | **205** | **296** | **307** | **316** |
| --- | --- | --- | --- | --- | --- | --- | --- | --- | --- | --- |
| Exov  (709) |  |  |  | 0.863A  0.137G |  |  |  |  | 0.103C  0.897T | 0.102A  0.898G |
| Exov-l  (455) | 0.035A  0.965G | 0.026G  0.974T | 0.515A  0.485G |  | 0.967C  0.033A | 0.532A  0.468T | 0.035A  0.965G | 0.974C  0.026G |  |  |

**Table S6a. GO enrichment of analysis of the set of genes that significantly differentially expressed between *exov* and *exov-l.***

| Biosynthesis of secondary metabolites | 40 | 6.6 | 5.70E-05 | 1.90E-03 |
| --- | --- | --- | --- | --- |
| 2-Oxocarboxylic acid metabolism | 7 | 1.1 | 3.60E-03 | 4.80E-02 |
| Plant hormone signal transduction | 12 | 2 | 1.70E-02 | 1.60E-01 |

**Table S6b. The significantly differentially expressed genes between *exov* and *exov-l***

**Table S6c. The significantly differentially expressed genes between *exov^crp^* and *exov-l^crp^***

See the excel document.

**Table S7. Pairwise comparisons for phenotypic traits of T-DNA mutants and CRISPR-Cas9 mutants (Wilcoxon rank sum test)**

A. T-DNA instertions: WT= wild type, *exov*: new gene T-DNA insertion line, *exov-l*: old gene T-DNA insertion line, *dm*= double mutant.

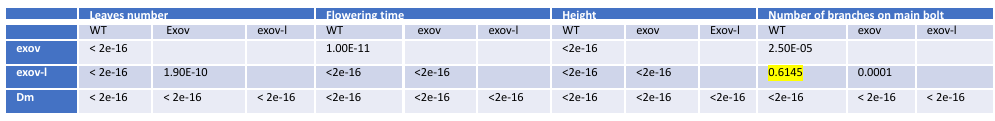

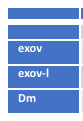

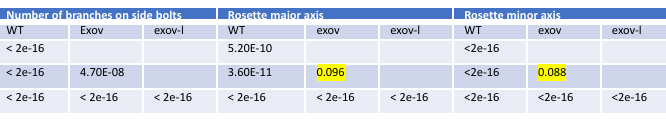

B. WT= wild type, *exol^crp^*: new gene CRISPR-Cas9 line, *exol-v^crp^*: old gene CRISPR-Cas9 insertion line

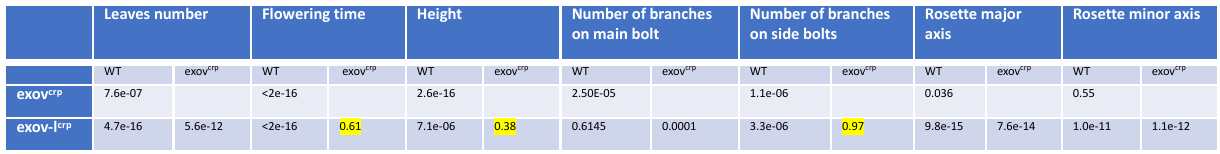

**Table S8a. GO enrichment of analysis of the set of genes that significantly differentially expressed between wild type and *exov-l*.**

| GO Term | Count | % | P-Value | Benjamini |
| --- | --- | --- | --- | --- |
| Biosynthesis of secondary metabolites | 39 | 8.5 | 3.80E-07 | 2.50E-05 |
| Phenylpropanoid biosynthesis | 14 | 3.1 | 1.30E-06 | 4.20E-05 |
| Metabolic pathways | 45 | 9.8 | 2.40E-03 | 5.10E-02 |
| Phenylalanine metabolism | 5 | 1.1 | 4.80E-03 | 7.60E-02 |
| Glucosinolate biosynthesis | 3 | 0.7 | 1.60E-02 | 1.90E-01 |
| 2-Oxocarboxylic acid metabolism | 5 | 1.1 | 3.30E-02 | 3.10E-01 |
| Cutin, suberine and wax biosynthesis | 3 | 0.7 | 7.20E-02 | 5.10E-01 |
| Cyanoamino acid metabolism | 4 | 0.9 | 7.60E-02 | 4.80E-01 |
| Stilbenoid, diarylheptanoid and gingerol biosynthesis | 4 | 0.9 | 7.90E-02 | 4.50E-01 |
| Indole alkaloid biosynthesis | 2 | 0.4 | 9.50E-02 | 4.80E-01 |

**Table S8b. GO enrichment analysis of the set of genes that significantly differentially expressed between wild type and *exov*.**

| GO Term | Count | % | P-Value | Benjamini |
| --- | --- | --- | --- | --- |
| Pentose and glucuronate interconversions | 8 | 1.3 | 1.20E-03 | 2.60E-02 |
| Alanine, aspartate and glutamate metabolism | 4 | 0.7 | 7.30E-02 | 4.00E-01 |
| Valine, leucine and isoleucine biosynthesis | 3 | 0.5 | 7.80E-02 | 3.90E-01 |
| Phenylalanine metabolism | 6 | 1 | 1.50E-03 | 2.40E-02 |
| Nitrogen metabolism | 4 | 0.7 | 5.30E-02 | 3.30E-01 |
| Phenylpropanoid biosynthesis | 16 | 2.6 | 4.30E-07 | 2.90E-05 |
| Glucosinolate biosynthesis | 3 | 0.5 | 2.40E-02 | 1.80E-01 |
| Metabolic pathways | 52 | 8.5 | 5.60E-03 | 6.10E-02 |

**Supplementary Figures:**

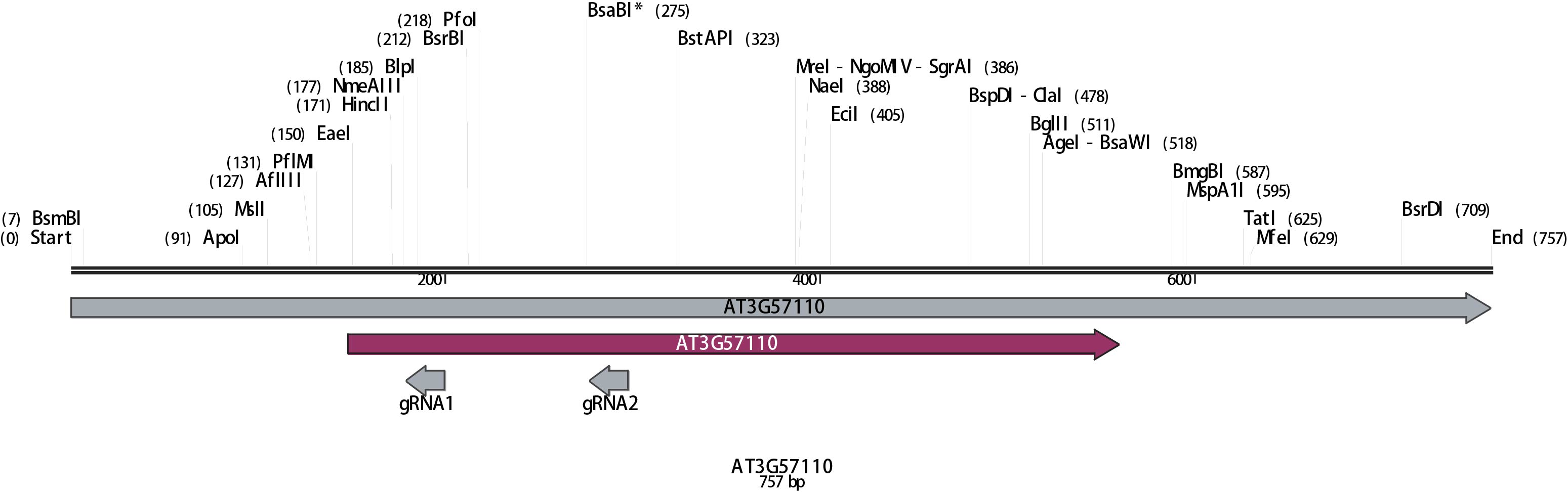

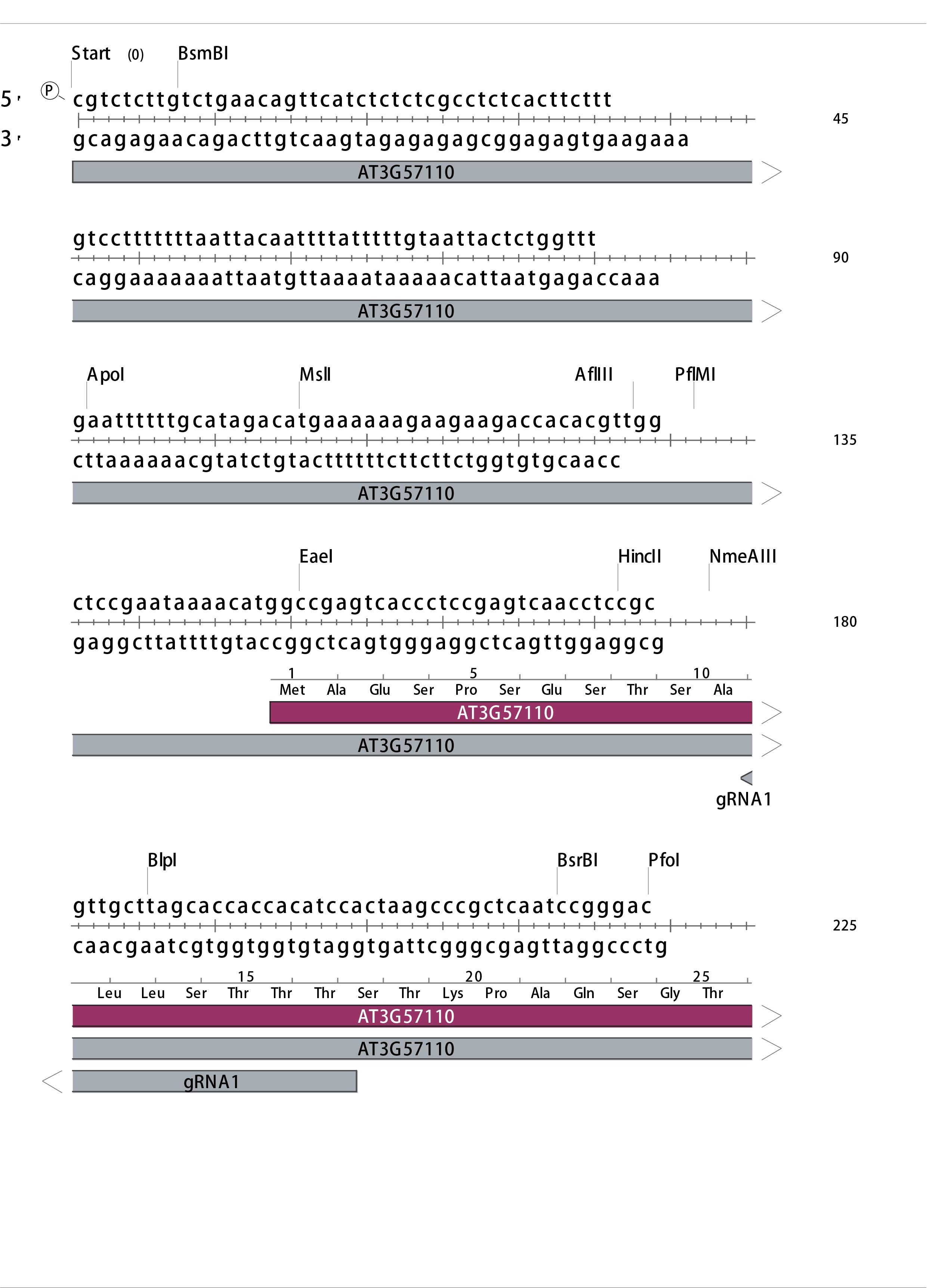

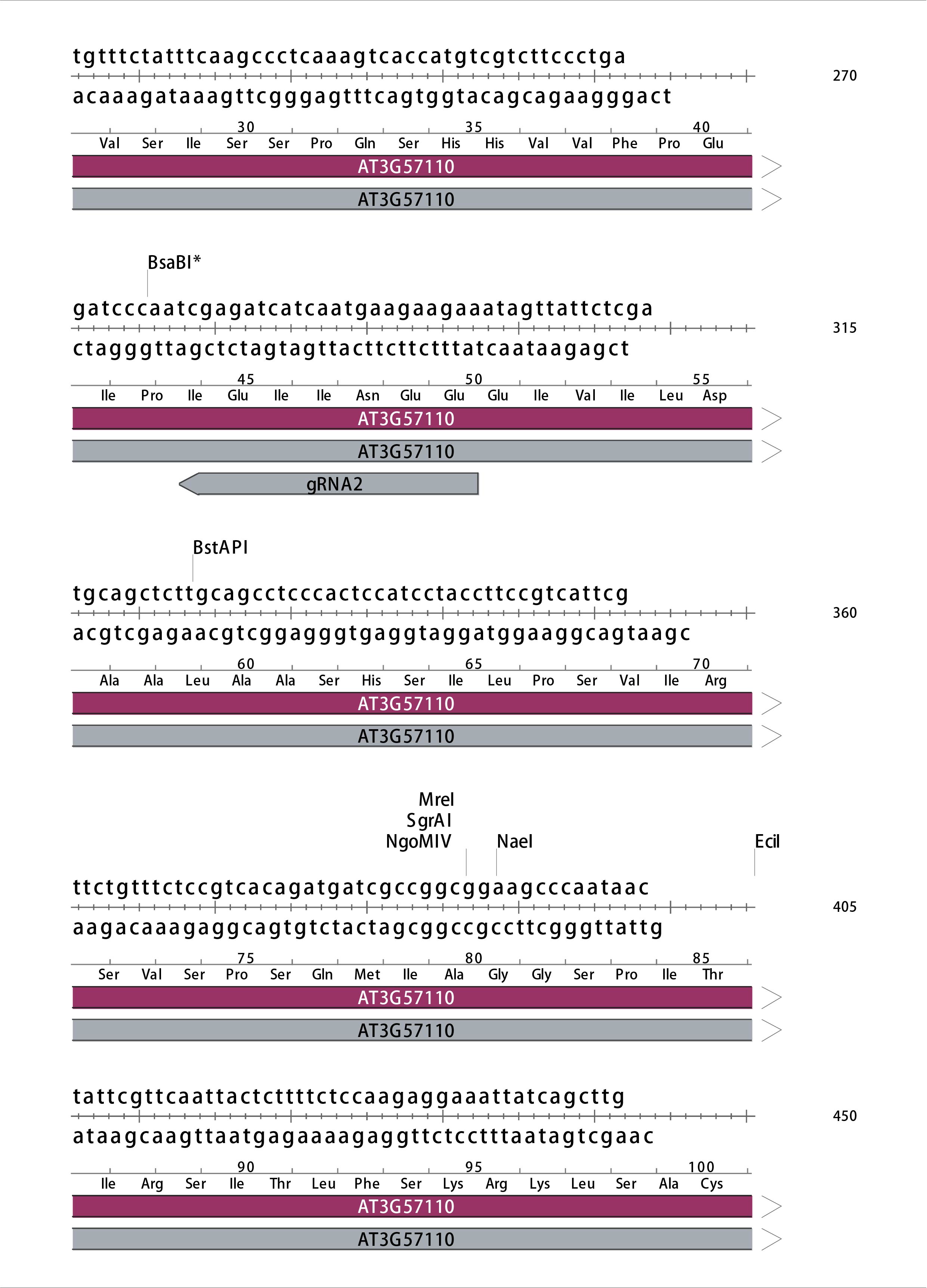

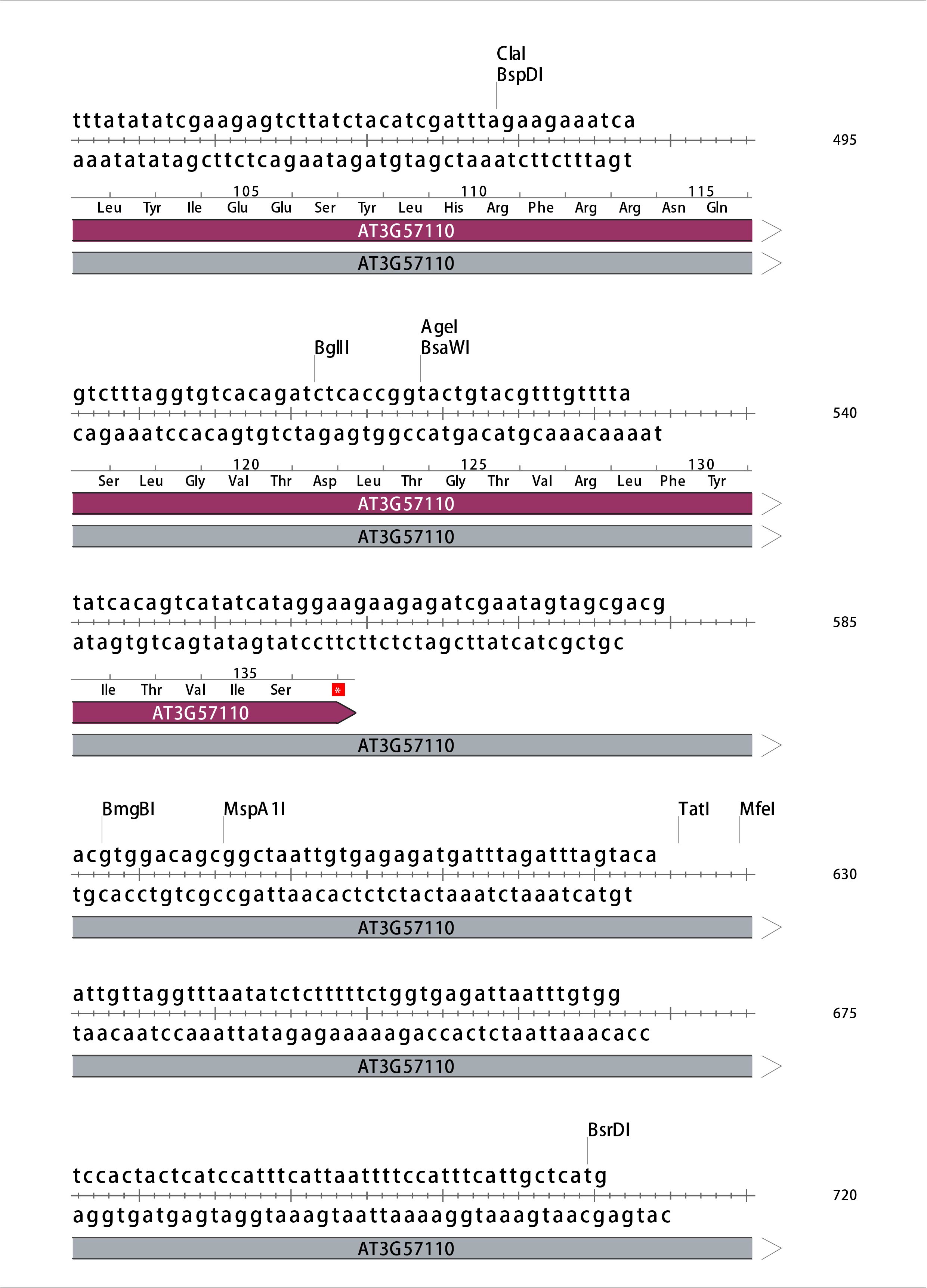

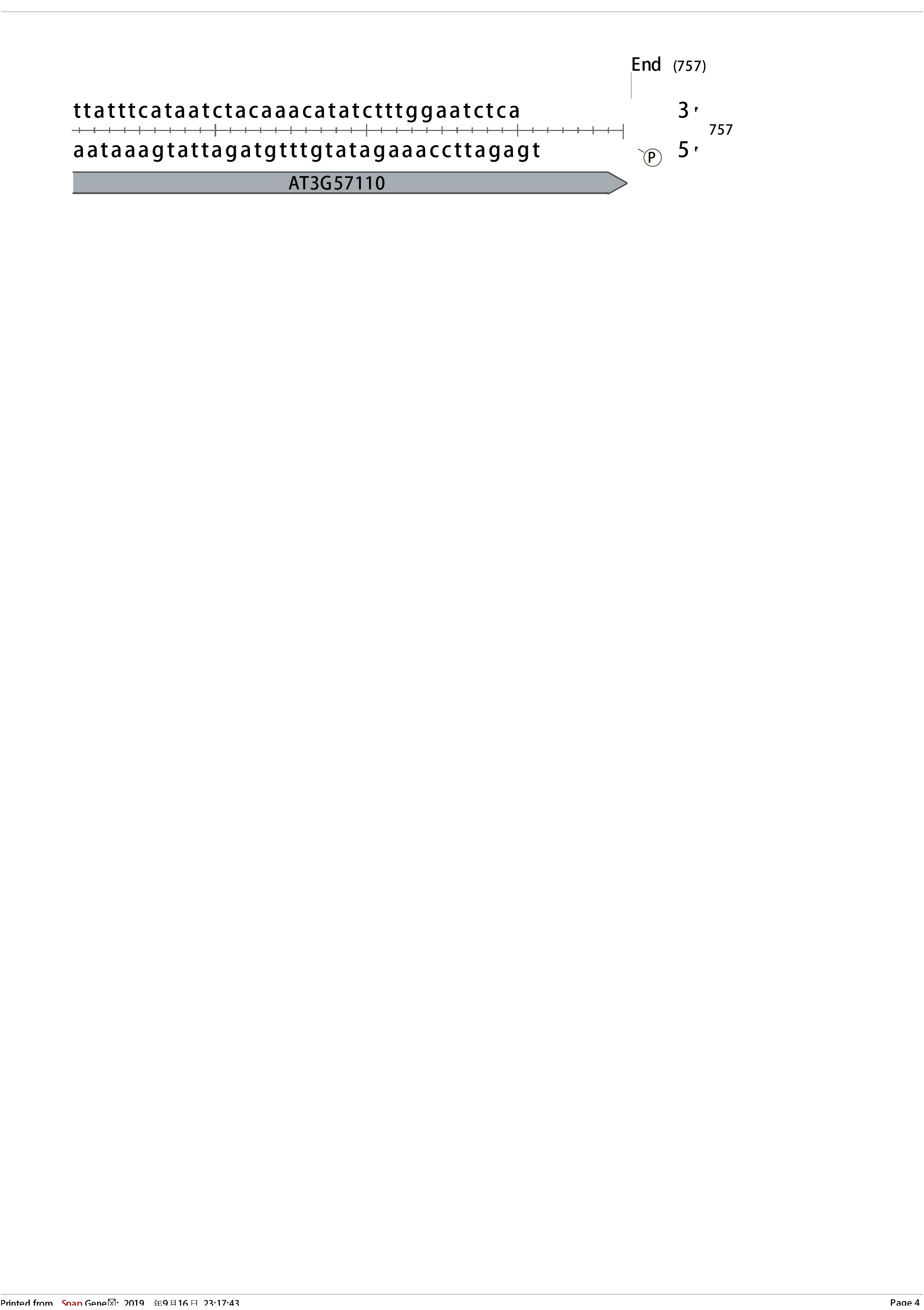

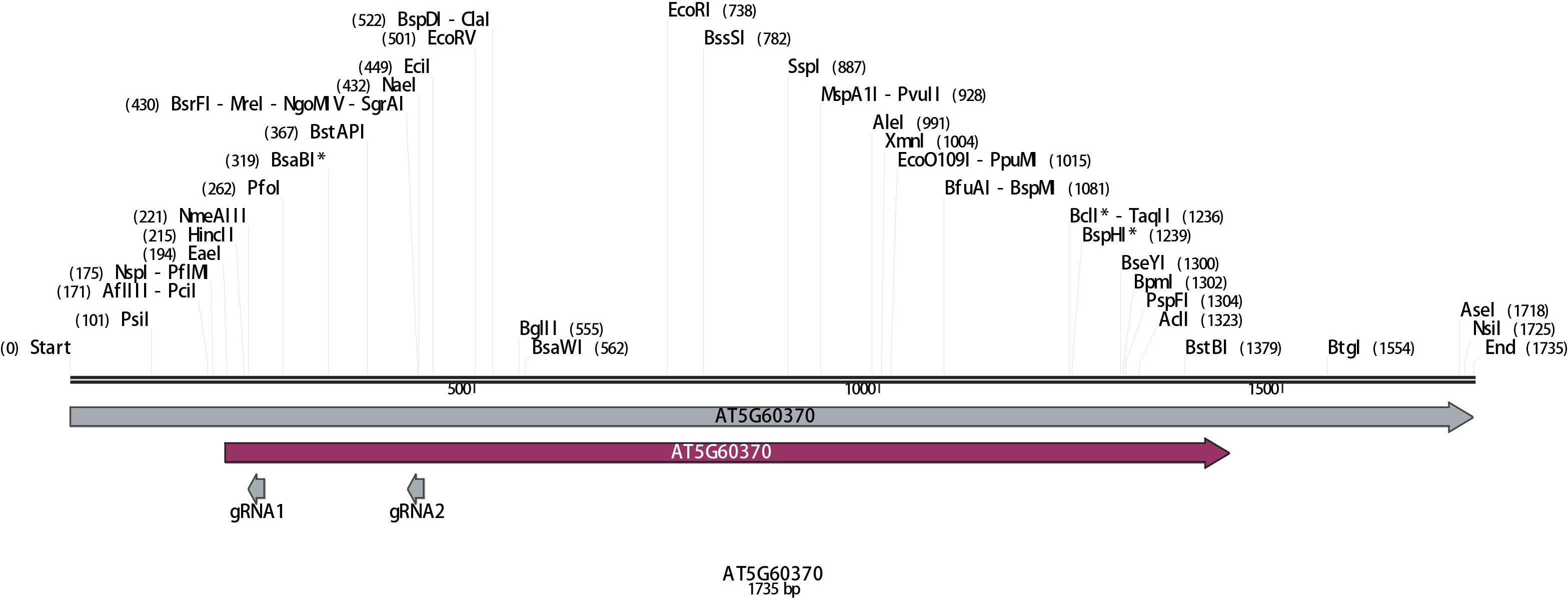

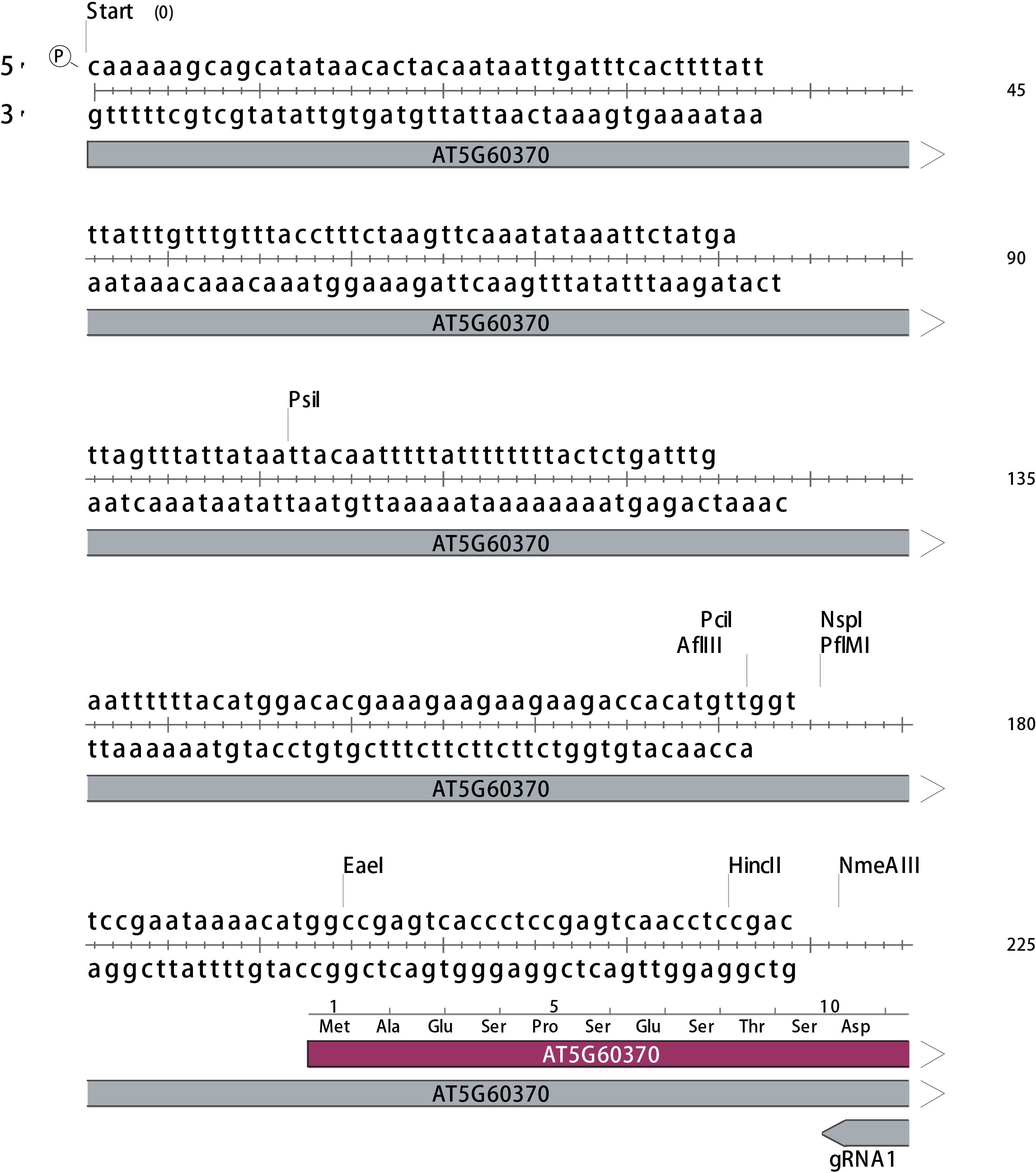

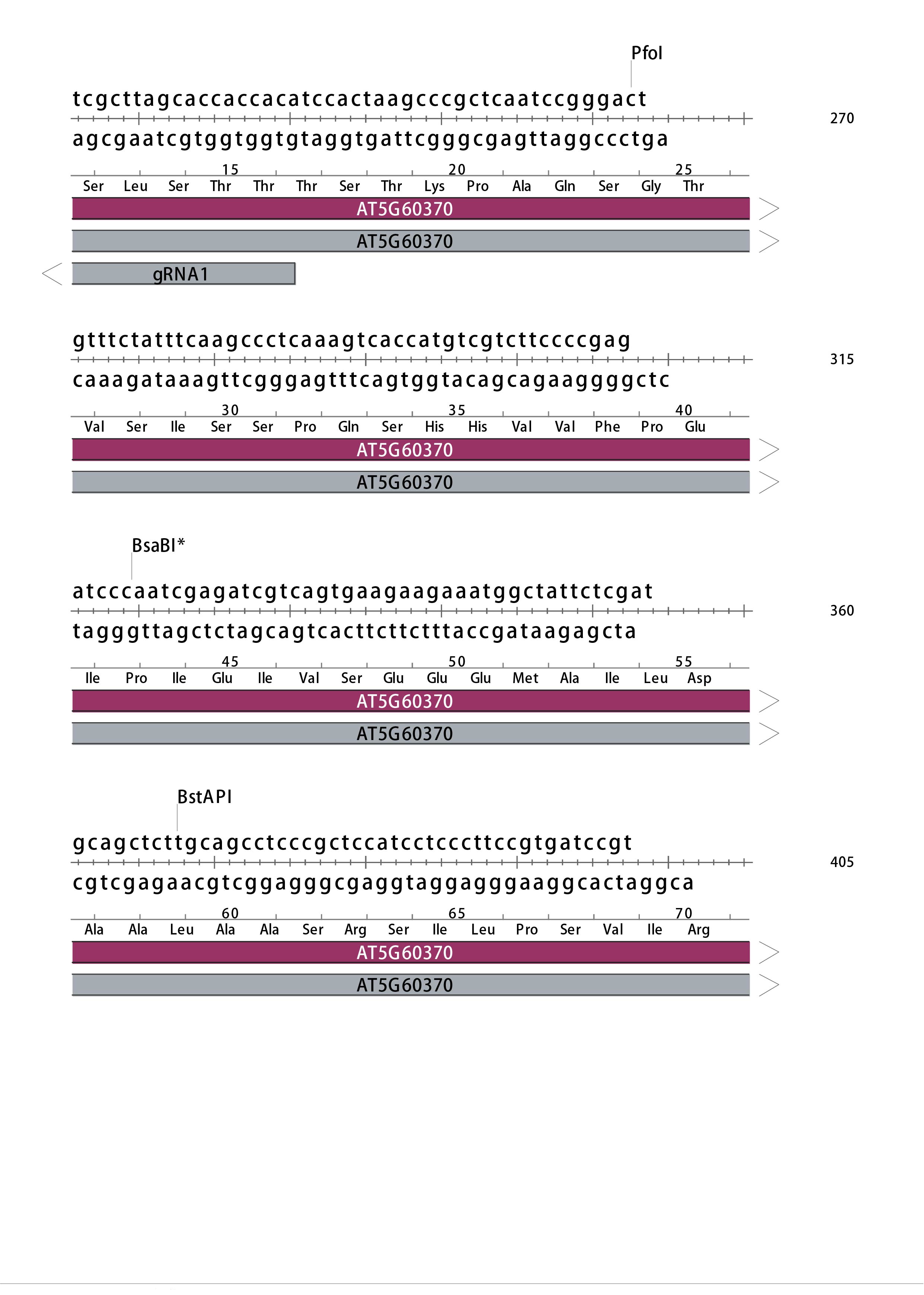

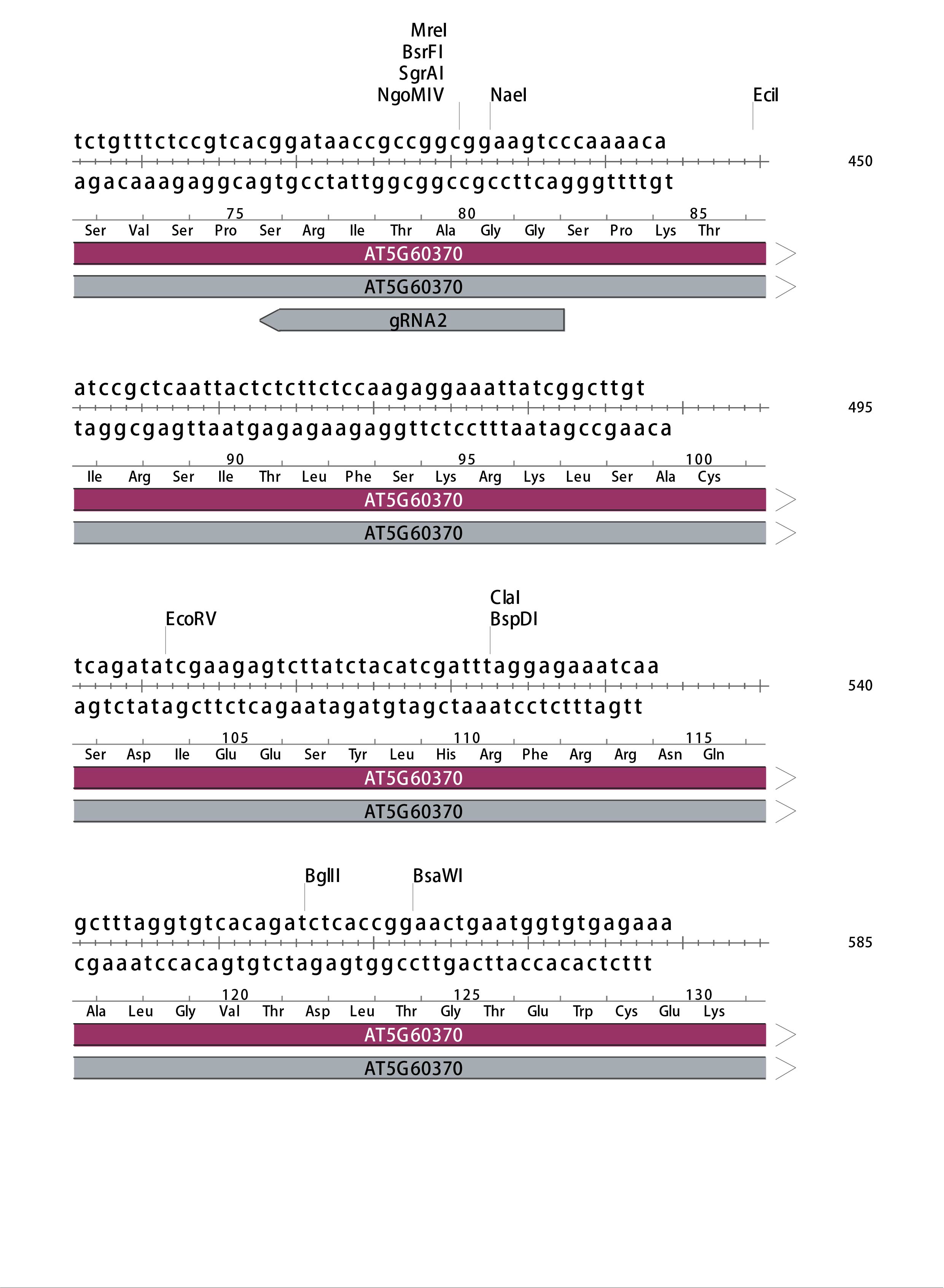

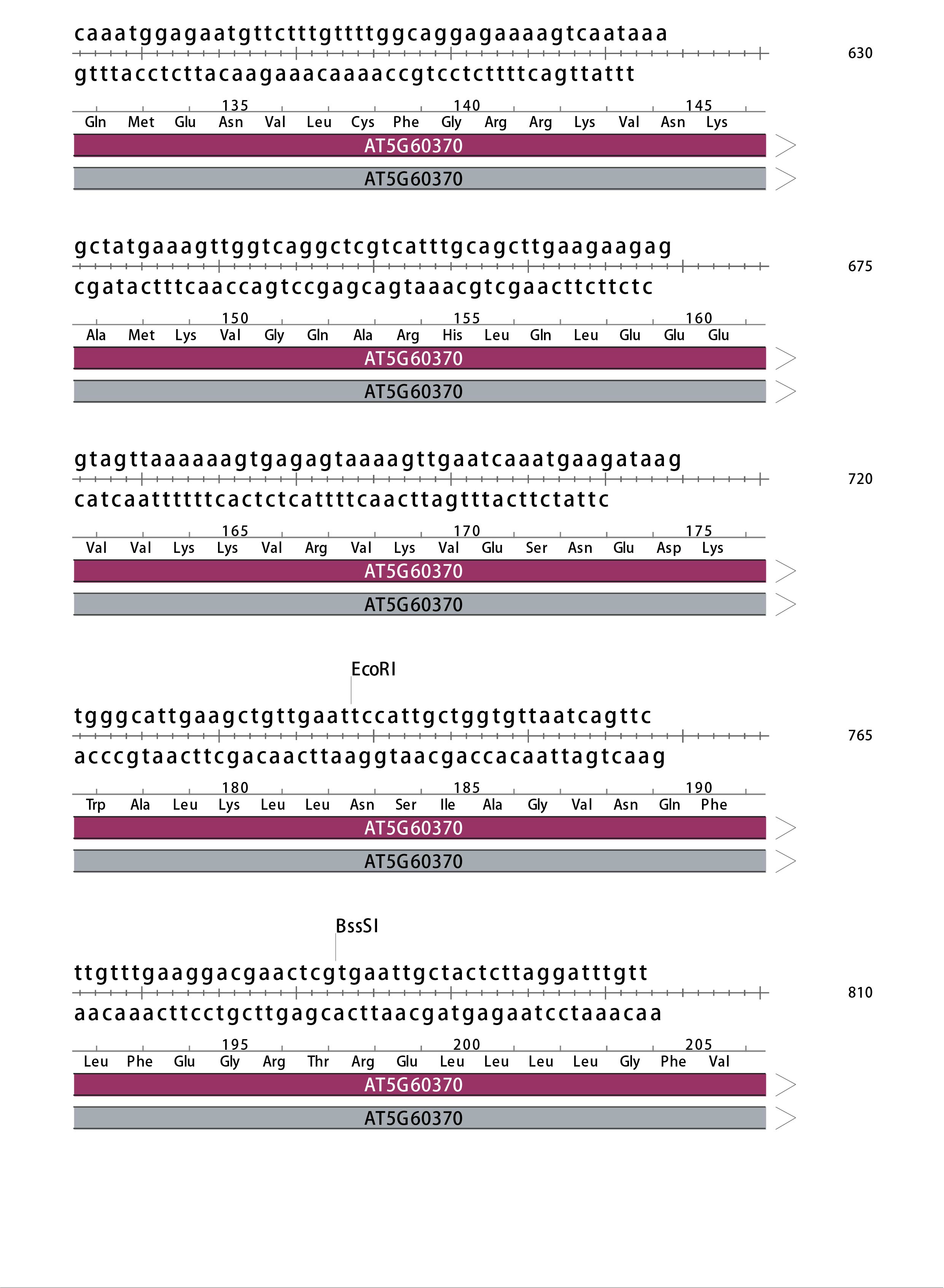

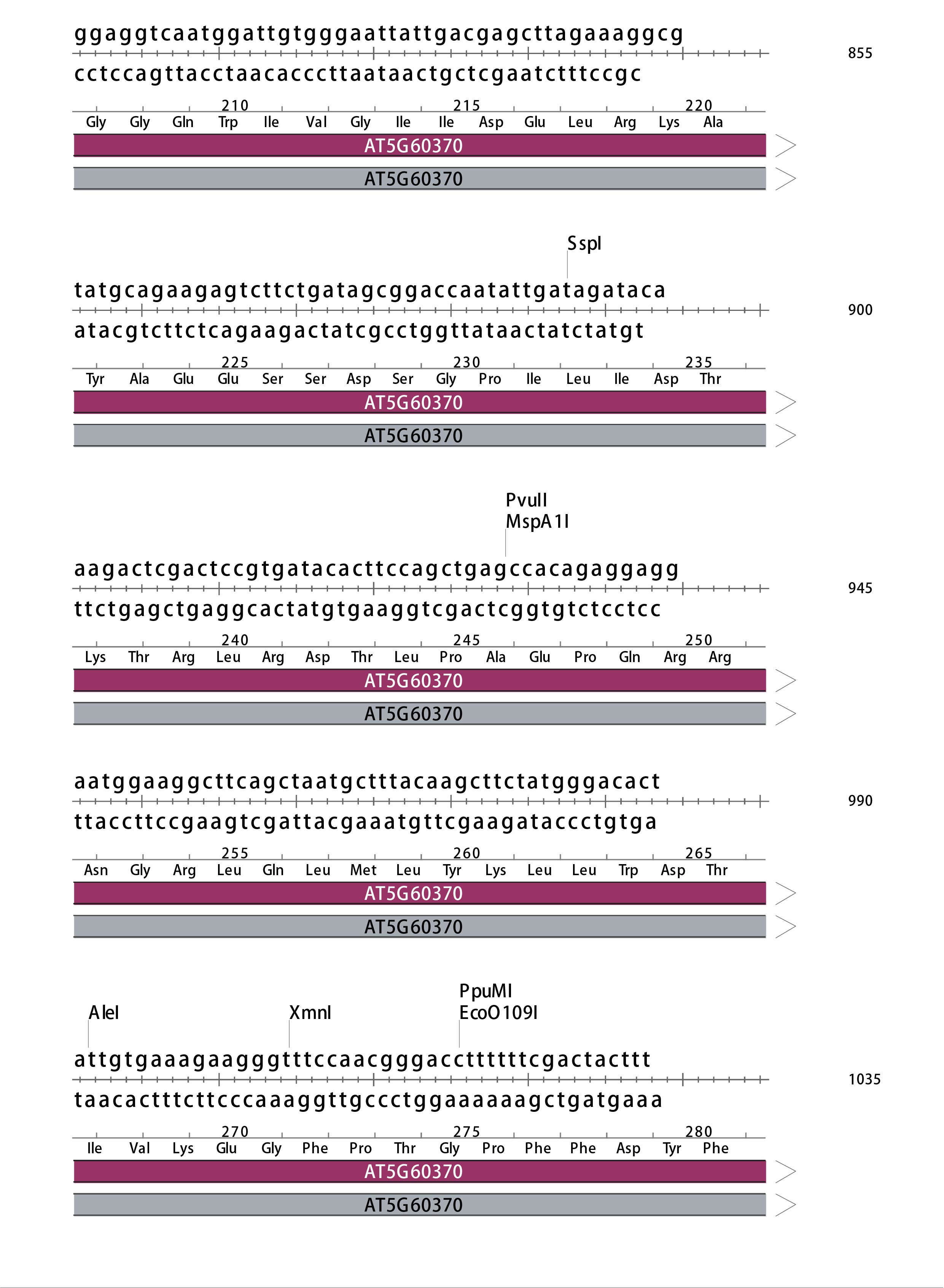

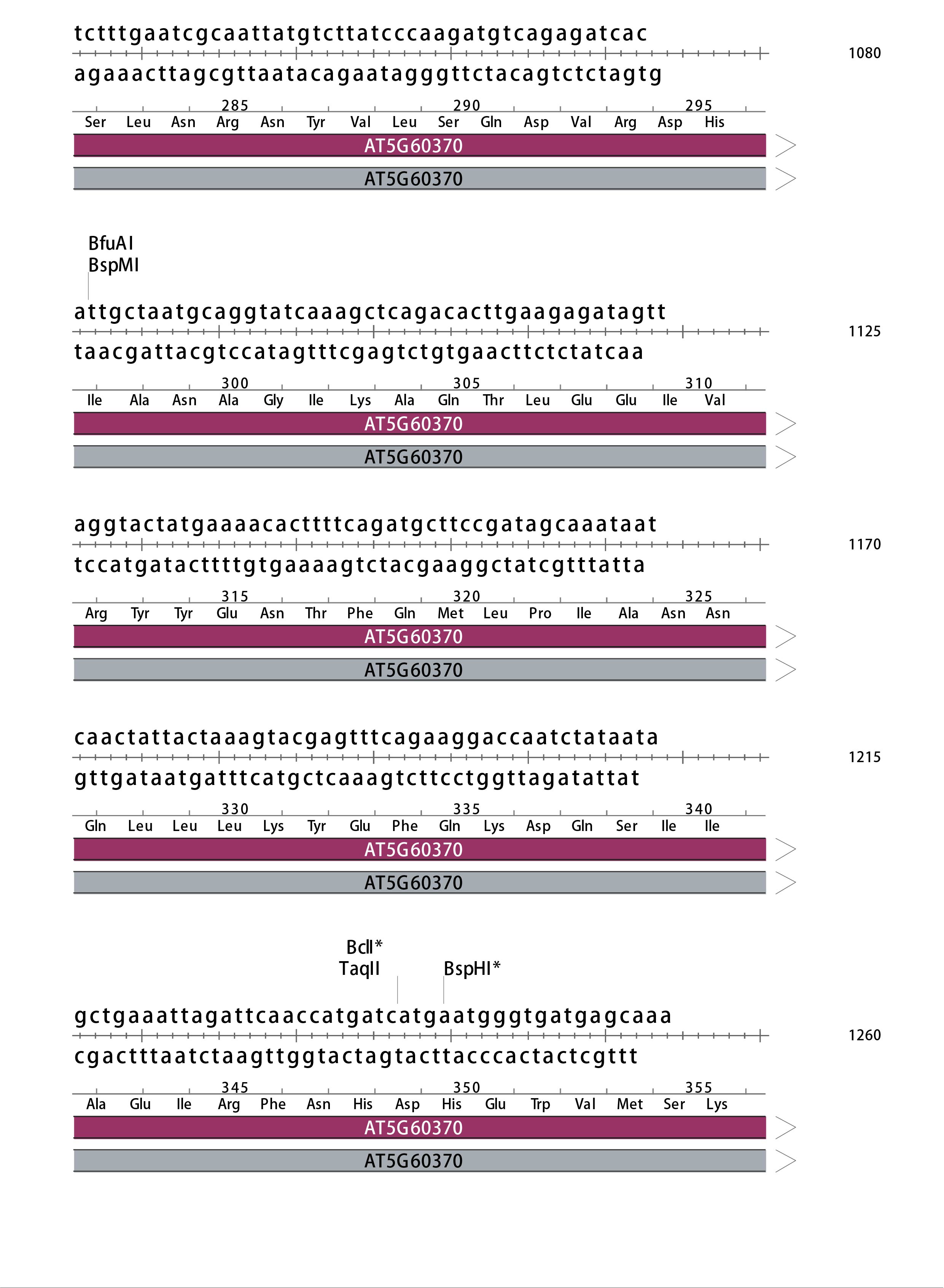

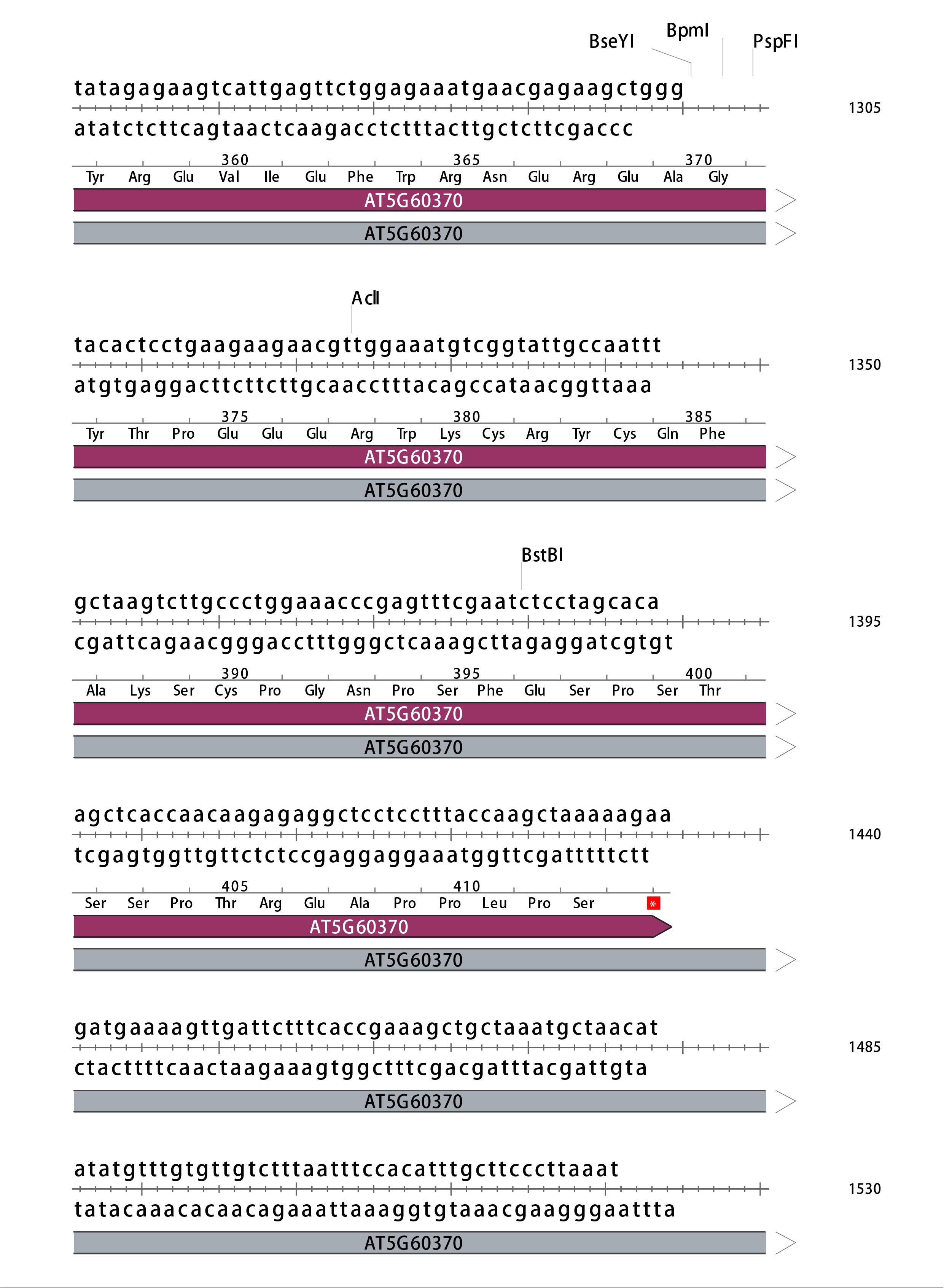

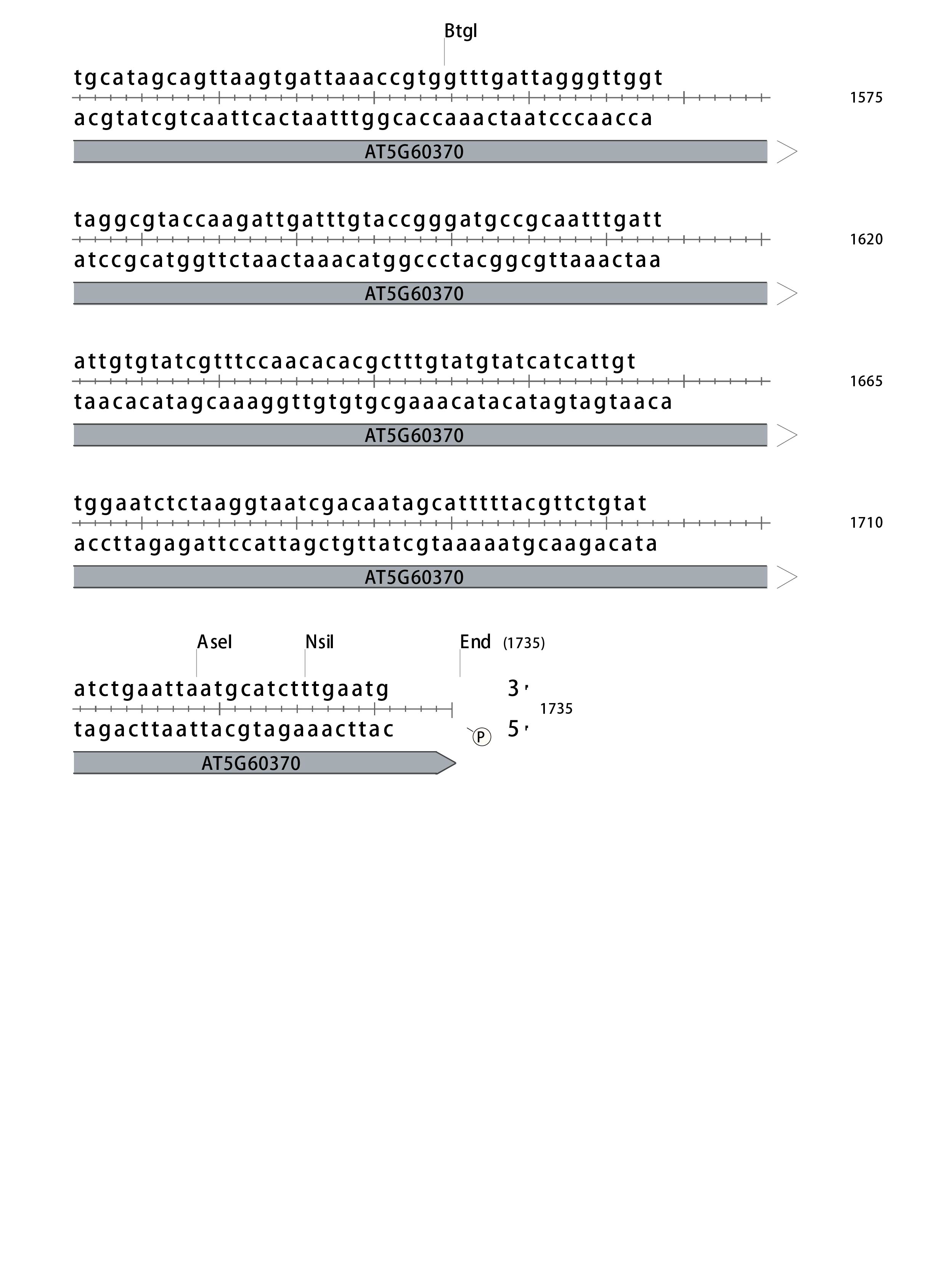

**Figure S1.** Sequences, annotations and restriction maps of AT3G57110 (Exov) and AT5G60370 (Exov-L) in the CRISPR/Cas9 vector pCAMBIA1300.

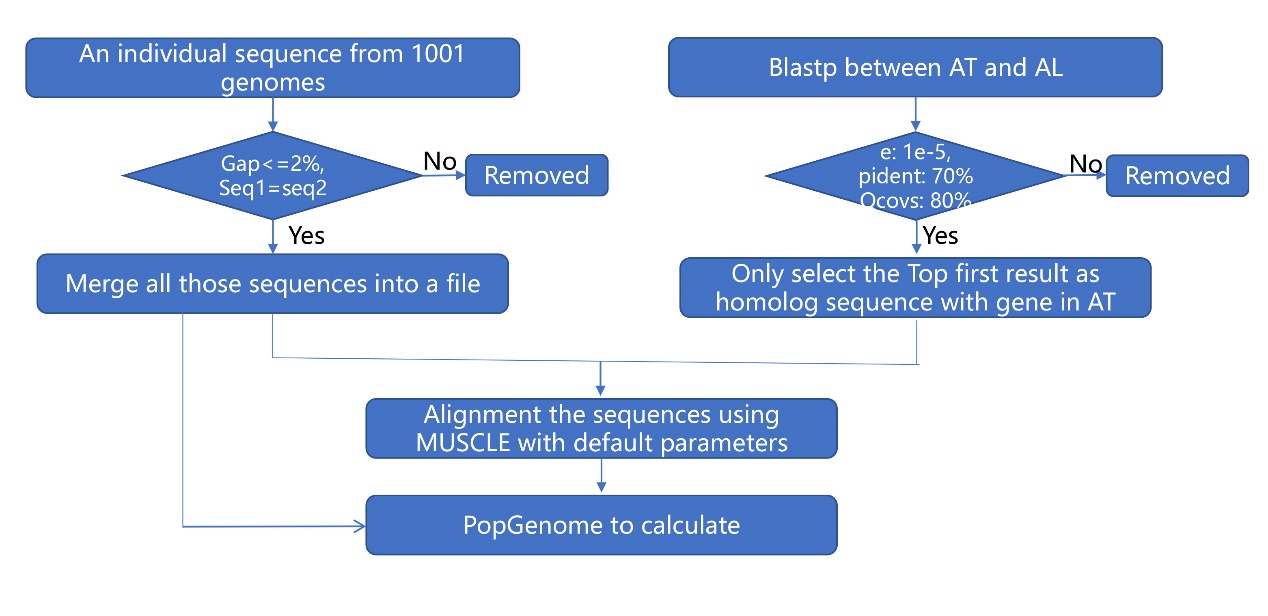

**Figure S2.** Summary of neutrality test pipeline.

**a.**

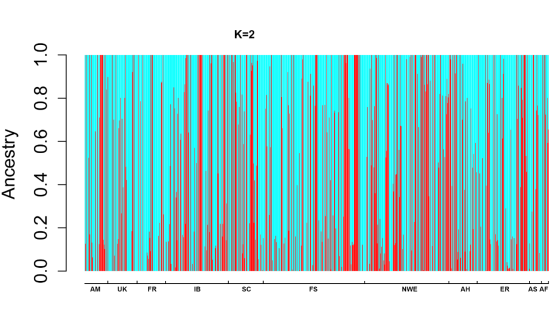

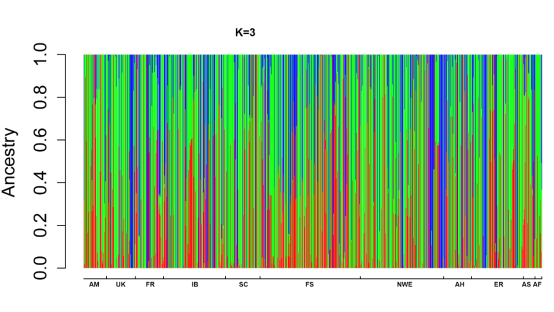

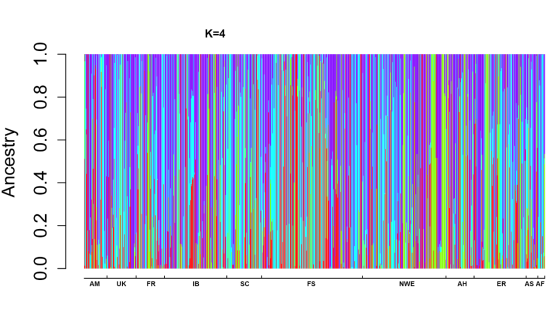

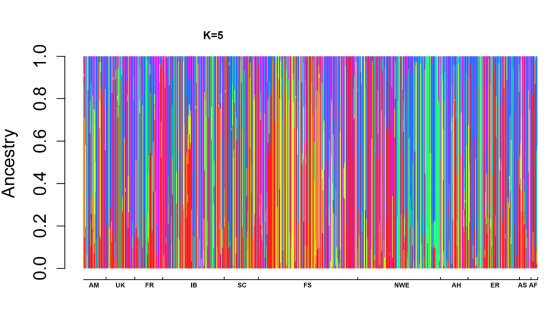

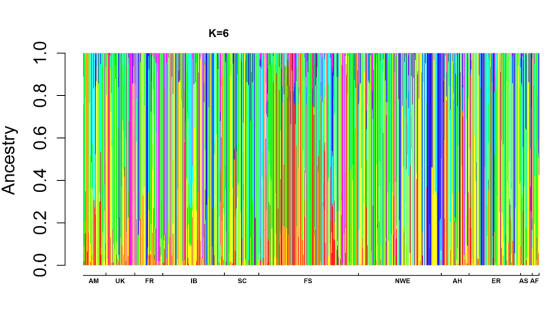

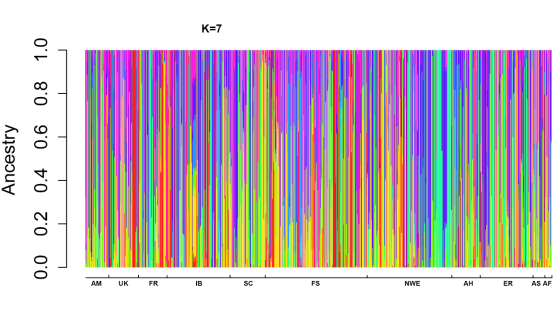

**b.**

Cross-validation error

| K | cross-validation error |
| --- | --- |
| 1 | 0.15445 |
| 2 | 0.13719 |
| 3 | 0.12977 |
| 4 | 0.11955 |
| 5 | 0.1121 |
| 6 | 0.10727 |
| 7 | 0.10323 |
| 8 | 0.10223 |

**c.**

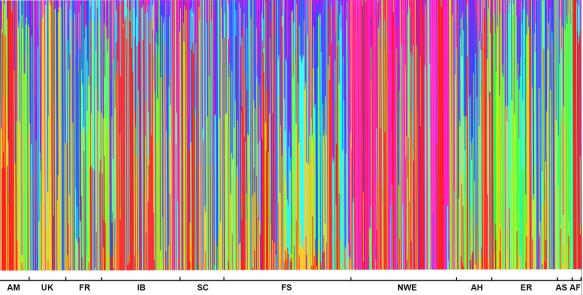

**Figure S3.** Analyses of population structure for the world-wide accessions used in this study (the 1001 Genomes). **a.** Population structure of under different assumptions about the number of clusters (K=2, 3, 4, 5, 6, 7). **b.** The cross-validation errors at various K values. **c.** Population structure analysis of worldwide 851 *A. thaliana* accessions (K = 8).

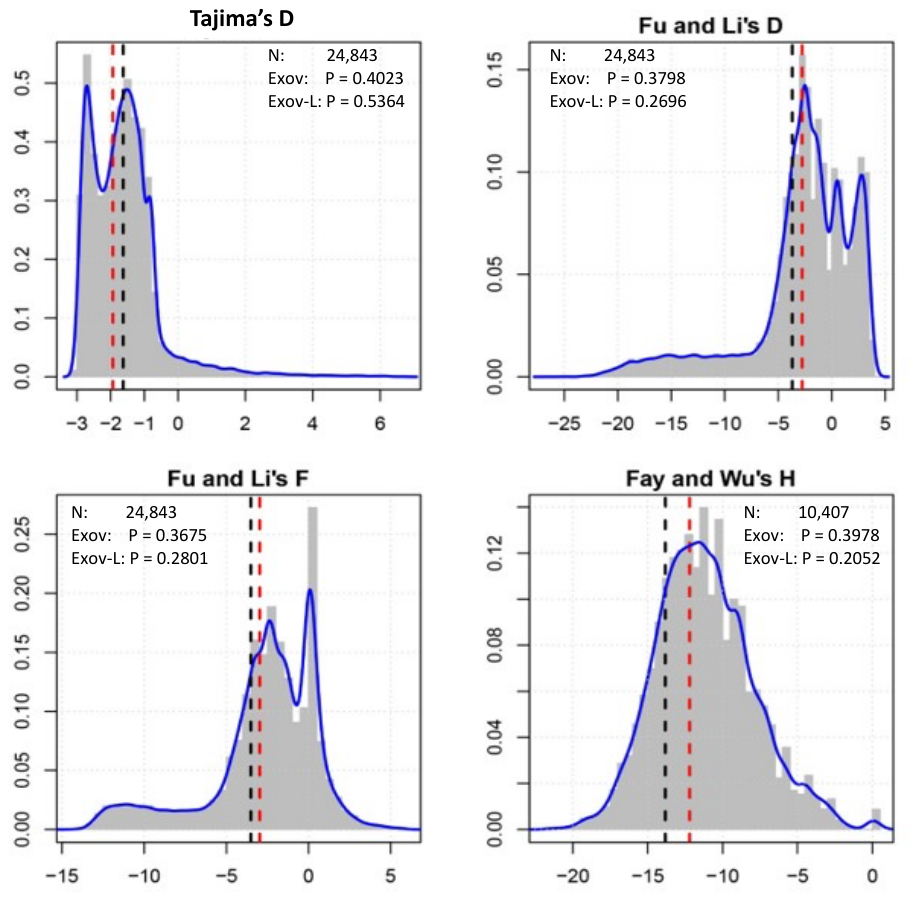

**Figure S4.** The empirical distributions of several population genetic test parameters across genome in A. thaliana and the probabilities of Exov and Exov-l in these distributions. N: the number of genes used in analysis. P is the probability equal to or lower than the observed parameters in *Exov* (red) or *Exov-l* (black).

**a.**

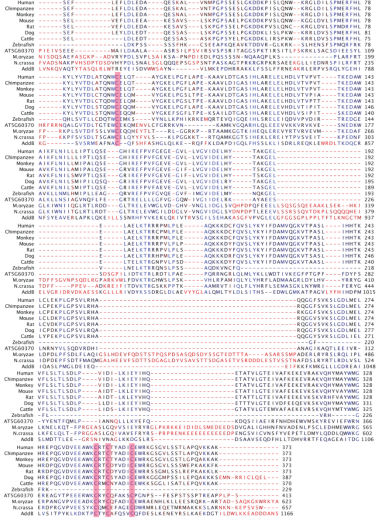

**b.**

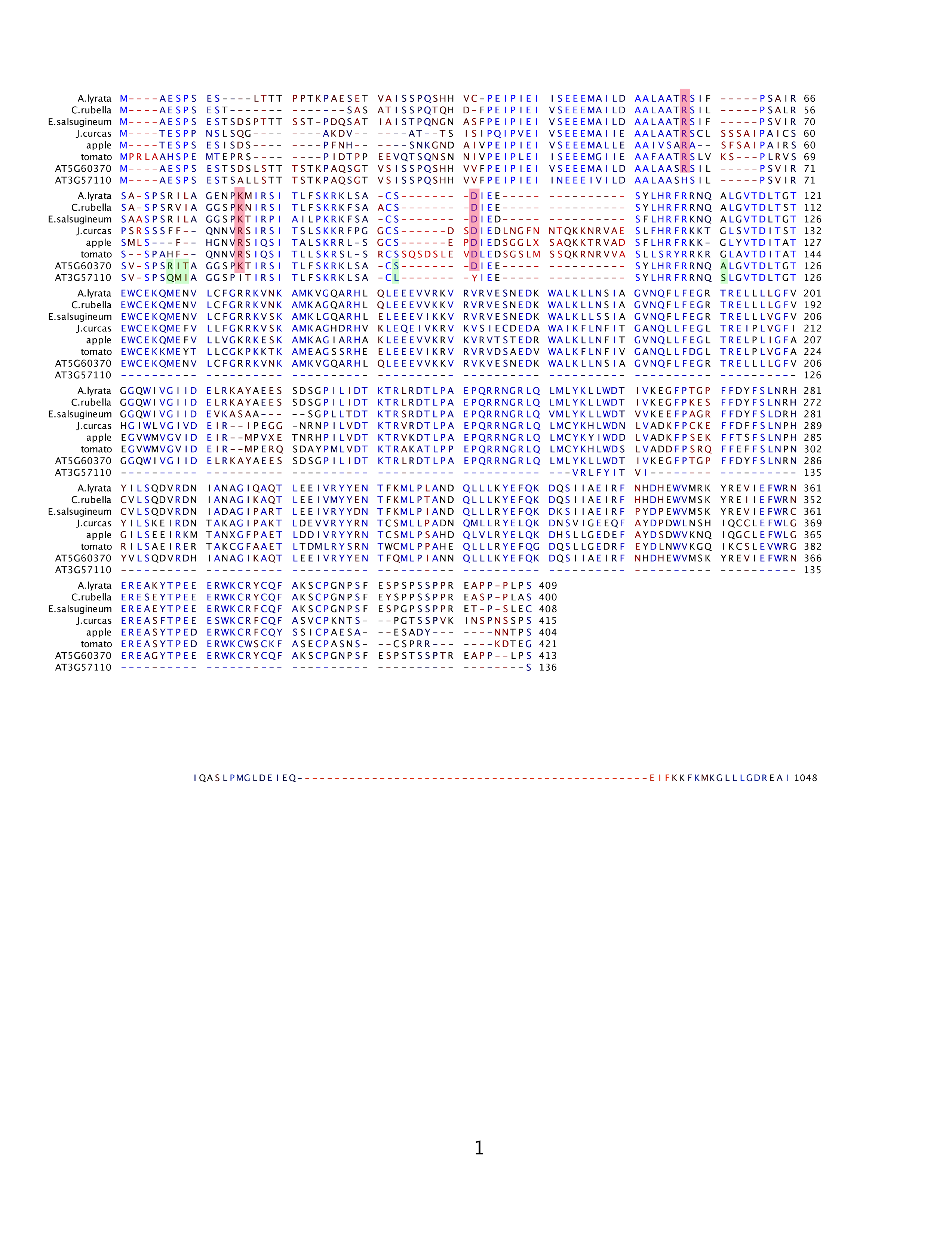

**c.**

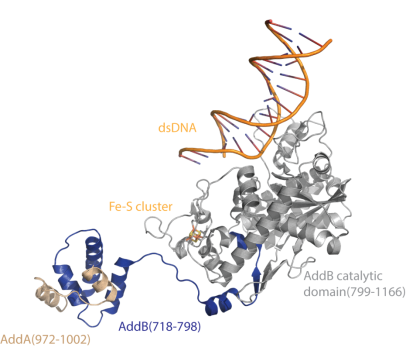

**Figure S5.** Protein sequence divergences of EXOV-L and EXOV. **a.** Alignment of EXOV-L (AT5G60370) with its homologs from _erent_ species and AddB (*B.subtilis*). These homologs are from Human (NP_073611.1), Chimpanzee (XP_003308065.1), Monkey (XP_001084006.1), Mouse (NP_001153515.1), Rat (NP_001101443.1), Dog (XP_532542.1), Cattle (NP_001075077.1), Zebrafish (NP_001032490.1), *M. oryzae* (XP_003718794.1), and *N. crassa* (XP_955908.1). The conserved Cysteine residues that coordinate the Fe-S cluster are highlighted in red. **b**. Alignment of EXOV (AT3G57110) and EXOV-L with its orthologs in the plant. The conserved polar residues at positions 63, 85, and 103 of AT5G60370 and its orthologs are highlighted in red. At position 63 of AT5G60370, the conserved residue is the basic polar residue arginine (R). In AT3G57110, this residue evolved to histidine (H). At position 85, the residue is either basic polar residue lysine (K) or arginine (R) in all instances except for that of AT3G57110, where it is substituted with the hydrophobic residue isoleucine (I). The conserved residue at position 103 is the acidic charged residue aspartate (D), which is changed to tyrosine (Y) in AT3G57110. Other residues such as R77, I78, T79, S102, and A119 were substituted with Q77, M78, I79, L102, and S119, highlighted in green. The NCBI accession number for the orthologs from *A. lyrata, C. rubella, E. salsugineum, J. curcas,* apple, and tomato are XP_002864682, XP_006280574, XP_006400854, KDP44101, XP_008358302, and XP_004251259. **c.** The proposed structural model of EXOV-L showing its conservation.

a.
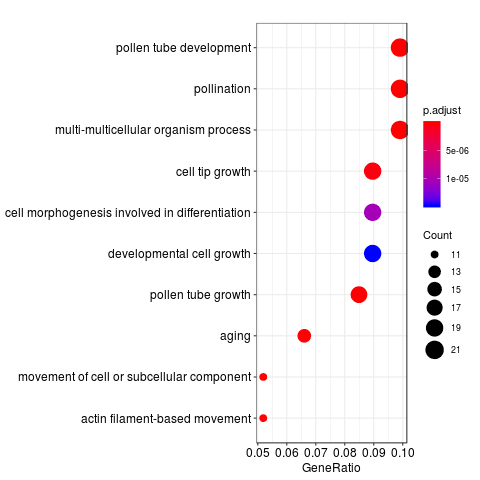

**b.**
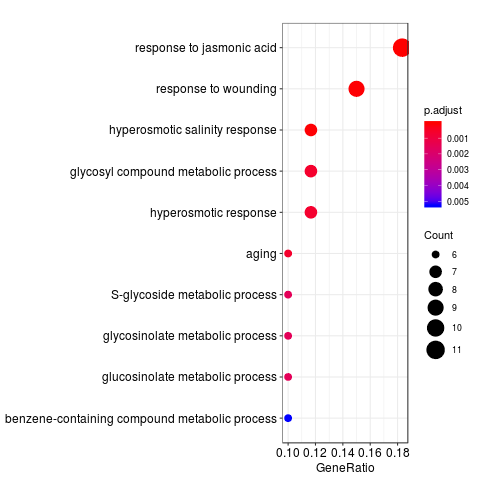

**Figure S6**. GO analyses. **a.** GO enrichment of analysis of the set of genes that significantly differentially expressed between wild type and *exov^crp^* . **b.** GO enrichment of analysis of the set of genes that significantly differentially expressed between wild type and *exov-l^crp^.*

**Figure S7.** Distribution of the phenotypic effects on seven traits of T-DNA mutants lines (single *exov, exov-l* and double *exov/exov-l*) and CRISPR/Cas9 mutant lines ( *exovcrp , exov-lcrp*) of the new gene and parental gene and wild type lines (Col-0). The curves are theoretical distributions modelled as Gaussian distribution.
