## Supplementary Figure S7 for "Species-specific gene duplication in *Arabidopsis thaliana* evolved novel phenotypic effects on morphological traits under strong positive selection"

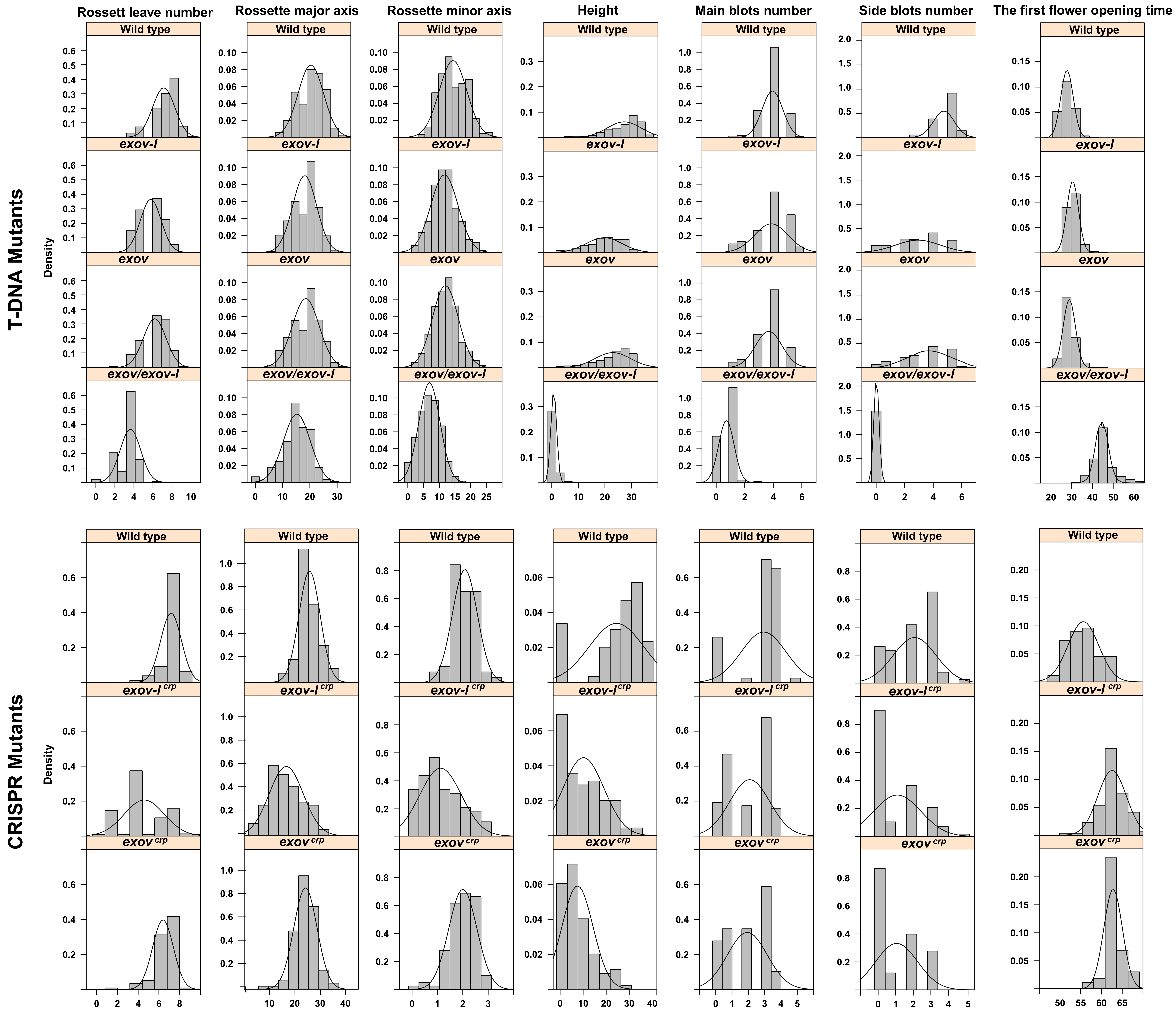


Figure S7 Distribution of the phenotypic effects on seven traits of T-DNA mutants lines (single *exov, exov-l* and double *exov/exov-l*) and CRISPR/Cas9 mutant lines ( $exov^{crp}$, $exov\text{-}l^{crp}$) of the new gene and parental gene. The curves were modeled as Gaussian distribution.
