## Supplementary file 1 for "Species-specific gene duplication in *Arabidopsis thaliana* evolved novel phenotypic effects on morphological traits under strong positive selection"

**CRISPR-P 2.0 : http://crispr.hzau.edu.cn/cgi-bin/CRISPR2/CRISPR**

57110 gRNA1：

TGTGGTGGTGCTAAGCAACG CGG

| Sequence | Off-score | MMs | Locus | Gene | Region |
| --- | --- | --- | --- | --- | --- |
| TGTGATGGTGCGGAGCAACCGGG | 0.121 | 4MMs | 5:-19295886 | AT5G47580 | exon |
| AGTGGTGGTGCAAAGTGACGAGG | 0.043 | 4MMs | 4:-4435296 | AT4G07650 | exon |
| TGTGGTGGTGCTAAGCGAGTCGG | 0.015 | 3MMs | 5:-24282892 | AT5G60370 | exon |

57110 gRNA2：

TCTTCATTGATGATCTCGAT TGG

| **Sequence** | Off-score | **MMs** | **Locus** | **Gene** | **Region** |
| --- | --- | --- | --- | --- | --- |
| ACTTCATCGACGATCTCGATCGG | 0.667 | 3MMs | 4:+2804545 | AT4G05520 | exon |
| TCTTCACTGACGATCTCGATTGG | 0.458 | 2MMs | 5:-24282990 | AT5G60370 | exon |
| CCTTGATTGATAATCTCCATTGG | 0.229 | 4MMs | 2:+13716797 | AT2G32290 | exon |
| GCTTCATGAATGATCTCCATCGG | 0.215 | 4MMs | 3:-357834 | AT3G02060 | exon |
| TCTTCATTGATAATCTAATTTGG | 0.162 | 4MMs | 3:+6226936 |  | Intergenic |
| TCTTGAATGATGAACTTGATGGG | 0.152 | 4MMs | 1:-26432964 | AT1G70200 | exon |
| TCATCCTTGGAGATCTCGATCGG | 0.135 | 4MMs | 5:+26012207 | AT5G65110 | exon |
| TCTCCATTGGTCATCTCCATTGG | 0.091 | 4MMs | 1:+21640530 |  | Intergenic |
| TCTTCATTGATAAAATCAATAGG | 0.089 | 4MMs | 1:+5604058 | AT1G16390 | exon |
| TCTTCTTTGGTGATCTTCATCGG | 0.088 | 4MMs | 3:-7904408 | AT3G22360 | exon |
| TCTTGATGGATGCTCACGATTGG | 0.084 | 4MMs | 5:-6080618 | AT5G18360 | exon |
| TTATCACTAATGATCTCGATTAG | 0.074 | 4MMs | 1:+7800235 |  | Intergenic |
| TCTTCTTCGATGATCTCTGTTAG | 0.046 | 4MMs | 3:-21528910 | AT3G58130 | exon |
| TCTTCTTCGATGATCTCTGTTAG | 0.046 | 4MMs | 2:-11698357 | AT2G27340 | exon |
| TCTTCATTTCTGATATTGATAGG | 0.018 | 4MMs | 2:+634172 | AT2G02410 | intron |
| TCTTCATTGATGTTTTCCATAGG | 0.010 | 3MMs | 3:+5016598 | AT3G14910 | intron |
| TTTACATTGATGATGTTGATTGG | 0.010 | 4MMs | 5:+1076795 | AT5G03990 | exon |
| TCTTCTTTAATTATGTCGATGGG | 0.009 | 4MMs | 2:-1822137 |  | Intergenic |
| TCTTCTTTAATTATGTCGATGGG | 0.009 | 4MMs | 4:-5331905 | AT4G08406 | CDS |
| TCTTCATTGATGAGCTGCTTGGG | 0.000 | 4MMs | 3:+17335888 | AT3G47060 | exon |

60370 gRNA1:

GGTGGTGCTAAGCGAGTCGG AGG

| **Sequence** | **Off-score** | **MMs** | **Locus** | **Gene** | **Region** |
| --- | --- | --- | --- | --- | --- |
| GAAGGTGCTAAGCGAGTCTTCGG | 0.282 | 4MMs | 4:+18048611 | AT4G38600 | exon |
| GGTGATGCTAACCGAGGTGGAGG | 0.035 | 4MMs | 2:-14039782 | AT2G33100 | exon |
| GGTGGTGCTAAGCAACGCGGAGG | 0.000 | 3MMs | 3:-21134955 | AT3G57110 | CDS |
| GGTGGTGCTAAACGACACGGAAG | 0.000 | 3MMs | 2:+1033750 | AT2G03410 | utr |

60370 gRNA2:

CCGCCGGCGGTTATCCGTGA CGG

| **Sequence** | **Off-score** | **MMs** | **Locus** | **Gene** | **Region** |
| --- | --- | --- | --- | --- | --- |
| CCGTCGGAGGTTATCAGTGGGGG | 0.398 | 4MMs | 5:+7136245 | AT5G21010 | exon |
| CCGCCGGCGATCATCTGTGACGG | 0.272 | 3MMs | 3:-21135152 | AT3G57110 | CDS |

Sample 60370-4

gRNA1, detected on-target, no off-target

T 5	24282887	. CG	C	1079.03 . AC=2;AF=1.00;AN=2;DP=28;ExcessHet=3.0103;FS=0.000;MLEAC=2;MLEAF=1.00;MQ=59.64;QD=32.57;SOR=0.693	GT:AD:DP:GQ:PL 1/1:0,28:28:81:1093,81,0

gRNA2, no on-target, no off-target

sample 57110-4

gRNA1, detected on-target, no off-target

T 3	21134956	. G	GT	912.03 . AC=2;AF=1.00;AN=2;BaseQRankSum=-1.339;DP=28;ExcessHet=3.0103;FS=0.000;MLEAC=2;MLEAF=1.00;MQ=59.40;MQRankSum=5.389;QD=32.57;ReadPosRankSum=0.743;SOR=0.804	GT:AD:DP:GQ:PL 1/1:1,27:28:49:926,49,0

gRNA2, no on-target, no off-target

Sample 57110-1-15

gRNA1, detected on-target, no off-target

T 3	21134956	. G	GT	910.03 . AC=2;AF=1.00;AN=2;BaseQRankSum=-0.527;DP=31;ExcessHet=3.0103;FS=3.332;MLEAC=2;MLEAF=1.00;MQ=60.00;MQRankSum=0.000;QD=32.50;ReadPosRankSum=1.241;SOR=0.568	GT:AD:DP:GQ:PL 1/1:1,27:28:49:924,49,0

gRNA2, no on-target, no off-target
