## Supplementary file 2 for "Species-specific gene duplication in *Arabidopsis thaliana* evolved novel phenotypic effects on morphological traits under strong positive selection"

<https://blast.ncbi.nlm.nih.gov/Blast.cgi?PAGE_TYPE=BlastSearch&SEARCH_INIT=ReprGenomeDBSearch&TAXID=3702>

Sequence-103969-32

CGTCATGTAGATACCTAACTTAGATGCGTCTCACTGGTGAAAAGAAAAACCACCCCAGTACATTAAAAACGTCCGCAATGTGTTATTTTACTCTGGTTTGAATTTTTTGCATAGACATGAAAAAAGAAGAAGACCACACGTTGGCTCCGAATAAAACATGGCCGAGTCACCCTCCGAGTCAACCTCCGCGTTGCTTAGCACCACCACATCCACTAAGCCCGCTCAATCCGGGACTGTTTCTATTTCAAGCCCTCAAAGTCACCATGTCGTCTTCCCTGAGATCCCAATCGAGATCATCAATGAAGAAGAAATAGTTATTCTCGATGCAGCTCTTGCAGCCTCCCACTCCATCCTACCTTCCGTCATTCGTTCTGTTTCTCCGTCACAGATGATCGCCGGCGGAAGCCCAATAACTATTCGTTCAATTACTCTTTTCTCCAAGAGGAAATTATCAGCTTGTTTATATATCGAAGAGTCTTATCTACATCGATTTAGAAGAAATCAGTCTTTAGGTGTCACAGATCTCACCGGTACTGTACGTTTGTTTTATATCACAGTCATATCATAGGAAGAAGAGATCGAATAGTAGCGACGACGTGGACAGCGGCTAATTGTGAGAGATGATTTAGATTTAGTACAATTGTTAGGTTTAATATCTCTTTTTCTGGTGAGATTAATTTGTGGTCCACTACTCATCCATTTCATTAATTTTCCATTTCATTGCTCATGTTATTTCATAATCTACAAACATATCTTTGGAATCTCACCATGATTGATAATTTCATTAATTCCAAAAAATTAGAACTAAAGTTTTATCATAGAATTTATAAAATTTGTTGTTTTGGGGTTTCCTTAATATTACCTAATTTTTTAAG

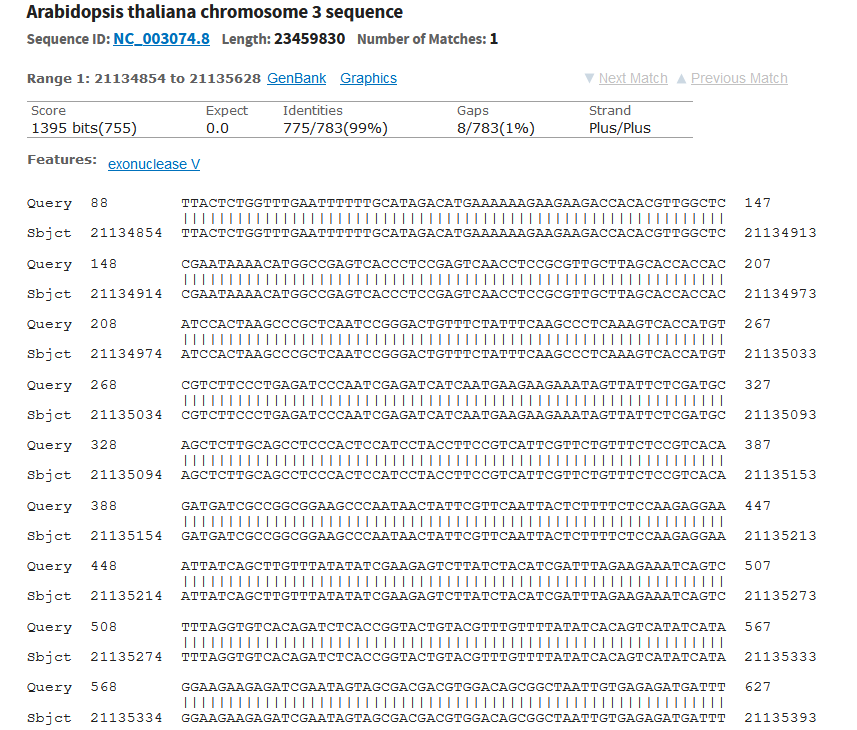

Sequence-103696-60

AGATACGTCTCACTGGTGAAGAAAACCACCCCAGTACATTAAAAACGTCCGCAATGTGTTATTTTACTCTGGTTTGAATTTTTTGCATAGACATGAAAAAAGAAGAAGACCACACGTTGGCTCCGAATAAAACATGGCCGAGTCACCCTCCGAGTCAACCTCCGCGTTGCTTAGCACCACCACATCCACTAAGCCCGCTCAATCCGGGACTGTTTCTATTTCAAGCCCTCAAAGTCACCATGTCGTCTTCCCTGAGATCCCAATCGAGATCATCAATGAAGAAGAAATAGTTATTCTCGATGCAGCTCTTGCAGCCTCCCACTCCATCCTACCTTCCGTCATTCGTTCTGTTTCTCCGTCACAGATGATCGCCGGCGGAAGCCCAATAACTATTCGTTCAATTACTCTTTTCTCCAAGAGGAAATTATCAGCTTGTTTATATATCGAAGAGTCTTATCTACATCGATTTAGAAGAAATCAGTCTTTAGGTGTCACAGATCTCACCGGTACTGTACGTTTGTTTTATATCACAGTCATATCATAGGAAGAAGAGATCGAATAGTAGCGACGACGTGGACAGCGGCTAATTGTGAGAGATGATTTAGATTTAGTACAATTGTTAGGTTTAATATCTCTTTTTCTGGTGAGATTAATTTGTGGTCCACTACTCATCCATTTCATTAATTTTCCATTTCATTGCTCATGTTATTTCATAATCTACAAACATATCTTTGGAATCTCACCATGATTGATAATTTCATTAATTCCAAAAATTAGAACTAAAGTTTTATCATAGAATTTATAAAATTTGTTGTTTGGGTTTCTTATATACTATTTTAGGGTTTAGTTTTCCTAATTAAAAACAACTAAAAATCTTATAGAAATATATAAATTGGAAAAGTTGTTGACTTTGAATCATTTGTGGAGAAATATCGCCTCTTCTATAGTCTGAAAATTTTAGCCGGCGATGTATACTAACTCATCAGTCATCACAAATAGTAACGACTTTATACAAATAGTATACTATTTGATAATGGAATGCTCTCTTCCTTGCGCATCTTCATTTATGT

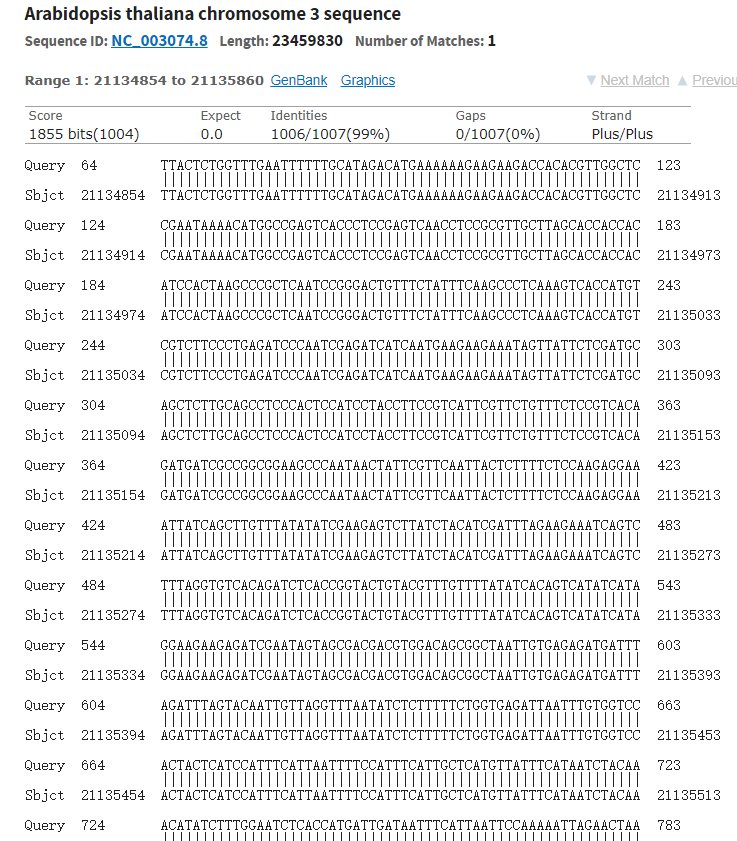

Seqence-101821

GACGAGTCTAATTGGTATTGTGATACGTCTCATGGTGAAAAAAAAACCCCCCAGTACATTAAAAACGTCCGCAATGTGTTATTAAGTTGTCTAAGCGTCAATTTGTTTACCCCACAATATATTGAGTATCCTCACCTGGAAATGGATCACGGAGCCCAGTCTTTGTATCTATCAATATTGGTCCGCTATTAGAAGACTCTTCTGCATACGCCTTTCTAATCTCGTCAATAATTCCCACAATCCATTGACCTCCAACAAATCCTAAGCTTTACAAGGTAAAATGCACCATATTTAAAGTCTCATTTGATGTATGCTACAAAACATTATCTTGCTAGGAAAAGGTACAAACCACGAAAATGTTACAATAGAAGATCGAACCTAGAGGGGTAAAAGGAACTCACAGTACCTAATCACGAGTTCGTCCTTCAAACAAAAACTGATTAACACCAAAAATGGAATTCAACAGCTTCAATGCCCACTTATCTTCATTTGATTCAACTTTTACTCTCACTTTTTTAACTACCTACCCAATTATATAAACACAACAACAAGTAGATTTCATCAAATTGTTTCACAAAACGATCACGTACTCTTAATAGTACCAATAAGCCCCTAAGATCCATCATAGTGAATAATACCTCTTCTTCAAGCTGCAAATGACGAGCCTGACCAACTTTTCATAGCTTTATTGACTTTTCTCCTGCCAAACAAAGAACATTCTCCATTTTGTTTCTGCACTACCCACCGCGTTGCCCTAATAGTTGAGTGCCTTAATTTACAAGAC

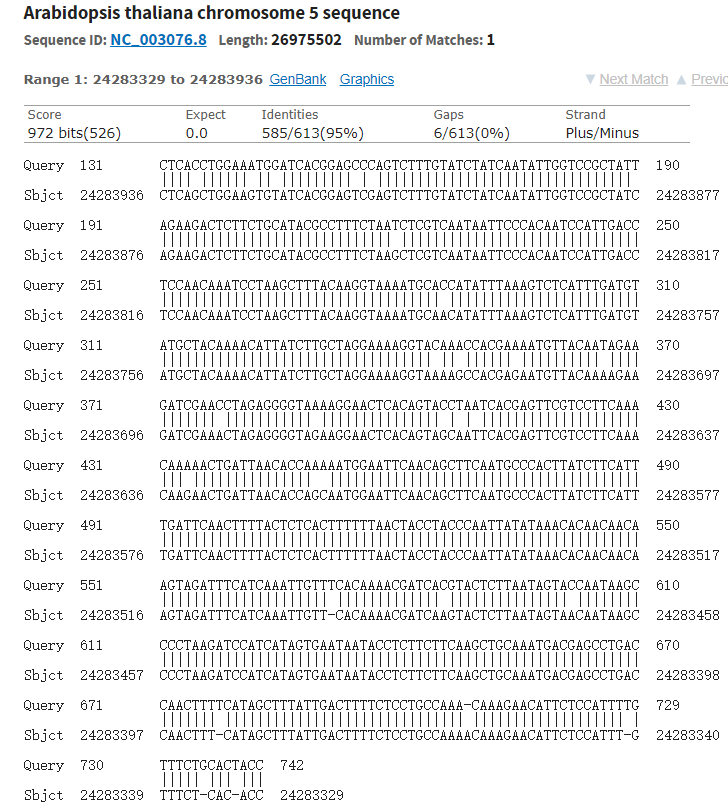
